# Parabrachial neuron types categorically encode thermoregulation variables during heat defense

**DOI:** 10.1101/2020.06.11.138370

**Authors:** Wen Z. Yang, Xiaosa Du, Wen Zhang, Cuicui Gao, Hengchang Xie, Yan Xiao, Xiaoning Jia, Jiashu Liu, Jianhui Xu, Xin Fu, Hongqing Tu, Xiaoyu Fu, Xinyan Ni, Miao He, Jiajun Yang, Hong Wang, Haitao Yang, Xiao-hong Xu, Wei L. Shen

## Abstract

Heat defense is crucial for survival and fitness, and its dysregulation may result in deaths due to poor management. Transmission of thermosensory signals into hypothalamic thermoregulation centers represent a key layer of regulation in heat defense. However, the mechanism by which these signals are transmitted into the hypothalamus remains poorly understood. Here, we reveal that glutamatergic prodynorphin and cholecystokinin neuron populations in the lateral parabrachial (LPB) are progressively recruited to defend elevated body temperature. These two nonoverlapping neuron types form circuitries with downstream preoptic hypothalamic neurons to inhibit BAT thermogenesis and activate tail vasodilation, respectively. Both circuitries are selectively activated by warm temperatures and are required for fever limiting. The prodynorphin circuitry is further required for regulation of energy expenditure and weight homeostasis. Thus, these findings establish that the genetic and functional specificity of heat defense neurons occurs as early as in the LPB and uncover categorical neuron types for encoding two heat defense variables, which may provide targets for treating thermoregulation disorders.

## INTRODUCTION

Homeostatic control of body temperature during heat defense is crucial for the survival and fitness in mammals. Heat defense-related disorders include heatstroke (*1, 2*) that causes hundreds of death each year in the US (*3*), and menopause thermal disequilibrium (MTD) (*4, 5*) that affects 75% of menopaused women (*5, 6*). Unfortunately, treatment is very limited against these disorders (*3, 7*), such as physical methods for heatstroke and hormone therapy for MTD, respectively. Heat defense is achieved by precise coordination between dedicated brain pathways and peripheral effector organs (*4, 8, 9*). Perturbation of these pathways may lead to temperature dysregulation (*4, 10*), and aggravate obesity and type 2 diabetes (*11, 12*). Previous findings have suggested that feed-forward temperature signals are detected by thermoreceptors expressed in dorsal root ganglion neurons (*13, 14*) and are transmitted into the spinal cord (*15*), which then are relayed by brain stem neurons the lateral parabrachial (LPB) (*16-18*) and reach the thermoregulation center the preoptic area (POA) (*4, 8, 9*). Several key types of POA neurons have been identified recently to control different aspects of heat-defense activities (*19-27*). For example, activation of neurons expressing the leptin receptor in the ventromedial preoptic area (VMPO) induces hypothermia along with a reduction of physical activities and energy expenditure (EE) (*23*). Neurons expressing brain-derived neurotrophic factor/pituitary adenylate cyclase-activating polypeptide (BDNF/PACAP) in the anterior ventromedial preoptic area (VMPO) induce hypothermia, reduce thermogenesis by interscapular brown fat tissues (iBATs), impair nest building in cold temperature, and increase tail vasodilation (*21, 24*). GABAergic neurons in the ventral part of the lateral preoptic nucleus (vLPO) also induce hypothermia and lower BAT activities (*21, 28*), while its inhibition causes fever-level hyperthermia (*21*). Activation of glutamatergic NOS1^+^ or galanin neurons in the ventrolateral preoptic area promotes sleep and hypothermia (*19, 20*). These together demonstrate the POA as the thermoregulatory center to differentially regulate diverse thermoregulation variables. However, the circuitries that provide key inputs to the POA to coordinately control these variables, remain poorly understood.

The LPB is a complex structure that regulates a series of physiological activities, including thermoregulation (*29-32*), pain and itch (*33-35*), sodium (*36*) and food intake (*37-40*), blood glucose (*41, 42*), and wakefulness (*43, 44*). Prior pioneering work suggests that the LPB glutamatergically transmits key cutaneous warm and cold sensory signals into the POA via two different subareas, the LPBd (dorsal) and the LPBel (external lateral) respectively (*16, 18, 45*). Pharmacological activation of neurons in the LPBd and the LPBel elicit autonomic responses to warm and cold exposures, respectively (*16, 18*). Electrophysiological studies suggest a direct connection between LPB neurons and POA neurons (*16, 18*). Besides, lesioning or silencing in the LPB abolishes behavioral thermal preferences (*29*). Noticeably, most cold-activated neurons in the LPBel are FoxP2^+^, while half of the warm-activated neurons in the LPBd are prodynorphin (Pdyn) neurons that coexpress a glutamatergic marker and project to the POA (*31*). Chemogenetic activation of these Pdyn neurons induces hypothermia (*46*). Although strongly implied by previous studies, it is unclear whether and how LPB Vglut2^+^ or Pdyn^+^ neurons are functionally connected with POA neurons as it might affect other thermoregulatory neurons such as the recently identified dorsal raphe nucleus neurons (*47*). Further, whether the LPB just plays a relay role by feeding-forward warm sensory signals or it is also involved in sorting these signals to control different thermoregulation variables specifically, is unknown. Decoding this specificity is vital for drug design targeting heat defense-related disorders.

Further, heat defense is a context-dependent process with timely adjustment of thermoregulation variables according to different ambient temperatures, yet the mechanism is still unclear. Therefore, we combined functional manipulations with projection-specific transcriptome analysis and activity recordings to define key elements of the LPB→POA circuitry during heat defense and establish the LPB is a critical thermoregulation center to regulate heat defense variables differentially. LPB Pdyn neurons and cholecystokinin (CCK) neurons are progressively recruited in response to increasing ambient temperatures. The two neuron types categorically regulate BAT thermogenesis and tail vasodilation, respectively, and differentially regulate muscle shivering. Neural blocking further suggests these neurons are required to limit fever development. These data suggest the thermoeffector specificity exists as early in the cutaneous afferent thermoregulatory reflex pathway as the LPB, and identify two potential target neurons for treating thermoregulation disorders, including heatstroke (*1, 2*) and menopause thermal disequilibrium (*4, 48*). Additionally, blocking LPB Pdyn neurons prevents weight gain driven by high-fat diet, suggesting a strong interconnection between body temperature and weight homeostatic systems.

## RESULTS

### Graded changes in iBAT thermogenesis and tail vasodilation during heat defenses

To document changes of thermoregulation variables during heat defense, we monitored the core temperature (T_core_, measured intraperitoneally) together with the surface temperature of iBATs (T_iBAT_) and tail skin (T_tail_) during exposures to different warm ambient temperatures (T_a_). As expected, these variables changed quickly after switching T_a_ from RT (25 °C) to warm temperatures (Fig. 1a-d). To quantify the level of iBAT thermogenesis, we substracted T_core_ from T_iBAT_ and found a graded inhibition to increasing T_a_ levels (Fig. 1e-f). We observed a complete inhibition of iBAT thermogenesis after 38 °C exposure, where T_iBAT_ dropped to the T_core_ level (Fig. 1f). Furthermore, to quantify the level of vasodilation, we substracted T_a_ from T_tail_ and found a graded activation to increasing T_a_ levels (Fig. 1g-h). Similarly, the level of tail vasodilation reached a maximum after 38 °C exposure, where T_tail_ dropped to the T_a_ level (Fig. 1h). Together, we reveal the dynamic changes of two variables during heat defense and suggest inhibition of iBAT thermogenesis might be also an important variable in addition to vasodilation.

**Figure 1.**
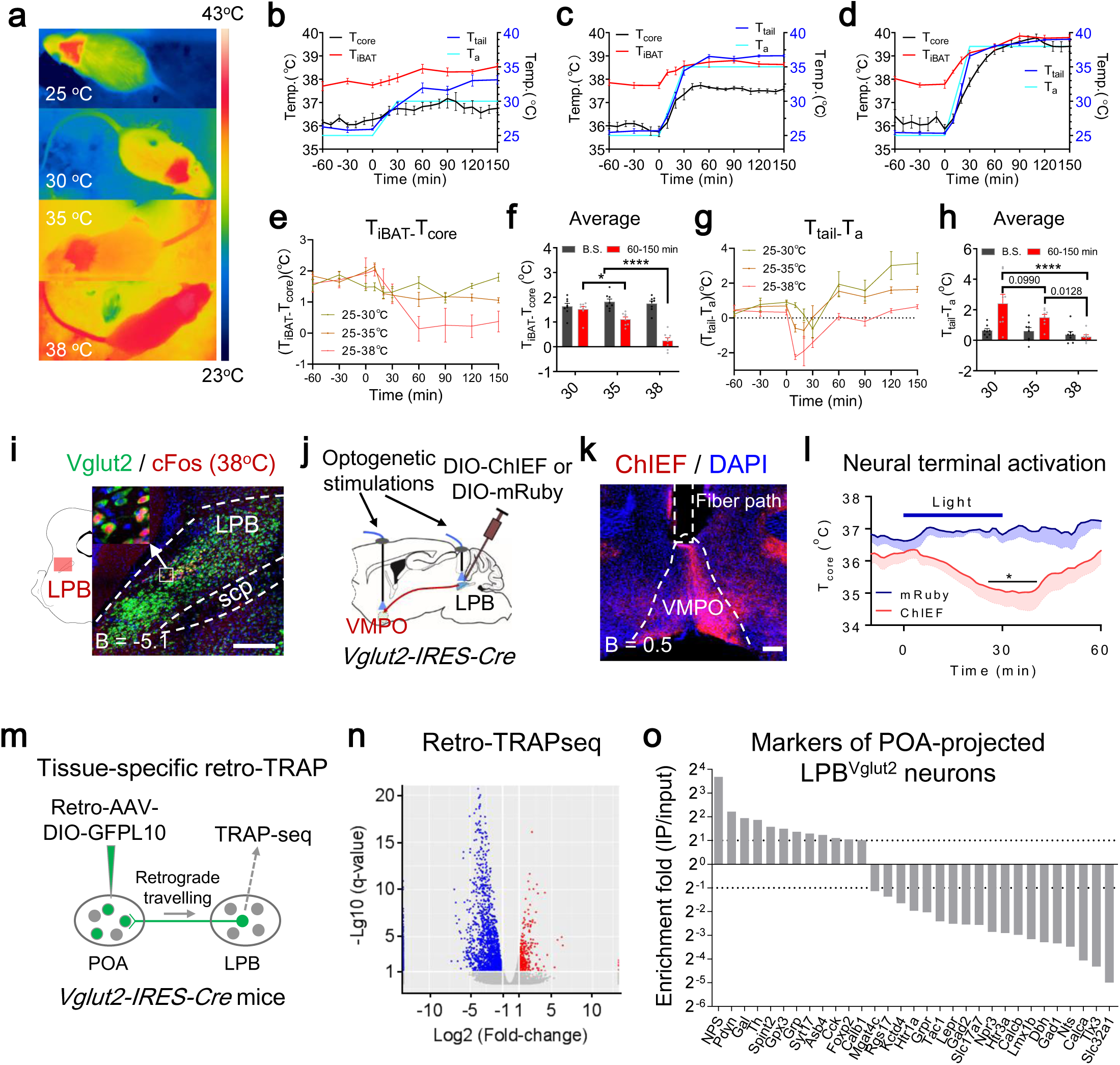
Projection-specific functional and transcriptome analysis of the LPB→POA pathway. **a-d**, Changes of T_core_, T_iBAT_, and T_tail_ under different ambient temperatures (T_a_) (n = 8 each). The coat hair on top of the iBATs was shaved. **e-f**, Changes of T_iBAT_ ^_^ T_core_ under different ambient temperatures (T_a_) (n = 8 each). **g-h**, Changes of T_tail_ ^_^ T_a_ under different ambient temperatures (T_a_) (n = 8 each). **i**, The overlapping between warm-induced cFos and Vglut2-IRES-Cre & LSL-GFPL10 in the LPB. **j**, Scheme of optogenetic activation of glutamatergic (Vglut2) LPB soma and terminals in the VMPO. **k**, Terminal expression of ChIEF from LPB^Vglut2^ neurons in the VMPO. **l**, Changes of T_core_ after photoactivation LPB^Vglut2^ terminal in the VMPO (n = 4 each). Laser pattern: 473 nm, 6 mW, 20 Hz, 10 ms, 2-s on 2-s off, 30 min. **m**, Scheme of tissue-specific retro-TRAP, where translational ribosomes from VMPO-projected LPB^Vglut2^ neurons were immunoprecipitated and associated mRNAs were used for sequencing. **n**, Volcano plots (q-value vs log_2_ fold change) for LPB mRNAs after retroTRAP sequencing. Logarithmic ratios of mRNA enrichment fold (IP/Input, n = 3) plotted against the q-value (where q ⩽ 0.5 is considered significant) of the hierarchical linear model. Positive fold changes indicate an enrichment, and negative fold changes indicate a depletion in precipitated mRNAs. **o**, Enrichment fold (IP/Input) of PB-expressed genes from Allen Institute. Scale bars, 200 µm. All data are shown as mean ± s.e.m. The p-values are calculated based on statistical tests in Supplemental Table 1. *p ⩽ 0.05; ****p ⩽ 0.0001. B, bregma; LPB, lateral parabrachial nucleus; scp, superior cerebellar peduncle; VMPO, ventromedial preoptic nucleus.

### Critical roles of glutamatergic LPB→POA projection in reducing body temperature

To find the neural substrates encoding different heat defense variables, we considered the LPB, an early player in the thermoregulatory reflex pathway. The LPB contains mostly glutamatergic neurons (*16, 31*). Both warm and cold-induced cFos in the LPB colocalized predominantly with glutamatergic marker Vglut2 (∼95% for both; Vglut2 stands for Vesicular Glutamate Transporter 2, encoded by *Slc17a6*; Fig. 1i and Fig. S1). To directly test their roles in thermoregulation, we used optogenetics to activate these neurons by targeted expression ChIEF, a current-stabilized channelrhodopsin (*49*), into the LPB of Vglut2-IRES-Cre mice. Photoactivation of these neurons led to unstable phenotypes, which induced hypothermia or hyperthermia after stimulating with different laser frequencies (Fig. S2a-c), which is consistent with the existence of thermoregulatory neurons to both warm and cold temperatures (*16, 18, 31*).

Next, we suspected that projection-specific functional manipulation might be able to separate the LPB function in warm- and cold-induced thermoregulation. Thus, we sought to verify the function of LPB glutamatergic projections to the POA in thermoregulation as proposed before (*4, 8, 9*). Indeed, photoactivation of the LPB^Vglut2^ terminals in the VMPO caused severe hypothermia scaled with laser frequencies (Fig. 1j-l and Fig. S2d).

Furthermore, to consolidate that POA neurons are the postsynaptic targets of LPB neurons, we activated VMPO neurons innervated by LPB neurons by using an anterograde transsynaptic tracer AAV1-Cre (*50*) injected into the LPB. This tracer induced expression of DIO-hM3D_q_ injected into the VMPO and these VMPO^hM3Dq^ neurons were mainly glutamatergic as verified by glutamate staining (Fig. S2e-g). Chemogenetic activation of these VMPO neurons reduced T_core_, EE, and physical activity (Fig. S2h-k), further supporting the role of this connection in thermoregulation. Thus, our data suggest the glutamatergic LPB→VMPO pathway is critical for body cooling.

### Transcriptome profiling of the LPB^Vglut2^→POA projection

The finding that the LPB^Vglut2^→VMPO projection is critical for thermoregulation prompts whether we could identify specific genetic markers in this pathway. To do this, we adopted a cell-type and projection specific translational ribosomal affinity purification technique (tissue-specific retro-TRAP) based on a retrograde AAV (*51*). We injected a retro-AAV carrying DIO-GFP-L10 transgene (L10 is a ribosomal protein.) (*52*) into the VMPO, which expressed GFP-L10 in the LPB of the Vglut2-IRES-Cre mice (Fig. 1m). We confirmed that warm-induced cFos colocalized with the GFP-L10 fluorescence in the LPB (Fig. S3a-d). The GFP-labeled ribosomes were immunoprecipitated by anti-GFP antibodies after the LPB microdissection, and the associated mRNAs were used for sequencing. We analyzed the enrichment fold (IP/input) and found that 364 genes were significantly upregulated and 1958 genes were downregulated (Fig. 1n). Then, we focused on the enrichment fold of PB-expressed genes downloaded from Allen Institute (https://alleninstitute.org/), and found upregulated genes included *neuropeptide S* (*nps*), *prodynorphin* (*Pdyn*), *galanin* (*gal*), *tyrosine hydroxylase* (*th*), *cholecystokinin* (*CCK*) and *FoxP2*. Top downregulated genes included the GABAergic markers *Gad1* and *Slc32a1*, and *LepR* (Fig. 1o; Fig. S3e). We chose these markers for further studies.

### Function of LPB neurons in heat defense

We then analyzed the co-labeling between temperature-induced cFos and genetic markers found in retro-TRAP. NPS was broadly expressed in the LPB, which colocalized with cFos induced by both heat exposure (∼90%) and cold exposure (∼88%) without an apparent selectivity (Fig. S4). Consistent with prior studies (*31*), Pdyn-IRES-Cre colocalized with Pdyn (∼87%) and glutamate staining (∼92%) (Fig. 2a-b; Fig. S5a-b), and labeled ∼45% of cFos induced by heat exposure and only ∼20% of cFos induced by cold exposure (Fig. 2c-d; Fig. S5c-d). Interestingly, Pdyn neurons were increasingly recruited during heat exposures as the increases of T_a_, which finally maintained a constant overlapping ratio over cFos regardless of exposure temperatures (Fig. 2e). Unexpectedly, there were very few Gal-Cre^+^ cells in the LPB. Instead, we found a cluster of Gal-Cre^+^ neurons in a nearby area, the laterodorsal tegmental nucleus (LDTg) (Fig. S6a), which presumably was due to tissue contamination during dissection. For TH, dense neural terminals and few cell bodies were observed in the LPB (Fig. S6b), suggesting the enriched TH might be from neural terminals.

**Figure 2.**
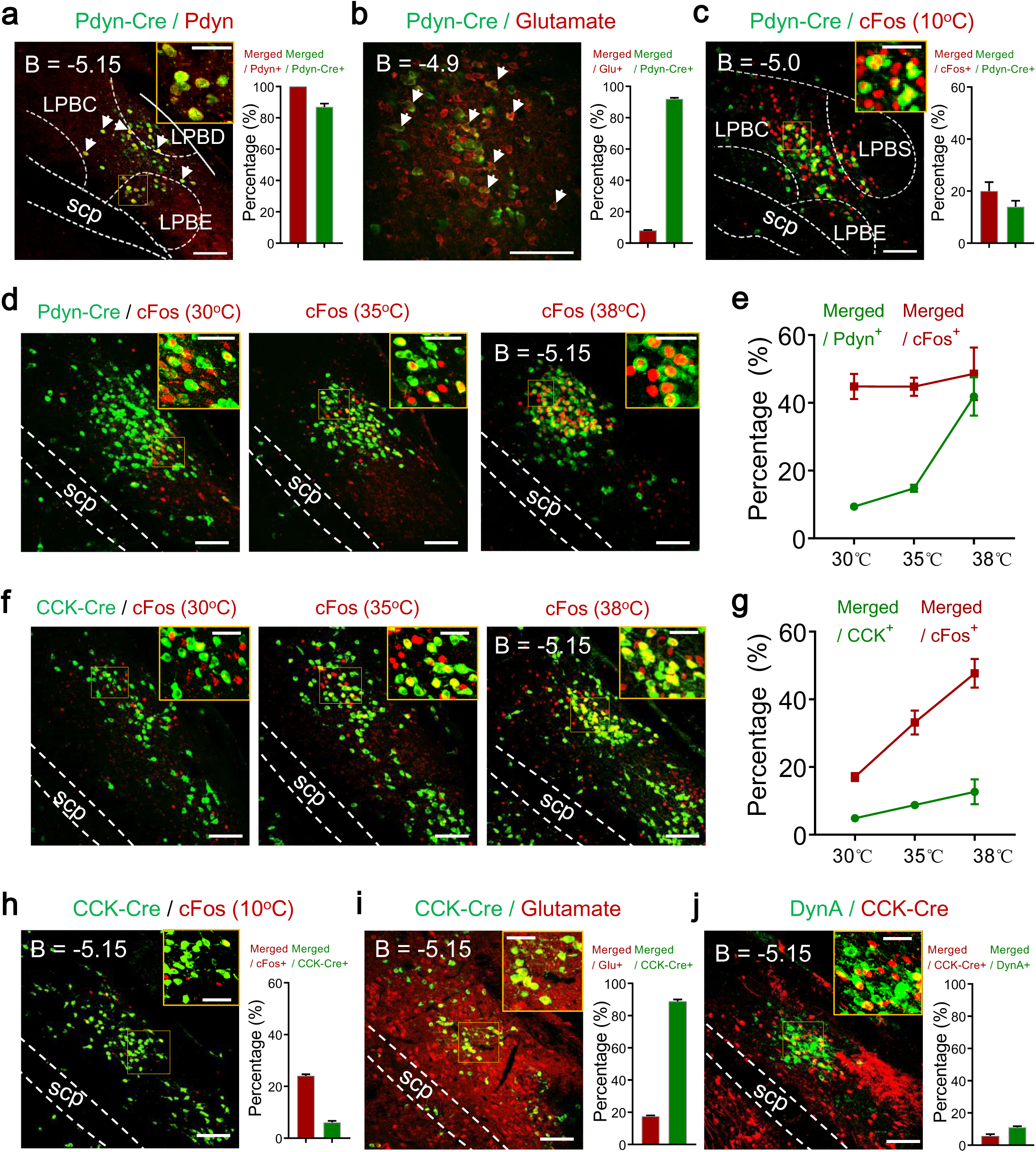
Progressive recruitment of LPB^Pdyn/CCK^ neurons in heat defense. **a**, The overlapping between Pdyn-IRES-Cre & AAV-DIO-GFP and Pdyn staining in the LPB (n = 3 each). Arrows, examples of merged cells. **b**, The overlapping between Pdyn-Cre & LSL-GFPL10 and glutamate staining in the LPB (n = 3 each). **c**, The overlapping between Pdyn-Cre & LSL-GFPL10 and cold-induced cFos (10 °C) in the LPB (n = 3 each). **d-e**, The overlapping between Pdyn-Cre & LSL-GFPL10 and cFos induced by different warm ambient temperatures in the LPB (**d**; n = 3 each), and the quantification of the overlapping rates (**e**). Ambient temperatures were indicated in the figure. **f-g**, The overlapping between CCK-IRES-Cre & LSL-GFPL10 and cFos induced by different warm ambient temperatures in the LPB (**f**; n = 3 each), and the quantification of the overlapping rates (**g**). Ambient temperatures were indicated in the figure. **h**, The overlapping between CCK-Cre & LSL-GFPL10 and cold-induced cFos (10 °C) in the LPB (n = 3 each). **i**, The overlapping between CCK-Cre & LSL-GFPL10 and glutamate staining in the LPB (n = 3 each). **j**, The overlapping between CCK-Cre & LSL-GFPL10 and DynA staining in the LPB (n = 3 each). Scale bars, 100 µm except in upper right boxes (50 µm). All data are shown as mean ± s.e.m. B, bregma; scp, superior cerebellar peduncle; LPBC, lateral parabrachial nucleus, central part; LPBD, lateral parabrachial nucleus, dorsal part; LPBS, lateral parabrachial nucleus, superior part; LPBE, lateral parabrachial nucleus, external part.

Interestingly, CCK-IRES-Cre^+^ neurons, which have been shown to regulate blood glucose (*41, 42*), showed increasing overlapping ratios (up to 48%) over cFos induced by increasing warm temperatures (Fig. 2f-g; Fig. S7a). Similar to Pdyn neurons, more CCK neurons were recruited when the T_a_ was increased, but the overlapping ratio over cFos also kept increasing (Fig. 2j). The overlapping ratio over cFos induced by cold exposure was only ∼20% (Fig. 2h; Fig. S7b). Also, most CCK neurons were glutamatergic as suggested by glutamate immunostaining (Fig. 2i; Fig. S7c). We also checked the overlapping between Dynorphin A (DynA) and CCK and found little overlapping (Fig. 2j; Fig. S7d). Nevertheless, we also analyzed the overlapping between temperature-induced cFos and Gad1^+^ cells and found few Gad1^+^ cells in the LPB, which showed nearly no overlapping with cFos (Fig. S6c-d).

To address whether these LPB neurons are essential for thermoregulation, we used DREADD-hM3D_q_ to activate them. Consistent with cFos pattern, activation of LPB^Pdyn^ neurons reduced T_core_ (Fig. S8a-c; also see (*46*)). It also lowered physical activity and EE (Fig. S8d-f). The reduction of T_core_ and EE were reversed by injection of adrenergic β3 agonist CL316243, supporting that sympathetic nerves might be the downstream targets (Fig. S8g-h). It did not affect food intake in fasted animals (Fig. S8i). Similarly, DREADD activation of LPB^CCK^ neurons reduced T_core_, EE and physical activities, but not food intake (Fig. S9a-f). It slightly yet insignificantly reduced blood glucose level (Fig. S9g), suggesting this is a separate CCK^+^ population from the reported glucose upregulation neurons (*41, 42*). Nevertheless, we tested the role of LPB GABAergic neurons labeled by Vgat-IRES-Cre (Vgat stands for vesicular GABA transporter encoded by *Slc32a1*.) and LepR-Cre. DREADD activation of Vgat neurons did not change T_core_ significantly (Fig. S9h-j). Activation of LepR-Cre neurons slightly increased T_core_ and insignificantly affected EE and physical activity (Fig. S9k-o). Together, we identified two types of neurons, Pdyn and CCK neurons, as heat defense neurons in the LPB.

### Neural dynamics of LPB^Pdyn/CCK^→POA circuitries to temperature changes

Inspired by the function LPB Pdyn and CCK neurons in heat defense, we sought to test the sensitivity of LPB^Pdyn/CCK^→POA circuitries to temperatures. Therefore, we performed fiber photometry to record the calcium dynamics using GCaMP6s (Fig. 3a-b). As expected, we observed a sustained calcium change to warming (Fig. 3c). In contrast, the cooling-induced calcium changes were smaller and lasted shorter than that of warming (Fig. 3d). These results are consistent with cFos patterns where the merge ratio of Pdyn and warm-induced cFos was twice of cold-induced (Fig. 2c-e). Then, we measured calcium activities to different floor temperatures and found the responses were scaled with changes of floor temperatures and nociceptive temperatures (45°C or 5°C) induced a much larger response than warm or cold temperatures (Fig. 3e-f; Fig. S10a-b). Also, we measured responses to capsaicin and menthol that elicit hot and cool sensation, respectively (*53*). Surprisingly, we did not detect any significant responses to these chemicals for Pdyn neurons and even for Vglut2 neurons in the LPB (Fig. S10c-h; Movie S1). The high baseline fluctuations in LPB^Vglut2^ neurons might be due to the pain during drug injection.

**Figure 3.**
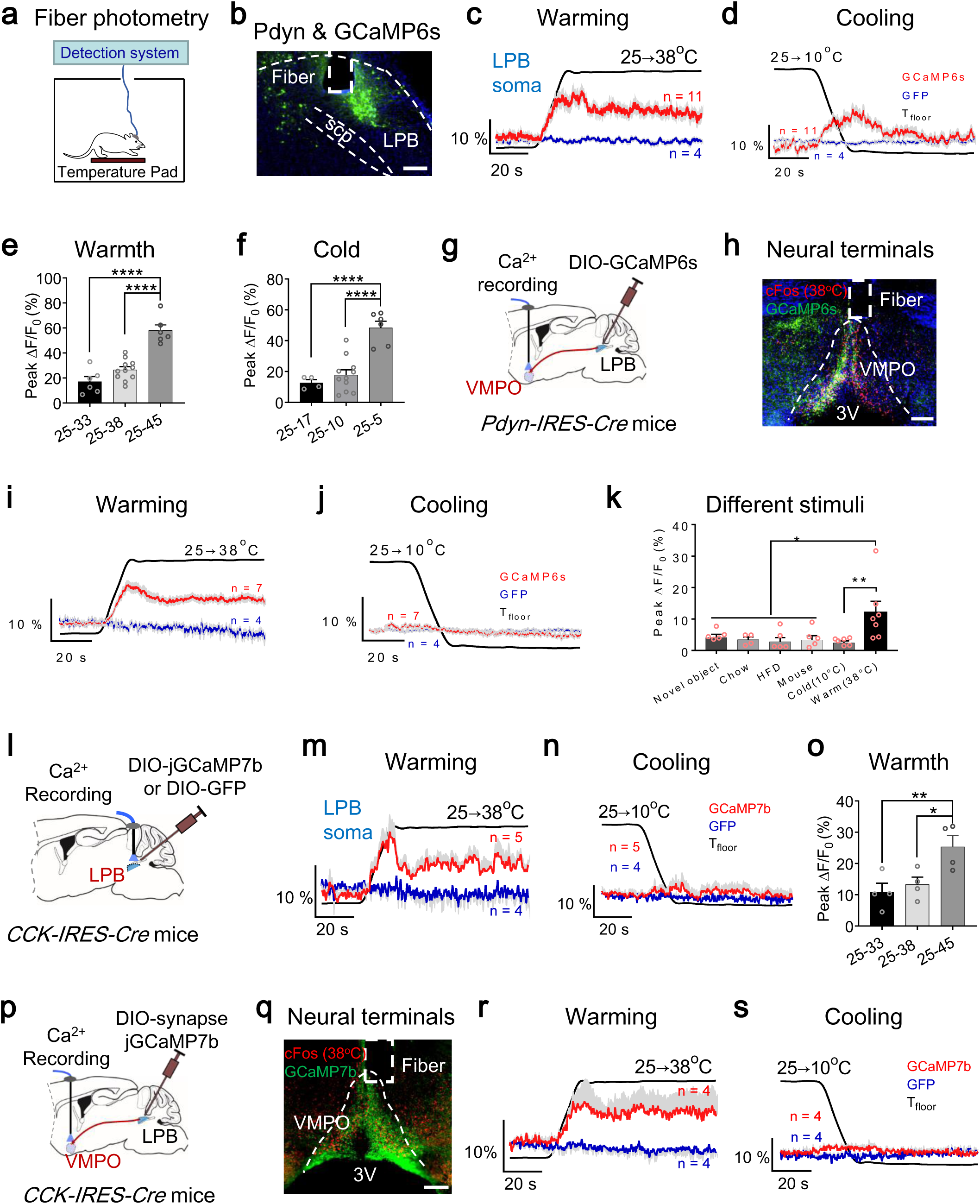
LPB^Pdyn/CCK^→POA circuitries are sensitive to warm temperatures. **a**, Scheme of fiber photometry. The floor temperature (T_floor_) was controlled by a Peltier device. **b**, Viral expression of DIO-GCaMP6s in LPB^Pdyn^ neurons. **c-d**, Calcium dynamics after warming (**c**) or cooling (**d**) of the floor respectively. ΔF/F_0_ represents the change in GCaMP6s fluorescence from the mean level (t = (– 120 – 0 s)) (as to **i, j, m, n, r, s**). The GFP was used as a control. **e-f**, Peak responses of Pdyn neuron to different warm (**e**) or cold (**f**) temperatures (25 – 33 °C, 25 – 45 °C, 25 – 5 °C, n = 6 each; 25 – 38 °C, 25 – 10 °C, n = 11 each; 25 – 17 °C, n = 4). **g**, Scheme for calcium recording of LPB^Pdyn^ terminal signals in the VMPO. **h**, Terminal expression DIO-GCaMP6s from LPB^Pdyn^ neurons and warm-induced cFos. **i-j**, Terminal calcium dynamics after warming (**i**) or cooling (**j**), respectively. **k**, Summary of terminal neural responses to the indicated stimuli, including novel objects, chow food, high-fat food, mouse of the same sex, and temperatures (Cold, Warm, n = 7 each; Novel object, HFD, Mouse, n = 6 each; Chow, n = 4). **l**, Scheme for soma recording of calcium signals in LPB^CCK^ neurons. **m-n**, Calcium dynamics after warming (**m**) or cooling (**n**) of the floor, respectively. **o**, Peak responses of LPB^CCK^ neurons to different warm temperatures (n = 4 each). **p**, Scheme for calcium recording of LPB^CCK^ terminal signals in the VMPO. **q**, Terminal expression DIO-synapse-jGCaMP7b from LPB^CCK^ neurons and warm-induced cFos. **r-s**, Calcium dynamics after warming (**r**) or cooling (**s**) of the floor, respectively. Scale bars, 200 µm. All data are shown as mean ± s.e.m. The p-values are calculated based on statistical tests in Supplemental Table 1. *p ⩽ 0.05; **p ⩽ 0.01; ****p ⩽ 0.0001; 3V, 3rd ventricle.

We suspected that the terminal-specific recording of LPB^Pdyn^ activity in the POA might be able to capture warm-selective responses. Therefore, we measured terminal calcium activity in the VMPO (Fig. 3g-h) and also found a sustained response to warming (Fig. 3i). In contrast to the soma, we did not detect any significant changes to cooling (Fig. 3j), suggesting the activity of this projection is warm specific. Consistent with the calcium activity, we found that LPB^Pdyn^ neural terminals in many subregions of the POA, including the VMPO and the LPO, closely associated with warm-induced cFos (Fig. S10i). Also, we measured responses to other stimuli, including the novel object, chow, HFD, capsaicin, menthol, icilin and mouse, and found no significant changes to these stimuli (Fig. 3k and Fig. S10j-k).

With similar approaches, we found that LPB^CCK^ soma and their terminals in the VMPO responded selectively to warming over cooling (Fig. 3l-s). Similar to Pdyn circuitry, the CCK circuitry was not responsive to capsaicin, menthol, and icilin (Fig. S10l). Together, we showed that LPB^Pdyn/CCK^ →POA circuitries are selectively sensitive to warming and the responses are increased as the increase of floor temperatures.

### Function of LPB^Pdyn/CCK^→POA circuitries in reducing body temperature

The selective sensitivity of LPB^Pdyn/CCK^→POA circuitries to warmth predicts these circuitries might be relevant in heat defense. To test this, we first verified the synaptic connections with POA neurons by recording the light-induced excitatory postsynaptic currents (EPSCs) from VMPO neurons in LPB^Pdyn & ChIEF^ mice. These currents were blocked by glutamate receptor antagonists CNQX/AP5 (Fig. 4a-b), further supporting a glutamatergic transmission mechanism. The latency after photoactivation was within the range of monosynaptic connection (*54*). To further verify this monosynaptic connection, we adopted a method used before (*55*). We first blocked the EPSCs with tetrodotoxin (TTX) and then applied 4-Aminopyridine (4-AP) to sensitize the postsynaptic current. We found the EPSCs recovered after 4-AP treatment, suggesting a monosynaptic connection (Fig. 4c). Similarly, LPB CCK neurons also formed monosynaptic connections with VMPO neurons (Fig. 4d-f).

**Figure 4.**
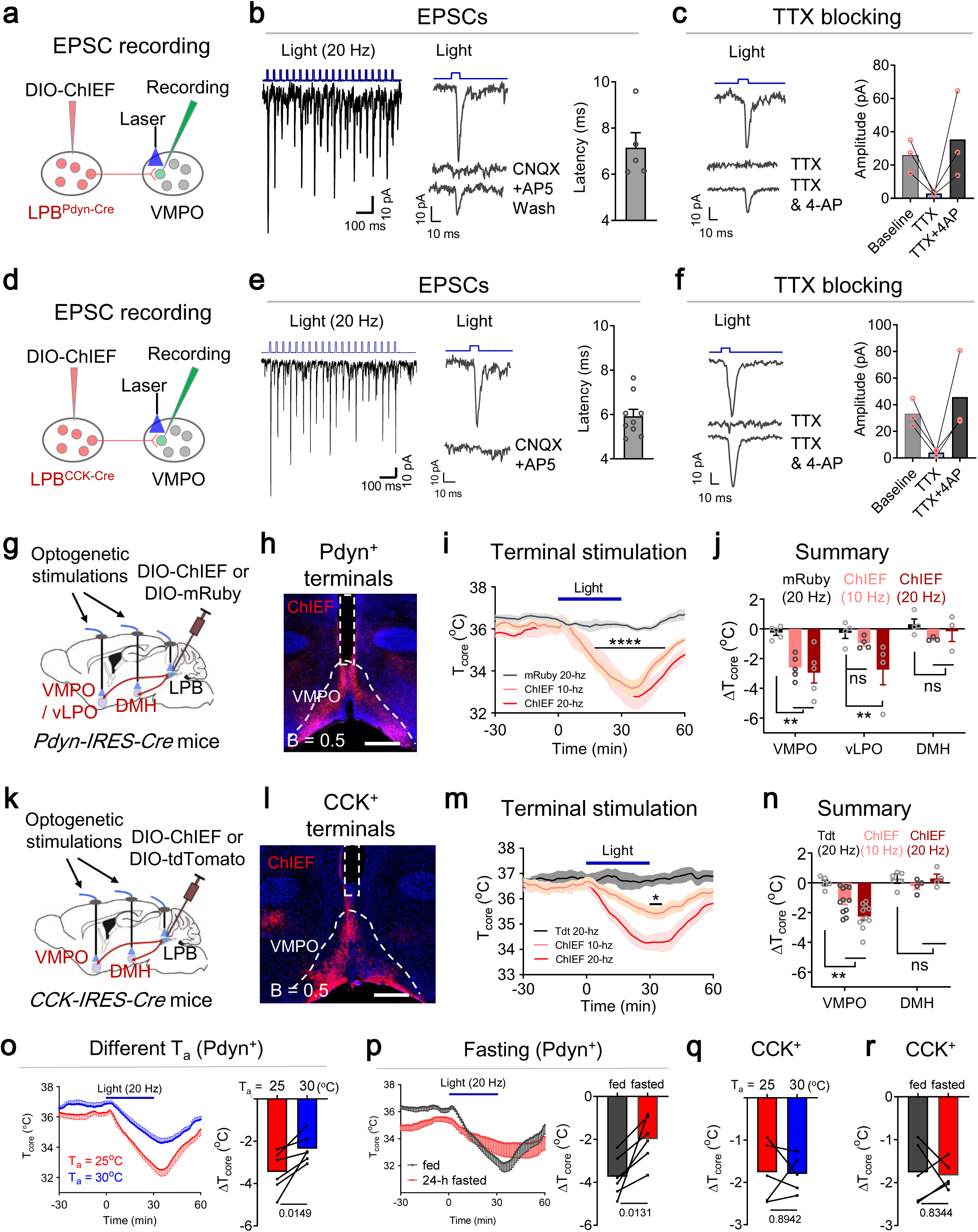
LPB^Pdyn/CCK^→POA circuitries induce hypothermia. **a**, Scheme to record excitatory postsynaptic currents (EPSCs) from VMPO neurons innervated by LPB^Pdyn^ neurons. **b**, Induction of EPSCs in VMPO neurons by light stimulation of LPB^Pdyn & ChIEF^ terminals (blue, 7 mW, 10 ms). EPSCs were blocked by GluR antagonists, D-AP5 and CNQX. **c**, EPSCs were blocked by tetrodotoxin (TTX), and were restored by treatment of 4-Aminopyridine (4-AP). **d**, Scheme to record excitatory postsynaptic currents (EPSCs) from VMPO neurons innervated by LPB^CCK^ neurons. **e**, Induction of EPSCs in VMPO neurons by light stimulation of LPB^CCK & ChIEF^ terminals (blue, 7 mW, 10 ms). EPSCs were blocked by D-AP5 and CNQX. **f**, EPSCs were blocked by TTX, and were restored by treatment of 4-AP. **g**, Scheme for optogenetic stimulation LPB^Pdyn & ChIEF^ neural soma and terminals at the VMPO, the vLPO and the DMH. **h**, The neural terminals in the VMPO projected from LPB^Pdyn & ChIEF^ neurons. **i**, Changes of T_core_ after photoactivation of Pdyn^+^ terminals in the VMPO. **j**, Quantification of ΔT_core_ after photoactivation in the VMPO (n = 5 each), the vLPO (n = 4 each) and the DMH (mRuby, n = 4; ChIEF, n = 3). Laser pattern: 473 nm, 6 mW, Hz as indicated, 30 min. **k**, Scheme for optogenetic stimulation LPB^CCK & ChIEF^ terminals in the VMPO and in the DMH. **l**, The projection of LPB^CCK & ChIEF^ in the VMPO. **m**, Changes of T_core_ after photoactivation of CCK^+^ terminals in the VMPO. **n**, Quantification of ΔT_core_ after photoactivation in the VMPO (tdTomato, n = 5; ChIEF, n = 11) and the DMH (n = 4 each). Laser pattern: 473 nm, 3 mW, Hz as indicated, 30 min. **o**, Changes of T_core_ after photoactivation of LPB^Pdyn^ neuron terminals in the VMPO (n = 6 each) under two different ambient temperatures (T_a_). ΔT_core_ was the difference between t = 0 and t = 30 min. **p**, Changes of T_core_ after photoactivation of LPB^Pdyn^ neuron terminals in the VMPO (n = 6 each) after 24-h fasting. ΔT was the T_core_ difference between t = 0 and t = 30 min. Behavioral tests were performed at T_a_ = 25 °C unless specified. **q-r**, ΔT_core_ after photoactivation of LPB^CCK^ neuron terminals in the VMPO (n = 5 each) under different ambient temperatures (**q**) or after 24-h fasting (**r**). ΔT_core_ was the difference between t = 0 and t = 30 min. Laser pattern for (**o-q**): 473 nm, 6 mW, 20 Hz, 30 min. Scale bars, 500 µm. All data are shown as mean ± s.e.m. The p-values are calculated based on statistical tests in Supplemental Table 1. *p ⩽ 0.05; **p ⩽ 0.01; ****p ⩽ 0.0001; ns, not significant. VMPO, ventromedial preoptic nucleus.

Next, to test the function of these connections, we sought to photoactivate neural terminals in the VMPO. Similar to DREADD activation, photoactivation of LPB^Pdyn^ soma using ChlEF induced hypothermia scaled with laser frequencies (Fig. S11a-d). It did not change the tail temperature (Fig. S11e). Next, we found 5-Hz photoactivation of Pdyn^+^ terminals in the VMPO was sufficient to induce significant hypothermia with a reduction of physical activity and the level of hypothermia saturated after 10 Hz (Fig. 4g-j; Fig. S11f-g). Besides the VMPO, LPB^Pdyn^ neurons send out projections to other regions (Fig. S12), including the vLPO and the DMH. Under 10-Hz stimulations, activation of terminals in the vLPO did not significantly reduce T_core_ (Fig. 4j). Under 20-Hz stimulations, activation of terminals in both the vLPO and the VMPO induced a similar level of hypothermia (Fig. 4j; Fig. S11h-j). There were no changes in T_core_ after activation of terminals in the DMH (Fig. 4j; Fig. S11k-m).

Similar to Pdyn^+^ terminals, activation of CCK^+^ terminals in the VMPO, but not in the DMH caused hypothermia and reduced physical activity (Fig. 4k-n; Fig. S13a-f). In contrast to Pdyn^+^ terminals, the level of hypothermia induced by activation of CCK^+^ terminals increased with laser frequencies until 40 Hz (Fig. 3m-n; Fig. S13c).

Furthermore, we tested whether this induced hypothermia was sensitive to ambient temperatures and energy state. Interestingly, the hypothermia induced by activation of Pdyn^+^ terminals in the VMPO was reduced by warming in 30 °C (Fig. 4o), and was substantially suppressed after 24-h fasting (Fig. 4p; Fig. S11n). In contrast, the hypothermia induced by activation of CCK^+^ terminals was not sensitive to warmth and fasting (Fig. 4q-r; Fig. S13g-h), suggesting the two circuitries may function independently. Together, these results suggest that LPB^Pdyn/CCK^→POA circuitries are sufficient to induce body cooling.

### Differential or categorical regulation of heat-defense variables by LPB^Pdyn/CCK^→ POA circuitries

Thermoregulation variables during heat defense include (at least) inhibition of thermogenesis and activation of vasodilation (Fig. 1). We suspected whether LPB Pdyn or CCK neurons would encode these variables. Therefore, we recorded T_tail_ and T_iBAT_ during photoactivation of their terminals in the VMPO. Interestingly, we did not detect any changes in T_tail_ after activation of Pdyn^+^ terminals (Fig. 5a-b). In contrast, there was a sharp increase in T_tail_ after broad activation of glutamatergic (Vglut2) terminals in the VMPO (Fig. 5b). Next, we recorded T_iBAT_ using a small wireless thermal probe embedded into iBATs. Substantially, we found a rapid decrease of T_iBAT_ after Pdyn^+^ terminal activation, and the decrease of T_iBAT_ was more substantial than that of Vglut2^+^ terminal activation (Fig. 5c), suggesting the Pdyn circuitry is a potent negative regulator for iBAT thermogenesis. Nevertheless, we also tested whether the Pdyn circuitry could inhibit cold-induced muscle shivering activity. We found that Pdyn^+^ terminal photoactivation significantly suppressed cold-induced shivering EMG activity (Fig. 5d; Fig. S14a-c).

**Figure 5.**
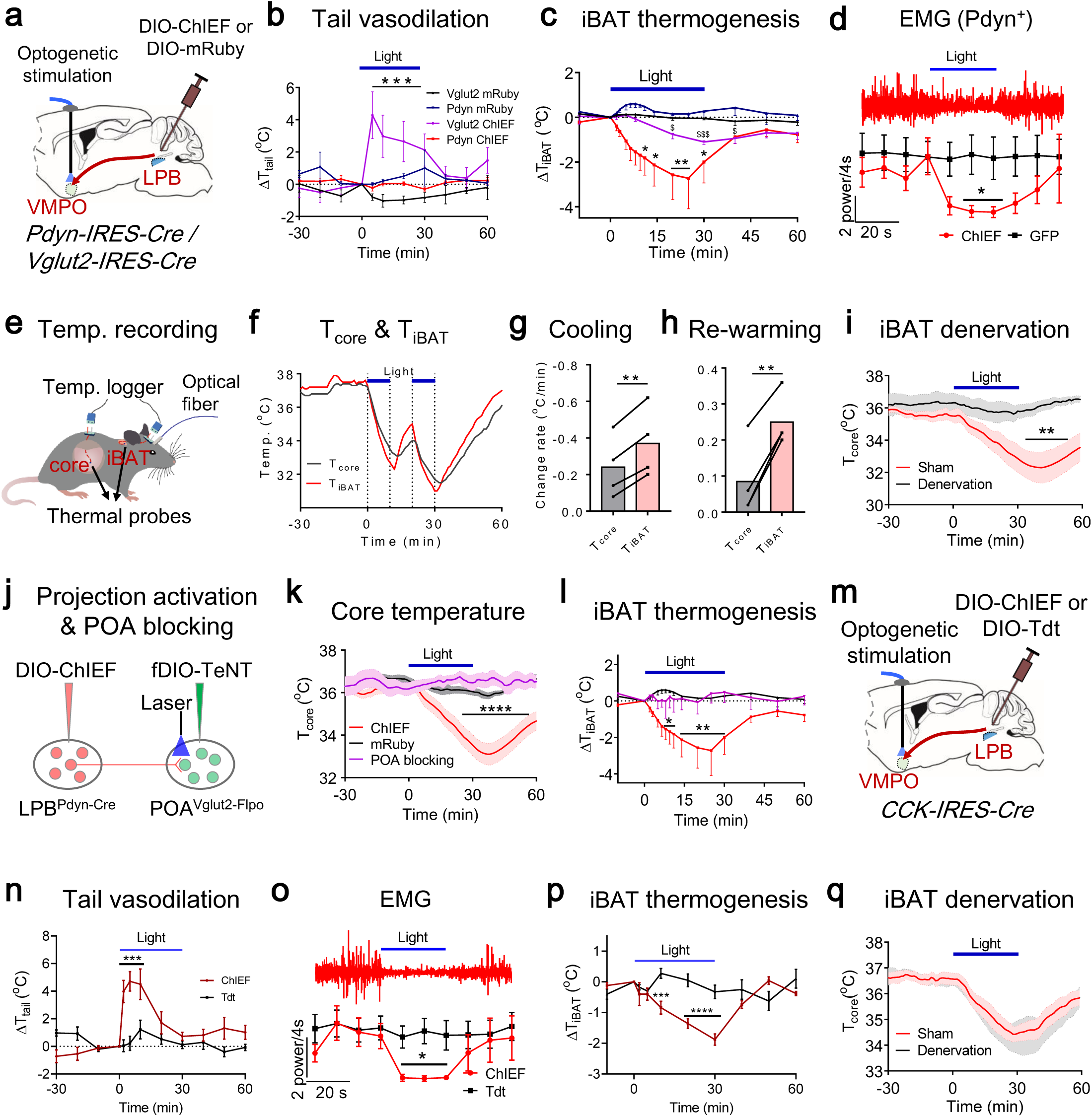
LPB^Pdyn/CCK^→POA circuitries differentially regulate iBAT thermogenesis, vasodilation, and muscle shivering. **a**, Scheme for optogenetic stimulation LPB^Pdyn & ChIEF^ and LPB^Vglut2 & ChIEF^ terminals in the VMPO. **b**, Changes of tail temperatures after photoactivation of terminals from LPB^Vglut2^ and LPB^Pdyn^ neurons (Pdyn ChIEF, n = 7; Pdyn mRuby, n = 3; Vglut2 ChIEF and mRuby, n = 4 each). **c**, Changes of iBAT temperatures after photoactivation of terminals in the VMPO from LPB^Vglut2^ and LPB^Pdyn^ neurons (Pdyn ChIEF, n = 5; Pdyn mRuby, n = 4; Vglut2 ChIEF, n = 4; Vglut2 mRuby, n = 4). **d**, Electromyography (EMG) of the nuchal muscle after cold exposures (0 °C) showed a partial inhibition cold-evoked shivering muscle activities by photoactivation of terminals in the VMPO from LPB^Pdyn^ neurons (ChIEF, n = 5; GFP, n = 6). **e**, Scheme for simultaneous recording of T_core_ and T_iBAT_ by pluggable T-type thermocouples that were directly connected with the logger in freely behaving mice. **f**, An representative trace of T_core_ and T_iBAT_ after photoactivation of terminals in the VMPO from LPB^Pdyn^ neurons. **g-h**, The mean change rate (first 5 min) of T_core_ and T_iBAT_ during body cooling (**g**) and re-warming (**h**) (n = 4 each). **i**, Denervation of the iBAT sympathetic nerves nearly abolished the T_core_ reduction induced by photoactivation of terminals in the VMPO from LPB^Pdyn^ neurons (n = 4 each). **j**, Scheme to block POA glutamatergic neurons while photoactivation of terminals in the VMPO from LPB^Pdyn^ neurons. **k-l**, POA glutamatergic blocking abolished the effect of T_core_ reduction (**k**) (ChIEF, n = 10; mRuby, n = 5; POA blocking, n = 4) and iBAT inhibition (**l**) (ChIEF, n = 5; mRuby, n = 4; POA blocking, n = 4). **m**, Scheme for optogenetic stimulation LPB^CCK & ChIEF^ terminals in the VMPO. **n**, Changes of tail temperatures after photoactivation of terminals in the VMPO from LPB^CCK^ neurons (ChIEF, n = 6; tdTomato, n = 4). **o**, EMG of the nuchal muscle after cold exposures (0 °C) showed a complete inhibition cold-evoked shivering muscle activities by photoactivation of terminals in the VMPO from LPB^CCK^ neurons (ChIEF, n = 4; tdTomato, n = 6). **p**, Changes of iBAT temperatures after photoactivation of terminals in the VMPO from LPB^CCK^ neurons (ChIEF, n = 5; tdTomato, n = 4). **q**, Denervation of the iBAT sympathetic nerves did not affect T_core_ reduction induced by photoactivation of terminals in the VMPO from LPB^CCK^ neurons (n = 8 each). Laser patterns: 473 nm, 6 mW (3 mW in **q**), 20 Hz, 2-s on after 2-s off, time as indicated. All data are shown as mean ± s.e.m. The p-values are calculated based on statistical tests in Supplemental Table 1. *p, ^$^p ⩽ 0.05; **p ⩽ 0.01; ***p, ^$$$^p ⩽ 0.001; ****p ⩽ 0.0001.

The concurrent reduction of the core and iBAT temperatures hints that the lowered BAT thermogenesis might contribute to the reduction of T_core_. However, it is still possible that the lowered T_iBAT_ is a secondary effect of T_core_ reduction. To rule out such a possibility, we simultaneously recorded T_core_ and T_iBAT_ in freely behaving mice at high temporal resolution by using two wired thermocouples that were connected with the logger through pluggable connectors (Fig. 5e). Remarkably, in response to terminal activation, the T_iBAT_ reduced before the T_core_ and reached a lower value than T_core_ (Fig. 5f). Conversely, during rewarming, the T_iBAT_ increased before the T_core_ and reached a higher value than T_core_ (Fig. 5f). The changing rates of T_iBAT_ were significantly larger than those of T_core_ (Fig. 5g-h). Thus, the kinetics rules out the possibility that T_iBAT_ reduction is a side-effect of T_core_ reduction and further supports that the change of iBAT thermogenesis is a driver of T_core_ changes. Next, to directly test whether iBATs are required for T_core_ changes, we denervated their sympathetic nerves and found it indeed blocked the T_core_ reduction (Fig. 5i). Denervation did not affect basal temperatures when kept at 25 °C (Fig. S14f; also see (*56*)).

Next, we suspected POA glutamatergic neurons might be the target of LPB^Pdyn^ neurons since previous reports have shown that POA glutamatergic neurons induced hypothermia (*21, 23, 25*) and we showed that LPB innervated POA neurons were mainly glutamatergic (Fig. S2g). To test this suspicion, we made a Vglut2-2A-Flpo mouse line (Fig. S15a) so that we could simultaneously control the activities of LPB^Pdyn^ neurons and POA^Vglut2^ neurons. We validated its hypothalamic expression pattern by crossing it with a reporter mice (FSF-Tdt) (*57*), and by injection of flippase dependent tetanus neurotoxin (TeNT, AAV-fDIO-TeNT-mCherry) where the mCherry faithfully overlapped with glutamate staining (∼96%) (Fig. S15b-d). Also, we verified the efficacy of TeNT by showing that POA^Vglut2^ blocking indeed impaired both warm- and cold-induced thermoregulation (Fig. S15e-f). Then, we blocked POA^Vglut2-Flpo^ neurons while photoactivation of LPB^Pdyn^ terminals in the VMPO (Fig. 5j). As expected, this blocking abolished the reduction of T_core_ and T_iBAT_ induced by terminal activation (Fig. 5k-l). Yet, blocking LepR neurons in the VMPO did not affect the hypothermia induced by the Pdyn^+^ terminal activation (Fig. S16a-c). Therefore, our data suggest that LPB^Pdyn^ →POA^Vglut2^ circuitry inhibits iBAT thermogenesis to lower body temperature.

Using similar methods, we found that activation of CCK^+^ terminals in the VMPO drastically increased the tail temperature (Fig. 5m-n), suggesting this projection is a potent activator of vasodilation. The cold-induced shivering EMG activity was completely suppressed (Fig. 5o; Fig. S14d-e), which is in contrast to the partial blocking by Pdyn^+^ terminal activation. Also, the T_iBAT_ was reduced (Fig. 5p). To further check the role of iBATs, we denervated their sympathetic nerves and found it did not affect the hypothermia induced by CCK^+^ terminal activation (Fig. 5q), suggesting iBAT thermogenesis was not required for this hypothermia. The reduction of T_iBAT_ was probably due to a secondary effect of body cooling.

Nevertheless, we also checked whether the nucleus of the solitary tract (NTS), the baroreflex control center (*58*), would provide inputs to the LPB to modulate vasodilation. Activation of glutamatergic NTS → LPB pathway indeed induced hypothermia but did not affect tail vasodilation (Fig. S16d-j), suggesting this connection might not be essential for vasodilation. Together, our data define LPB Pdyn and CCK neuron types function to categorically regulate two heat defense variables, inhibition of iBAT thermogenesis and activation of vasodilation, respectively, and to differentially regulate cold-induced muscle shivering.

### Requirement of LPB^Pdyn^→POA circuitry in heat defense, fever limiting, and weight control

The finding that LPB neuron types could regulate heat defense variables predicts that they might be required for heat defense. To test this, we blocked POA-projected LPB Pdyn neurons by targeted expression of TeNT in this circuitry (Fig. 6a). Blocking these Pdyn neurons elevated the basal body temperature and EE (Fig. S17a,c). As expected, this blocking impaired thermoregulation under warm challenge, with a much higher T_core_ than that of the control expressing mCherry (Fig. 6b). In contrast, there was only a small and insignificant effect on the T_core_ during cold exposure (Fig. 6c), suggesting this blocking selectively affects heat defense. Consistent with T_core_ changes, the neural blocking caused slightly yet significantly elevated EE in warm temperature, but not in cold temperature (Fig. 6d; Fig. S17d-e). No significant changes were found in physical activity at the basal level or in warm/cold temperatures (Fig. S17b, f-g).

**Figure 6.**
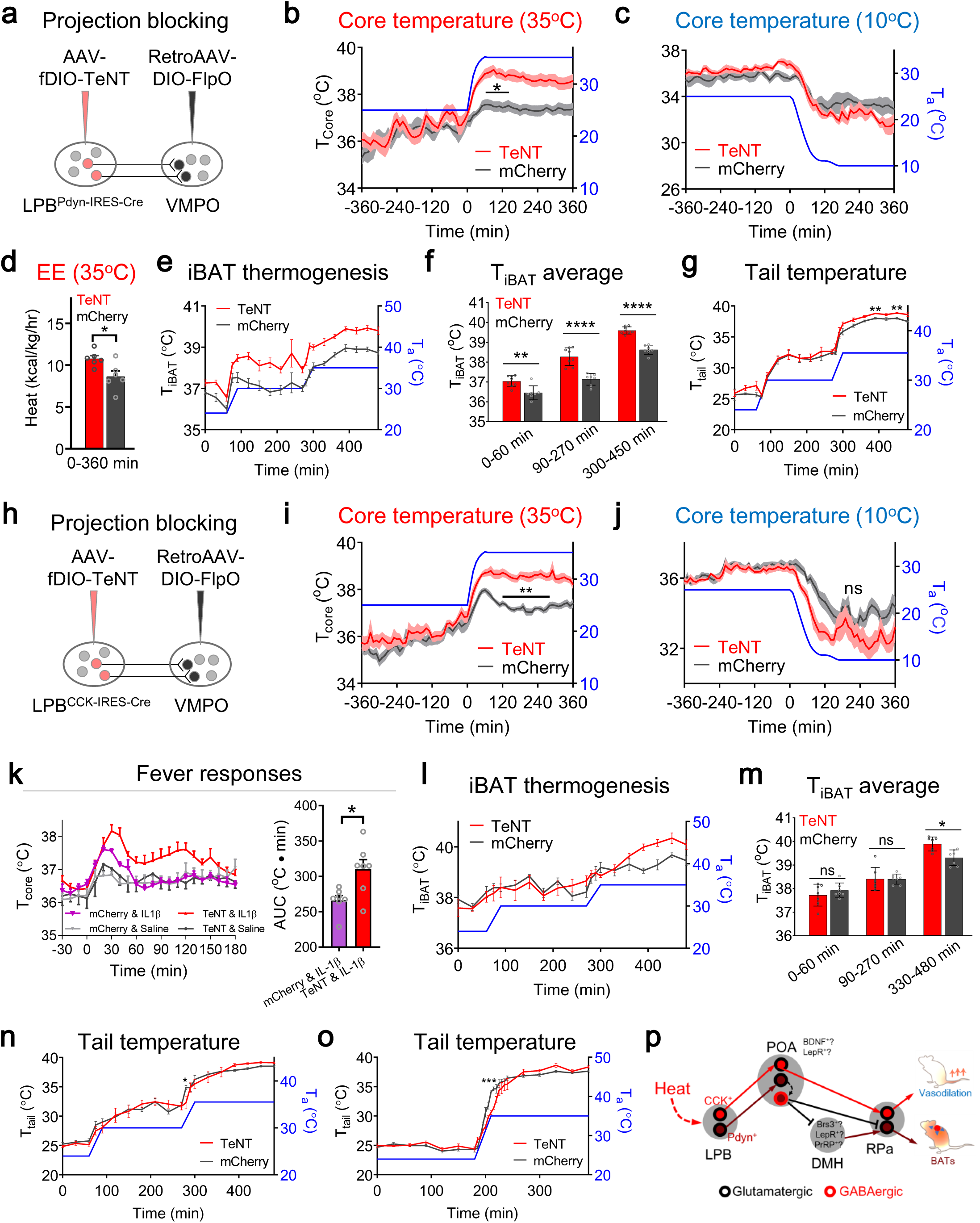
Requirement of LPB→POA circuitries in heat defense, fever limiting, and energy expenditure. **a**, Scheme for blocking of POA-projected LPB^Pdyn^ neurons using neurotoxin TeNT. Retrograde AAVs carrying Cre-dependent FlpO were injected in the VMPO, which drives expression of FlpO-dependent TeNT in the LPB. **b-c**, Changes of T_core_ during the warm challenge (**b**) and cold challenge (**c**) respectively after blocking (n = 6 each). **d**, Changes of energy expenditure (EE) during warm challenge after blocking (n = 6 each). (0 – 360 min) represents the averaged EE between t = (0 – 360 min). **e-f**, Changes of T_iBAT_ (**e**) and their quantification (**f**) under different ambient temperatures (T_a_) recorded by an infrared camera (TeNT, n = 8; mCherry, n = 7). The coat hair on top of the iBATs was shaved. (0– 60 min), (90 – 270 min) and (300 – 450 min) in (**f**) represent the averaged T_iBAT_ between t = (0– 60 min), t = (90 – 270 min), and t = (0 – 360 min), respectively. **g**, Changes of T_tail_ under different T_a_ recorded by an infrared camera (TeNT, n = 8; mCherry, n = 7). **h**, Scheme for blocking of POA-projected LPB^CCK^ neurons using TeNT. **i-j**, Changes of T_core_ during warm challenge (**i**) or cold challenge (**j**) respectively after blocking (TeNT, n = 7; mCherry, n = 8). **k**, Fever responses after injection of IL-1β (n = 7 each group; 10-12 weeks post viral injection). **l-m**, Changes of T_iBAT_ (**l**) and their quantification (**m**) under different T_a_ (n = 7 each). **n-o**, Changes of T_tail_ under different T_a_ (n = 7 each). **p**, Proposed model for LPB^Pdyn/CCK^ circuitries to regulate iBAT thermogenesis and vasodilation. All data are shown as mean ± s.e.m. The p-values are calculated based on statistical tests in Supplemental Table 1. *p ⩽ 0.05; **p ⩽ 0.01; *** p ⩽ 0.001; **** p ⩽ 0.0001; ns, not significant.

We suspected this Pdyn neural blocking may increase iBAT thermogenesis as their activation inhibited the iBAT thermogenesis (Fig. 5c). Interestingly, we found an elevated T_iBAT_ in RT and in warm temperature (Fig. 6e-f). This indicates iBAT adaptive thermogenesis is disinhibited and provides an explanation for the increases in basal body temperature and EE. T_tail_ increased significantly only at the delayed stage after warm exposure (Fig. 6g), which presumably was due to the increased T_core_.

The increased EE after blocking POA-projected LPB^Pdyn^ neurons prompted us to test whether broadly blocking LPB^Pdyn^ neurons would further increase basal EE and affect body weights. Thus, we directly blocked LPB^Pdyn^ neurons with TeNT. As expected, this blocking caused significant increases in basal T_core_ and EE, but not the physical activity and food intake (Fig. S18a-e). It impaired thermoregulation and EE under warm challenge, but not under cold challenge (Fig. S19a-g). Also, it caused more significant increases of T_iBAT_ at RT or in warm temperatures (Fig. S19h) and aggravated fever induced by IL-1β injection (i.p.) (Fig. S19i). Besides, we found that β3 agonist CL316243 evoked a more significant increase in T_core_ after neural blocking when compared to controls (Fig. S18f). Then, we measured body weight changes under chow or high-fat diet (HFD) and found neural blocking significantly reduced the weight gain driven by HFD, but did not change HFD intake (Fig. S18g-h). Moreover, we tested behavioral avoidance to warm and cold temperatures and found it impaired warm avoidance significantly, but not cold avoidance (Fig. S18i-j), suggesting an altered behavioral propensity towards increasing T_core_. Together, we reveal that the LPB^Pdyn^→POA circuitry encodes the inhibition of iBAT thermogenesis in heat defense, regulates thermal behavior, limits fever, and regulates body weight.

### LPB^CCK^→POA circuitry encodes vasodilation during heat defense

To test the necessity of the CCK circuitry in heat defense, we blocked POA-projected LPB CCK neurons by targeted expression of TeNT (Fig. 6h). Similar to Pdyn blocking, it selectively impaired thermoregulation under warm challenge, with a much higher T_core_ than that of the control expressing mCherry (Fig. 6i-j). Also, the blocking increased the fever responses after the injection of il1β (Fig. 6k). However, in contrast to Pdyn blocking, it did not affect basal T_core_, body weight, as well as physical activity and EE in basal levels and after thermal challenges (Fig. S20). Additionally, it did not significantly increase T_iBAT_ until the T_a_ was very high (35 °C) when compared to mCherry controls (Fig. 6l-m). The delayed increase of T_iBAT_ at 35 °C was presumably due to high T_core_ at that ambient temperature (Fig. 6i). Interestingly, the increase of T_tail_ was significantly delayed when switching the T_a_ from 30 to 35 °C as compared to controls (Fig. 6n), suggesting tail vasodilation was impaired. To amplify this defect in vasodilation, we switched T_a_ from 25 to 35 °C and found a more substantial delay of T_tail_ increase when compared to the controls (Fig. 6o). Together, we show that the LPB^CCK^→POA circuitry encodes vasodilation during heat defense.

## DISCUSSION

### Roles of the LPB in thermoregulation

The LPB has been proposed as a relay center to transmit ‘feed-forward’ thermosensory signals into the thermoregulation center, the POA. Whether the LPB is actively involved in processing warm sensory signals is unknown and the genetic identity of LPB thermoregulatory neurons remains poorly characterized. By transcriptome analysis, activity recordings and functional manipulations, we genetically define two subsets of LPB glutamatergic neurons, the Pdyn and CCK neurons as thermoregulatory neurons for autonomic responses to heat defense and fever limiting. Importantly, by finding that Pdyn and CCK neurons categorically encode inhibition of thermogenesis and activation of vasodilation, respectively, and differentially regulate muscle shivering, we suggest that the LPB is a critical center for sorting warm afferent signals that finally leads to differential regulation of peripheral targets. The CCK circuitry represents a selective activator for vasodilation, which might be a promising target for treating thermoregulation disorders such as heat stroke (*1, 2*) and menopause thermal disequilibrium (*4, 48*).

### Thermogenesis control neurons link thermoregulation to weight control

The neural substrates for temperature-induced changes of adaptive thermogenesis are still unclear (*8, 9, 24, 28, 59, 60*), and the interaction and integration between thermal and energy homeostatic systems remain to be better understood (*23, 47, 59, 61, 62*). The BAT represents a common crucial target of the two homeostatic systems, which is a major organ for non-shivering adaptive thermogenesis in cold temperature and contributes to total energy expenditure in mice (up to 60%, (*63*)) and humans (*64, 65*). We reveal that inhibition iBAT thermogenesis is an important hallmark during heat defense and warm-activated LPB^Pdyn^ neurons are selectively required to inhibit BAT thermogenesis to lower body temperature and EE. Blocking LPB^Pdyn^ neurons is sufficient to limit weight gain driven by HFD. Also, LPB^Pdyn^ neural function is sensitive to fasting and warm temperature, which is consistent with the fact that fasting and warmth suppress BAT thermogenesis (*66-70*). Therefore, our molecular identification of a selective negative regulator of BAT thermogenesis may present a genetic entry to investigate the fundamental interconnections between homeostatic systems, which might be helpful for the anti-obesity and diabetic endeavor.

### Thermoregulatory neurons to cold temperature in the LPB and the POA

The LPBd and LPBel are important for thermoregulation in response to both warm and cold temperatures, respectively (*45*),(*16, 18*). Understandably, our broad optogenetic activation of LPB glutamatergic soma can induce hypothermia or hyperthermia (Fig. S2c). However, the LPB in mice is rather small, which prevents subarea-specific functional modulations from separating warm- and cold-induced thermoregulatory neurons. Instead, we sought for genetic and projection-specific approaches to define warm-induced thermoregulatory neurons. Nevertheless, it is still not clear about the genetic identity of thermoregulatory neurons in response to cold temperatures in the LPB.

Similarly, many pieces of evidence show that the POA is a center for thermoregulation to both warm and cold temperatures,(*8, 9, 71-73*). However, several recent reports only support the existence of thermoregulatory neurons in response to warm temperature as activation of various POA cell types only induced hypothermia (*19-21, 23-25*), which contradicts previous lesions and pharmacological studies (*71-74*). By using genetically encoded neurotoxin TeNT, we blocked POA glutamatergic neurons and found they are required for both types of thermoregulation (Fig. S15e-f). Therefore, more specific molecular interrogations are required to reveal mechanisms activated by cold temperatures.

### Role of Pdyn signaling in thermoregulation

The prodynorphin protein is enzymatically cleaved to produce Dynorphins (Dyn), which are endogenous opioids for kappa opioid receptors (KORs). The Dyn/KOR signaling has been suggested to regulate body temperatures besides to induce analgesia for decades (*75-79*). Activation of Pdyn^+^ neurons in the LPB and the arcuate nucleus induce hypothermia (*46*). We show that LPB^Pdyn^ neurons release glutamate to excite POA neurons to induce hypothermia. Also, low-frequency photostimulation (5-Hz), which may not be high enough to trigger the release of neuropeptides, still induces hypothermia (Fig. S11f). Thus, our finding suggests the LPB^Pdyn^ circuitry controls body temperature via releasing glutamate. Interestingly, activation of KOR^+^ neurons in the POA slightly reduced body temperature (*46*), yet it is not surprising as activation of many POA neuron types induces hypothermia (*19-27*). Thus, it would be essential to test how dynorphins are released and whether blocking POA^KOR^ neurons or deleting KOR would block hypothermic responses induced by LPB Pdyn neurons.

### Downstream heat defense pathways

As illustrated in Fig. 6p, we suggest the bifurcation of BAT thermogenesis and vasodilation occurs as early as the LPB in the thermoregulatory reflex pathway. POA glutamatergic neurons are critical for the LPB→POA circuitry based on transsynaptic labeling (Fig. S2g) and POA glutamatergic neural blocking (Fig. 5k). VMPO LepR neurons, which can induce hypothermia (*23*), are not required for LPB^Pdyn^ neuron’s hypothermic effect. VMPO^BDNF/PACAP^ neurons might (in part) be the targets as they can reduce iBAT thermogenesis and increase tail vasodilation (*21, 24*). More specific genetic manipulations are needed to separate the two functions, as well as the shivering control function in the POA. Then, the vasodilation neurons in the POA may target neurons in the raphé pallidus nucleus (RPa) to regulate vasomotor responses (*80, 81*). The DMH might not be needed for vasodilation as broad activation of glutamatergic neurons there does not induce tail vasodilation (*82*). Neurons for inhibiting shivering and BAT thermogenesis appear to share common pathways. First, they excite POA GABAergic neurons such as these in the vLPO (*21, 28*). Then, they may directly inhibit RPa neurons to reduce shivering or BAT thermogenesis (*21, 28, 83, 84*), or they may first inhibit DMH neurons before targeting RPa neurons (*59, 82, 85, 86*) (Fig. 6p). Yet, it should be noticed that neurons for inhibiting shivering and BAT thermogenesis can be separable as LPB^CCK^ neurons, which do not affect BAT thermogenesis, actively suppress shivering activity (Fig. 5o).

## MATERIALS AND METHODS

### Experimental amincals

Animal care and use conformed to institutional guidelines of ShanghaiTech University, Shanghai Biomodel Organism Co., and governmental regulations. All experiments were performed on male adult mice (8–16 weeks old; for TeNT mice, may last to 25 weeks). Mice were housed under controlled temperature (22–25 °C) in a 12-hour reverse light/dark cycle (light time, 8 PM to 8 AM) with ad libitum chow food (4% fat SPF Rodent Feed) and water. The mice strains were listed in the Key Resources Table. Vglut2-T2A-Flpo mice were made through the CRISPR/Cas9 gene targeting by the core facility of Peking Union Medicine College. Briefly, Mus musculus C57BL/6J-Vglut2-T2A-Flpo was generated by inserting a T2A-NLS-Flpo sequence right before the stop codon through CRISPR-Cas9 gene editing. The T2A-NLS-Flpo sequence is:

gagggcagaggaagtcttctaacatgcggtgacgtggaggagaatcccggcccttccatggctcctaagaagaagaggaaggtgatgagccagttcgacatcctgtgcaagaccccccccaaggtgctggtgcggcagttcgtggagagattcgagaggcccagcggcgagaagatcgccagctgtgccgccgagctgacctacctgtgctggatgatcacccacaacggcaccgccatcaagagggccaccttcatgagctacaacaccatcatcagcaacagcctgagcttcgacatcgtgaacaagagcctgcagttcaagtacaagacccagaaggccaccatcctggaggccagcctgaagaagctgatccccgcctgggagttcaccatcatcccttacaacggccagaagcaccagagcgacatcaccgacatcgtgtccagcctgcagctgcagttcgagagcagcgaggaggccgacaagggcaacagccacagcaagaagatgctgaaggccctgctgtccgagggcgagagcatctgggagatcaccgagaagatcctgaacagcttcgagtacaccagcaggttcaccaagaccaagaccctgtaccagttcctgttcctggccacattcatcaactgcggcaggttcagcgacatcaagaacgtggaccccaagagcttcaagctggtgcagaacaagtacctgggcgtgatcattcagtgcctggtgaccgagaccaagacaagcgtgtccaggcacatctactttttcagcgccagaggcaggatcgaccccctggtgtacctggacgagttcctgaggaacagcgagcccgtgctgaagagagtgaacaggaccggcaacagcagcagcaacaagcaggagtaccagctgctgaaggacaacctggtgcgcagctacaacaaggccctgaagaagaacgccccctaccccatcttcgctatcaagaacggccctaagagccacatcggcaggcacctgatgaccagctttctgagcatgaagggcctgaccgagctgacaaacgtggtgggcaactggagcgacaagagggcctccgccgtggccaggaccacctacacccaccagatcaccgccatccccgaccactacttcgccctggtgtccaggtactacgcctacgaccccatcagcaaggagatgatcgccctgaaggacgagaccaaccccatcgaggagtggcagcacatcgagcagctgaagggcagcgccgagggcagcatcagataccccgcctggaacggcatcatcagccaggaggtgctggactacctgagcagctacatcaacaggcggatctga. note=humanized flippase (85-1356), note=NLS (64-84), /note=“T2A”(1..54). Target site 1 (5’-GGTAGCACCGTAAGATTTGGTGG-3’) located on Vglut2 coding exon and target site 2 (5’-GGTTTCCTGTAGAAATAAGTAGG - 3’) located on its 3’ UTR were selected for the integration of the T2A-NLS-Flpo sequence right before the stop codon (TAA). Their corresponding guide sequences are M-Vglut2-E2A-g-up (5’-TAGGTAGCACCGTAAGATTTGG-3’), M-Vglut2-E2A2-g-down (5’-AACCCAAATCTTACGGTGCTA-3’) or M-Vglut2-E2B-g-up (5’-TAGGTTTCCTGTAGAAATAAGT-3’), M-Vglut2-E2B-g-down (5’-AAACACTTATTTCTACAGGAAA-3’) were annealed and inserted into the entry site of pX330, and named pX330-Vglut2-EA, pX330-Vglut2-EB respectively. Together with a donor plasmid pVglut2/ T2A-Flpo contained two arms of Vglut2 gDNAs and a T2A-Flpo sequence. The 5’-arm is an 1111-bp sequence before the Vglut2 stop codon, and 3’-arm is an 1175-bp sequence after the Vglut2 stop codon. A T2A-Flpo coding sequence was inserted between the two arms. To avoid targeting by the sgRNAs, the corresponding sites on the donor were mutated without changing amino acid sequence (CC(A)ACCAAATCT(C)TACGGTGCTACC), and GGTTTCCTGTAGAAATAAGTAGG(C)). The pVglut2/T2A-Flpo, pX330-Vglut2-EA, and pX330-Vglut2-EB were injected into fertilized oocytes and survived embryos were transferred to pseudopregnant females. Mice produced were genotyped by genomic DNA purified from the tail and amplified with LA Taq HS DNA polymerase (Takara, RR042) with the upstream primer (5’-AGAATGGAGGCTGGCCTAAC-3’) and the downstream primer (mu-R: 5’-atgtcgaagctcaggctgtt-3’ and wt-R: 5’ CAAGACTTGCTTGGTTGATATGTT-3’) to produce a 397-bp product for the modified allele and a 217-bp production for the wild-type allele.

### Stereotaxic brain injection

We delivered ∼0.15 μl (unless specified) of AAV virus through a pulled-glass pipette and a pressure micro-injector (Nanoject II, #3-000-205A, Drummond) at a slow rate (23 nl min^−1^) with customized controllers. The injection needle was withdrawn 10 min after the end of the injection. And the injection coordinates were calculated according to Paxinos & Franklin mice brain coordinates (3^rd^ edition) and the injection was performed using a small animal stereotaxic instrument (David Kopf Instruments, #PF-3983; RWD Life Science, #68030; Thinker Tech Nanjing Biotech, #SH01A) under general anesthesia by isoflurane. A feedback heater was used to keep mice warm during surgeries. Mice were recovered in a warm bracket before they were transferred to housing cages for 2–4 weeks before performing behavioral evaluations. The fiber optic cables (200 μm in diameter, Inper Inc., China) were chronically implanted and secured with dental cement (C&B Metabond® Quick Adhesive Cement System, Parkell, Japan). The coordinates are summarized in Table S2, and the bregma sites and anatomy are indicated directly on relevant figures.

### Immunohistochemistry

Mice were anesthetized with isoflurane and perfused transcardially with PBS followed by 4% paraformaldehyde (PFA) in PBS. Brain tissues were post-fixed overnight at 4°C, then sectioned at 50 μm thicknesses using a vibratome (Leica, VT1200S). For glutamate staining, mice were perfused transcardially with PBS followed by 50-mL 4% paraformaldehyde (PFA) in PBS without post-fixation. Brains were isolated and immersed in 20% sucrose for 1 day and 30% sucrose for 2 days at 4°C before being sectioned at 40 μm thicknesses on a cryostat microtome (Leica, CM3050s). Brain slices were collected and blocked with blocking solution containing 2% Normal goat serum (v/v), 2.5% Bovine serum albumin (w/v), 0.1% Triton X-100 (v/v) and 0.1% NaN3 (w/v) in PBS for 2-h at room temperature, and subsequently incubated with primary antibodies for 2 days at 4°C. Then, the samples followed by washing three times in PBST (PBS with 0.1% Triton X-100, v/v) before incubation in secondary antibodies for overnight at 4°C. For glutamate staining, brain slices were collected and blocked with blocking solution containing 10% Normal goat serum (v/v) and 0.1% Triton X-100 (v/v) in PBS for overnight at 4°C, and subsequently incubated with primary antibodies for 8-h at room temperature. After that, the slices were washed three times in PBST (PBS with 0.7% Triton X-100, v/v) before incubation in secondary antibodies for 2-h at room temperature. Primary antibodies include chicken anti-GFP (Abcam, #ab13970, 1: 1000), rabbit anti-Neuropeptide S (Abcam, #ab18252, 1: 500), rabbit anti-cfos (Synaptic systems, #226003, 1: 10000), guinea pig anti-cfos (Synaptic systems, #226004, 1: 500), rat anti-RFP (Chromotek, #5F8, 1: 1000), rabbit anti-Glutamate (Sigma, #G6642-.2ML, 1: 1000; see (*87-89*) for validation.), rabbit anti-Dynorphin A (Phoenix Pharmaceuticals, #H-021-03, 1:500) and Guinea pig anti-ProDynorphin (GeneTex, GTX10280, 1: 500). Secondary antibodies include DyLight 488 conjugated goat anti-chicken IgG (Invitrogen, #SA5-10070, 1: 1000), Alexa Fluor 594 conjugated goat anti-rat IgG (Invitrogen, #A-11007, 1: 1000), Alexa Fluor 594 conjugated goat anti-rabbit IgG (Jackson, #111-585-144,1: 1000), Alexa Fluor 594 conjugated Goat Anti-Guinea Pig IgG (Invitrogen, #A-11076, 1: 1000) and Alexa Fluor 647 conjugated goat anti-rabbit IgG (Invitrogen, #A21244, 1: 1000). Brain sections were then washed three times in PBST and cover-slipped with DAPI Fluoromount-G mounting medium (SouthernBiotech, #0100-20). Images were captured on a Nikon A1R or Leica SP8 confocal microscope or Olympus VS120 Virtual Microscopy Slide Scanning System.

In order to stain Pdyn and Dynorphin A, mice were treated with 40 μg of colchicine (Sigma, C9754) in 1-μl H_2_O through intra-cerebra-ventricular injection with a pulled-glass pipette and a pressure micro-injector (Nanoject II, #3-000-205A, Drummond) at a slow rate (46 nl min^−1^) with customized controllers. Coordinates were 1.0-mm lateral, 0.4-mm back from the bregma and 2.5-mm below the skull. Injection began 5 min after tissue penetration and pulled-glass pipette was pulled out 10 min post-injection. After suturing, mice were recovered in a warm bracket before they were transferred to housing cages for 72-h before perfusion.

### Cell-type specific retro-TRAP sequencing

A recombinant Cre-dependent AAV plasmid expressing GFP-tagged ribosomal subunit L10a (AAV2-EF1a-DIO-EGFP-L10a) was gifted by Dr. Jeff Friedman and then packaged using rAAV2-retro capsids by Taitool. 300 nl of the viral aliquot was injected into the VMPO of Vglut2-IRES-Cre mice. Animals were sacrificed for immunoprecipitation 4 weeks after injection.

For TRAP experiments, brain slices containing the LPB regions were prepared from virus injected mice anesthetized with isoflurane before decapitation. Brains were removed and placed in ice-cold DEPC-PBS. Coronal brain slices (300 μm) were cut using a vibratome (VT 1200S, Leica Microsystems, Germany). The slices were incubated at ice-cold dissection buffer in a 60-mm dish and placed under a stereomicroscope. Then the two LPB regions of each slice were cut out with microsurgical forceps with a 150-μm width tip. Tissue from 6 brains was pooled for each experimental repeat, and a total of three experimental repeats were performed. Pooled tissue was homogenized and clarified by centrifugation. Ribosomes were immunoprecipitated using anti-EGFP antibodies (an equal mixture of clones 19C8 and 19F7 from Monoclonal Antibody Core Facility at Memorial Sloan-Kettering cancer center) that were previously conjugated to Protein L coated magnetic beads (ThermoFisher Scientific) for 16-h in 4°C with rotating. An aliquot of input RNA taken prior to immunoprecipitation and total immunoprecipitated RNA was then purified using the RNAeasy Micro kit (QIAGEN). The products were then purified and enriched with PCR to create the final cDNA library. Purified libraries were quantified by Qubit® 2.0 Fluorometer (Life Technologies, USA) and validated by Agilent 2100 bioanalyzer (Agilent Technologies, USA) to confirm the insert size and calculate the mole concentration. The cluster was generated by cBot with the library diluted to 10 pM and then was sequenced on the Illumina NovaSeq 6000 (Illumina, USA). The library construction and sequencing were performed by Shanghai Sinotech Genomics Corporation (China).

### TRAPseq data analysis

The RNA quantification data after sequencing were analyzed as the following. First, the RNA in the IP for each gene was divided by the input, to determine the statistical significance and fold-enrichment (IP/Input). The hits were narrowed down by only analyzing the PB-enriched gene list downloaded from Allen Institute (https://alleninstitute.org/; Tool one: MOUSE BRAIN CONNECTIVITY-source search-filter source structure: Parabrachial nucleus; Tool two: MOUE BRAIN-Fine Structure Search: Parabrachial nucleus). The top candidate genes were then selected for further quantitative PCR analysis.

### Metabolic measurement

For DREADDs and TeNT mice, energy expenditure, locomotor activity, food intake, and body temperature were monitored by Comprehensive Lab Animal Monitoring System with Temperature Telemetry Transmitter (CLAMS; Columbus Instruments, with G2 E-Mitter transponders). The data was acquired at a 10-min interval per data point as shown in the figures. Temperature transponders were implanted into the peritoneal cavity 3 – 5 days before testing. Stimuli (drugs, or temperature) were delivered between 10 am and 3 pm (for long time stimuli, it may occur between 2 pm to 8 pm) in the dark phase. Mice were adapted in the metabolic chambers for 2 days before giving saline (volume (μl) = 10 x body weight (grams)), CNO (ENZO, #BML-NS105-0025, i.p., 0.1 - 2.5 mg/kg body weight, as indicated in each figure), and CL316243 (Merck, #5.04761.0001, i.p., 1 mg/kg body weight), IL-1β (SIGMA, #SRP8033, i.p., 3 μg/kg body weight).

### Tail, iBAT, and core body temperature measurement

For C57BL/6J and TeNT (as to their controls) injected mice, tail temperatures and brown adipose tissue temperatures were measured using an infrared thermal camera (A655sc, FLIR). Snapshot images were taken at the specified time points and an average of three spot readings was taken at the middle of the tail and interscapular region which was shaved 3 – 5 days before measurement. For optogenetic mice, the brown adipose tissue temperature was measured three times for average using subcutaneous implanted thermal probes (IPTT-300, Biomedic Data Systems, DE) inserted at the midline in the interscapular region under anesthesia at least one week before the start of the experiment. Measurements were taken with the non-contact DAS-7007 reader. Core temperature and physical activity of optogenetic mice were recorded at a 1-min interval by VitalView Data Acquisition System Series 4000 (Mini-Mitter, Oregon, USA) with G2 E-Mitter transponders. All measurements were conducted during the dark phase.

For wired recording, we using a modified thermocouple temperature measuring device. Two T-type thermocouples (TT-40) were implanted in the abdomen and interscapular region of the mice, respectively, to measure core body temperatures and brown fat temperatures. Before implantation, we tested the linear temperature change to make sure that the measurement was accurate. And the implantation was done 3 - 5 days before measurement. NI data acquisition card (USB-TC01) and supporting software were used for data collection. The sampling rate was 1 Hz.

### iBAT sympathetic nerve denervation

Mice were anesthetized with isoflurane. The hair in the targeted area was shaved and the area was wiped with 95% ethanol-soaked sterile gauze. A midline incision was made in the skin along the upper dorsal surface to expose both iBAT pads followed by additional application of styptic powder to the area to prevent bleeding. We gently exposed the medial, ventral surface of both pads to visualize nerves beneath the pad. Nerves were checked under a microscope. All five nerves of both pads were cut off a length of 2-3 mm to avoid nerve regeneration. Mice were housed at the thermoneutral environment for 5-7 days for recovery before measurements.

### Electromyogram (EMG) recording

Mice were anesthetized with isoflurane and fixed with the animal stereotaxic instrument to avoid movement. The mouse abdomen was put on an aluminum thermal pad whose temperature was controlled by circulating water to maintain body temperature at 36 ± 1.5°C. Rectal temperature was monitored with a thermocouple as an indication of body core temperature. Another thermocouple to monitor skin temperature was taped onto the abdominal skin which was put on the thermal pad. Handmade electrodes for EMG recording were inserted into nuchal muscles and the signal was filtered (10–1000 Hz) and amplified (×1000) with a Model 1800 2-Channel Microelectrode AC Amplifier (A-M Systems, Sequim, WA, USA). To induce shivering, the circulating water was switched to ice-cold water, and each cooling episode lasted about 10 minutes until the laser stimulations were done. The EMG amplitude was quantified (Spike 2, CED, Cambridge, UK) in sequential 4-sec bins as the square root of the total power (root mean square) in the 0–500 Hz band of the auto spectra of each 4-s segment of EMG as shown in (*90*). Temperatures were recorded by NI data acquisition card (USB-TC01) and supporting software.

### Temperature preference test

Mice were separated into a new cage for at least seven days. A custom 2-chamber cage was placed on temperature-controlled plates, adjusted such that one end of the cage was at 32 ± 0.5°C, while the other end of the cage was at 20 or 35 ± 0.5°C. Mice were allowed to explore the chamber for 30-min per day for three days. During this habituation period, both of the two plates were adjusted to 32 ± 0.5°C. On the fourth day, the plates were adjusted to either 32 ± 0.5°C or 20 (35) ± 0.5°C. During this testing phase, individual mice were put into either chamber and allowed to freely explore all chambers for 30 min. For each trial, the orientation of the plate and cage was randomized. The position of the mouse was recorded using Digbehv video-recording software (Jiliang, Shanghai, China). All tests were performed from 10 am and 5 pm in the dark phase.

### Food intake assay and blood glucose measurement

For the DREADD experiment shown in Fig. S8i and 9f, the food intake assays were performed in the home cage. Mice were separated for at least 5 days and then fasted for 24 hours from 10 am – 10 am before the assay. Mice were injected with saline or CNO, and measurement of food intake was made at 1, 2, 3, 4 and 5-hr post-injection. CNO injections were at a concentration of 2.5 mg/kg body weight. For the TeNT blocking experiment shown in Fig. S18d, food intake was monitored by the Comprehensive Lab Animal Monitoring System with free access to food. Blood glucose concentrations were measured from the tail vein using a hand-held glucometer (Accu-Chek Performa Connect, Roche, Switzerland). CNO injections were at a concentration of 2.5 mg/kg body weight.

### Slice physiological recording

Brain slices recording and data analysis were performed similarly as described in (*91*). Briefly, slices containing the VMPO regions were prepared from adult mice anesthetized with isoflurane before decapitation. Brains were removed and placed in ice-cold oxygenated (95% O_2_ and 5% CO_2_) cutting solution (228 mM sucrose, 11 mM glucose, 26 mM NaHCO_3_, 1 mM NaH_2_PO_4_, 2.5 mM KCl, 7 mM MgSO_4_, and 0.5 mM CaCl_2_). Coronal brain slices (250 μm) were cut using a vibratome (VT 1200S, Leica Microsystems, Germany). The slices were incubated at 32°C in oxygenated artificial cerebrospinal fluid (ACSF: 119 mM NaCl, 2.5 mM KCl, 1 mM NaH_2_PO_4_, 1.3 mM MgSO_4_, 26 mM NaHCO_3_, 10 mM glucose, and 2.5 mM CaCl_2_) for 1 h, and were then kept at room temperature under the same conditions before transfer to the recording chamber. The ACSF was perfused at 2 ml/min. The acute brain slices were visualized with a 40x Olympus water immersion lens, differential interference contrast (DIC) optics (Olympus Inc., Japan), and a CCD camera (IR1000, Dage-MTI, USA).

Patch pipettes were pulled from borosilicate glass capillary tubes (#BF150-150-86-10, Sutter Instruments, USA) using a P-97 pipette puller (Sutter Instruments, USA). For EPSC recordings, pipettes were filled with solution (in mM: 135 potassium gluconate, 4 KCl, 10 HEPES, 4 Mg-ATP, 0.3 Na_3_-GTP, 10 sodium phosphocreatine) (pH 7.2, 275 mOsm). For EPSC recordings, cells were clamped at −70 mV; to block EPSCs, 10 μM CNQX (Sigma) and 50 μM DL-AP5 (Tocris) was applied to the bath solution. The resistance of pipettes varied between 3–5 MΩ. The signals were recorded with MultiClamp 700B, Digidata 1440A interface and Clampex 10 data acquisition software (Molecular Devices). After the establishment of the whole-cell configuration, series resistance was measured. Recordings with series resistances of > 20 MΩ were rejected. Blue light pulses were delivered through the 40X objective of the microscope, with the X-Cite LED light source. Light power density was adjusted to 7 mW/mm^2^.

### Calcium fiber photometry

Following injection of an AAV2/9-hSyn-Flex-GCaMP6s or AAV2/9-hSyn-DIO-jGCaMP7b-WPRE-pA viral vector, an optical fiber (200 μm O.D., 0.37 numerical aperture, Inper Inc., China) was placed 150 μm above the viral injection site. Post-surgery mice were allowed to recover for at least 2 weeks before any studies were performed. Fluorescence signals were acquired with a dual-channel fiber photometry system (Fscope, Biolinkoptics, China) equipped with a 488-nm excitation laser (OBIS, Coherent), 505-544 nm emission filter and a photomultiplier tube (Hamamatsu, #R3896). The gain (voltage) on PMT was set to 600 V. The laser power at the tip of the optical fiber was adjusted to as low as possible (∼ 25 μW) to minimize bleaching. The analog voltage signals fluorescent were low-pass (30 Hz) filtered and digitalized at 100 Hz, which were then recorded by Fscope software (Biolinkoptics, China). Mice behaviors were recorded by top-view positioned camera (Logitech, #C930e). We used a screen recorder (EV Capture, Hunan Yiwei LTD, China) to synchronize behavioral videos and analog voltage signals of fluorescence.

All animals were allowed to recover after fiber cord attachment for at least 30 min. For temperature challenges, the custom-build Peltier floor plate (15 × 15 cm) temperature for each individual mouse was controlled by a Peltier controller (#5R7-001, Oven Industry) with customized Labview code (National Instrument), and mouse was caged in an acrylic chamber (15 × 15 x 30 cm) which placed on the plate.

For capsaicin/menthol/icillin and other tested stimuli presentations, experiments were conducted in a fresh empty housing cage. In capsaicin/menthol/icilin experiments, drugs were prepared freshly on the day of experiment by dilution in PBS and injected intraperitoneally at a volume (mL) equal to 10X body weight (g) to achieve a final dosage of 1/2/4 mg/kg (capsaicin) or 10 mg/kg (menthol) or 2.5 mg/kg (icillin). Capsaicin and menthol responses were assayed on separate days. For other tested stimuli presentation, the following items were sequentially placed into the cage for 30-min each: novel object (plastic ball and aluminum block), normal chow, high fat diet, novel mouse of the same sex.

### Fiber photometry data analysis

The data analysis has been described previously (*21, 88*). Basically, to calculate the fluorescence change ratios, the raw data were analyzed using Fscope software and customized MATLAB code. We segmented the data based on behavioral events within individual trials. We derived the values of fluorescence change (ΔF/F_0_) by calculating (F−F_0_)/F_0_, where F_0_ is the baseline fluorescence signal averaged in a time window before events (temperature changes, drug injections, and introductions of food, novel objects or mouse as indicated) indicated in the figure legends. Finally, we plotted the average of fluorescence changes (ΔF/F_0_) with different behavioral events using Customized MATLAB code and GraphPad Prism 8 (GraphPad).

### Data availability

The data that support the findings of this study are available from the corresponding author upon reasonable request.

### Code availability

The code that supports the findings of this study is available from the corresponding author upon reasonable request.

## SUPPLEMENTAL MATERIALS

Figure S1-20

Table S1-2

Movie S1

## ACKNOWLEDGMENTS

We thank Drs. Ji Hu, Minmin Luo, Hailan Hu, Cheng Zhan, Peng Cao, and Haohong Li for reagent share; Dr. Xiaoming Li and the Molecular Imaging Core Facility (MICF) of School of Life Science and Technology, ShanghaiTech University for Microscopic imaging; Dr. Ying Xiong and the Molecular Cellular Core for slices and staining; All Shen lab members and “Shen Xian Hui (NPC)” Wechat group for valuable discussion. This study is funded by National Key Research and Development Program of China (2019YFA0801900), National Nature Science Foundation of China (W. Shen, 31771169 and 91857104), Shanghai Science and Technology Committee of Shanghai City (19140903800), Shenzhen-Hong Kong Institute of Brain Science-Shenzhen Fundamental Research Institutions (W. Shen), and Innovation Program of Shanghai Municipal Education Commission (W. Shen). We thank the staff members of the Animal Facility at the National Facility for Protein Science in Shanghai (NFPS), Zhangjiang Lab, China for providing technical support and assistance.

## AUTHOR CONTRIBUTIONS

Yang W and Du X performed most of the experiments. Gao C, Xie H, Xiao Y, Liu J, Xu J performed behavioral evaluations. Zhang W performed the electrophysiology. Jia X performed retroTRAP analysis. Ni X, Fu X, and Tu H performed the immunostaining. Fu X designed and performed a concurrent recording of BAT and core temperatures. Yang W, Yang J, He M, Wang H, Yang H, Xu X, and Shen W designed the experiments. Yang W, Du X, and Shen W wrote the manuscript.

---

**Supplemental Fig. 1.**
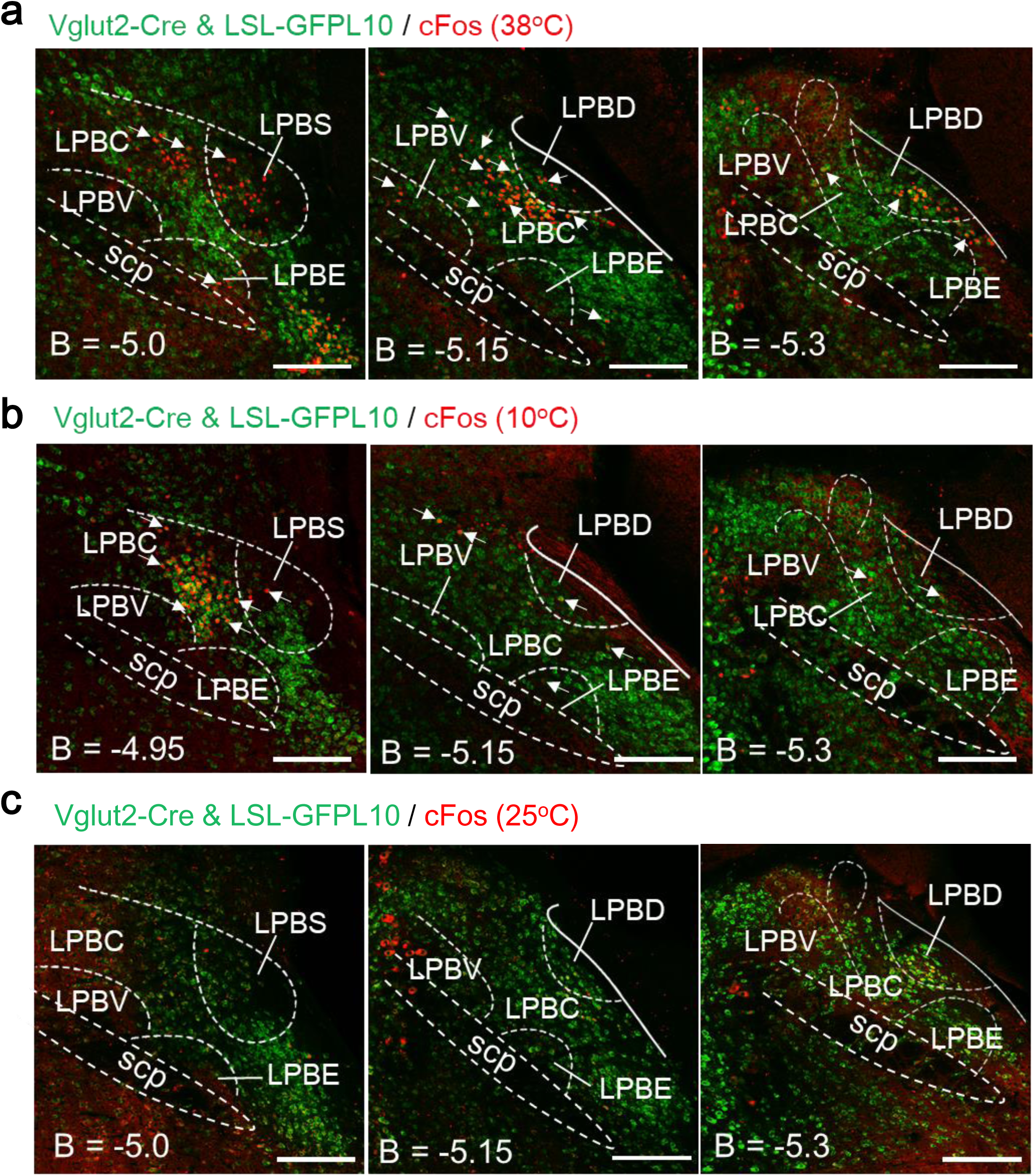
Thermal responses of LPB glutamatergic neurons, related to Fig. 1. **a**, The overlapping between warm-induced cFos and glutamatergic marker Vglut2-IRES-Cre & LSL-GFPL10 in the LPB. The merge ratio is ∼95 % of cFos^+^ cells (297 merged / 312 cFos^+^ cells = 95.2 %, n = 3). **b**, The overlapping between cold-induced cFos and glutamatergic marker Vglut2-IRES-Cre & LSL-GFPL10 in the LPB. The merge ratio is ∼95 % of cFos^+^ cells (224 merged / 237 cFos^+^ cells = 94.5 %, n = 3). **c**, The overlapping between cFos at the room temperature (25 °C) and glutamatergic marker Vglut2-IRES-Cre & LSL-GFPL10 in the LPB. The merge ratio is ∼73 % of cFos^+^ cells (23 merged / 37 cFos^+^ cells = 72.9 %, n = 3). Scale bars, 200 µm. Arrows, examples of merged cells. B, bregma; LPB, lateral parabrachial nucleus; LPBC, lateral parabrachial nucleus, central part; LPBD, lateral parabrachial nucleus, dorsal part; LPBE, lateral parabrachial nucleus, external part; LPBV, lateral parabrachial nucleus, ventral part; LPBS, lateral parabrachial nucleus, superior part; scp, superior cerebellar peduncle.

**Supplemental Fig. 2.**
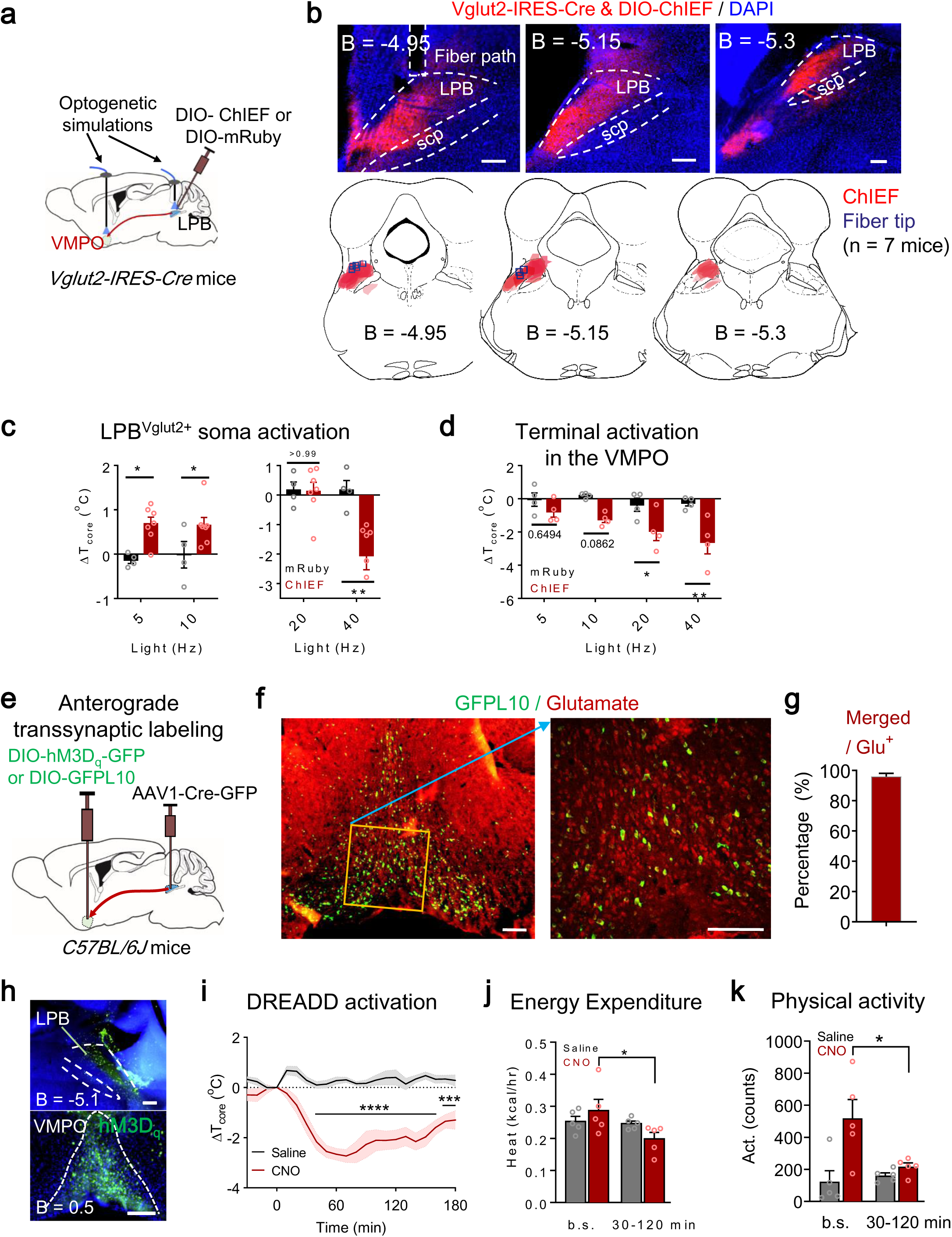
Roles of the LPB→POA circuitries in thermoregulation, related to Fig. 1. **a**, Scheme of optogenetic activation of glutamatergic (Vglut2) LPB soma and VMPO terminals. **b**, The anatomical map of DIO-ChIEF expression and the positions of optical fiber tips in the LPB. The ChIEF expression on the map was shown in red and the optical fiber tips were shown in blue rectangles. **c**, Changes of T_core_ after soma photoactivation of LPB^Vglut2^ neurons revealed a heterogeneous population that could induce hypothermia or hyperthermia (ChIEF, n = 7; mRuby, n = 4). Laser pattern: 473 nm, 3 mW, Hz as indicated, 30 min. **d**, Changes of T_core_ after photoactivation of neural terminals in the VMPO projected from LPB^Vglut2^ neurons (n = 4 each). Laser pattern: 473 nm, 6mW, Hz as indicated, 30 min. **e**, Scheme for anterograde transsynaptic labeling of VMPO neurons by LPB-injected AAV1-CMV-Cre-GFP. **f**, Viral expression of AAV1-CMV-Cre-GFP injected in the LPB which drives expression of DIO-GFPL10 in the VMPO. **g**, The overlapping of DIO-GFPL10 and glutamate staining in the VMPO. **h**, Viral expression of AAV1-CMV-Cre-GFP injected in the LPB which drives expression of DIO-hM3D_q_ in the VMPO. **i-k**, DREADD activation of VMPO neurons transsynaptically labeled by LPB neurons reduced T_core_ (**g**), EE (**h**) and physical activity (**i**) (n = 5). b.s. represents baseline, which is the averaged EE or physical activity between t = (– 30 – 0 min); (30 – 120 min) represents the averaged EE or physical activity between t = (30 – 120 min). CNO, Clozapine N-oxide. Scale bars, 200 µm. All data are shown as mean ± s.e.m. All the p-values are calculated based on ordinary two-way ANOVA Bonferroni’s corrections except in (**i**) (repeated measures two-way ANOVA with Bonferroni’s corrections). *p ≤ 0.05; ***p ≤ 0.001; ****p ≤ 0.0001; ns, not significant. VMPO, ventromedial preoptic nucleus; LPB, lateral parabrachial nucleus; scp, superior cerebellar peduncle.

**Supplemental Fig. 3.**
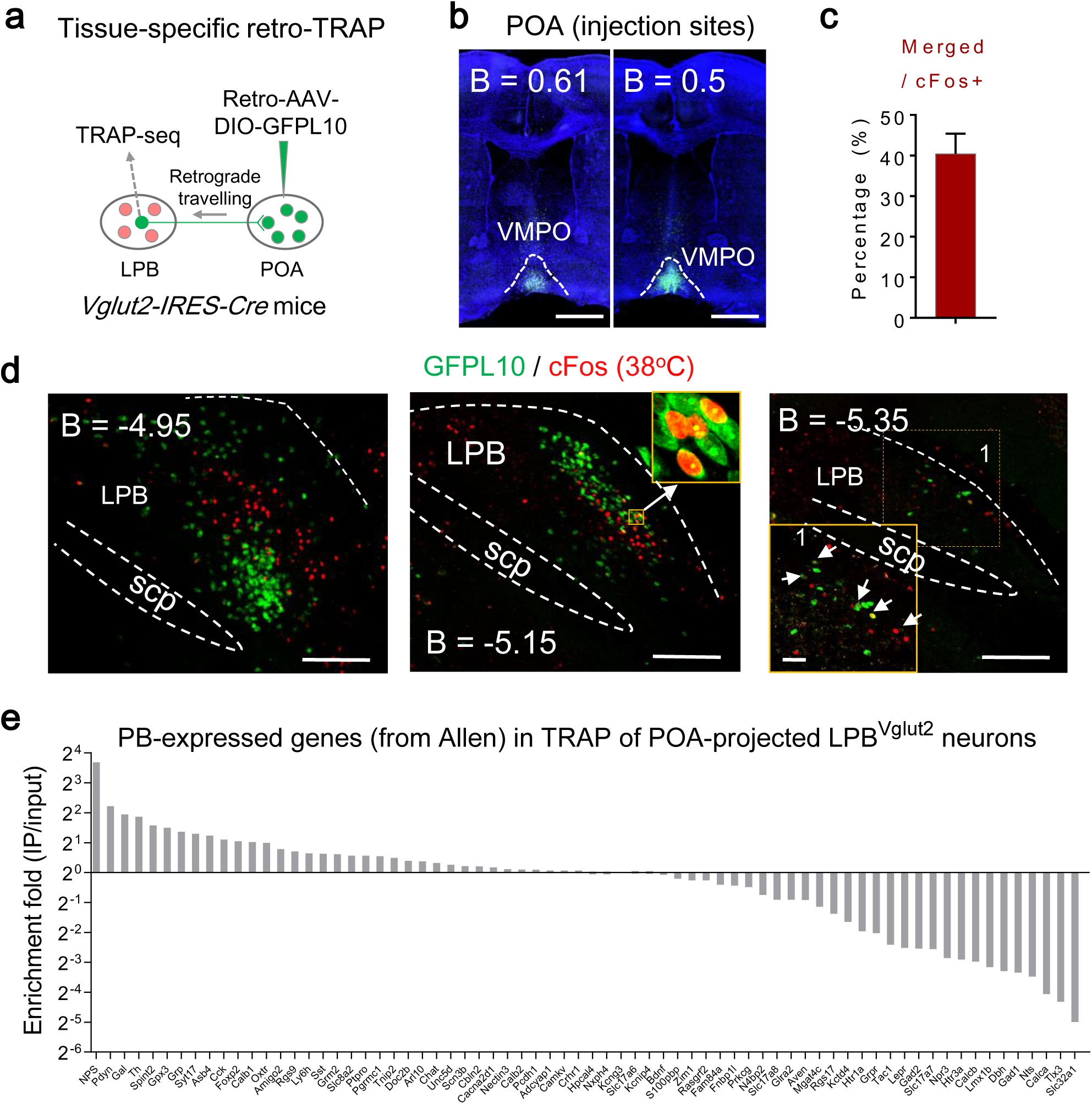
Tissue-specific retro-TRAPseq of VMPO-projected LPB glutamatergic neurons, related to Fig. 1. **a**, Scheme of tissue-specific retro-TRAP, where translational ribosomes from VMPO-projected LPB^Vglut2^ neurons were immunoprecipitated and associated mRNAs were used for sequencing. **b** & **d**, Expression of retro AAV2-DIO-GFPL10 in the injections sites (the VMPO) (**b**) and the LPB (**d**) of Vglut2-IRES-Cre mice. Scale bars, 200 µm. Arrows, examples of merged cells. **c**, Quantification of the overlapping ratio (76 merged / 188 cFos^+^ cells = 40.4 %, n = 3) between warm-induced cFos and GFPL10 in the LPB showed in (**d**). **e**, Ranking of PB-expressed genes downloaded from Allen Institute according to their enrichment fold in the retro-TRAP.

**Supplemental Fig. 4.**
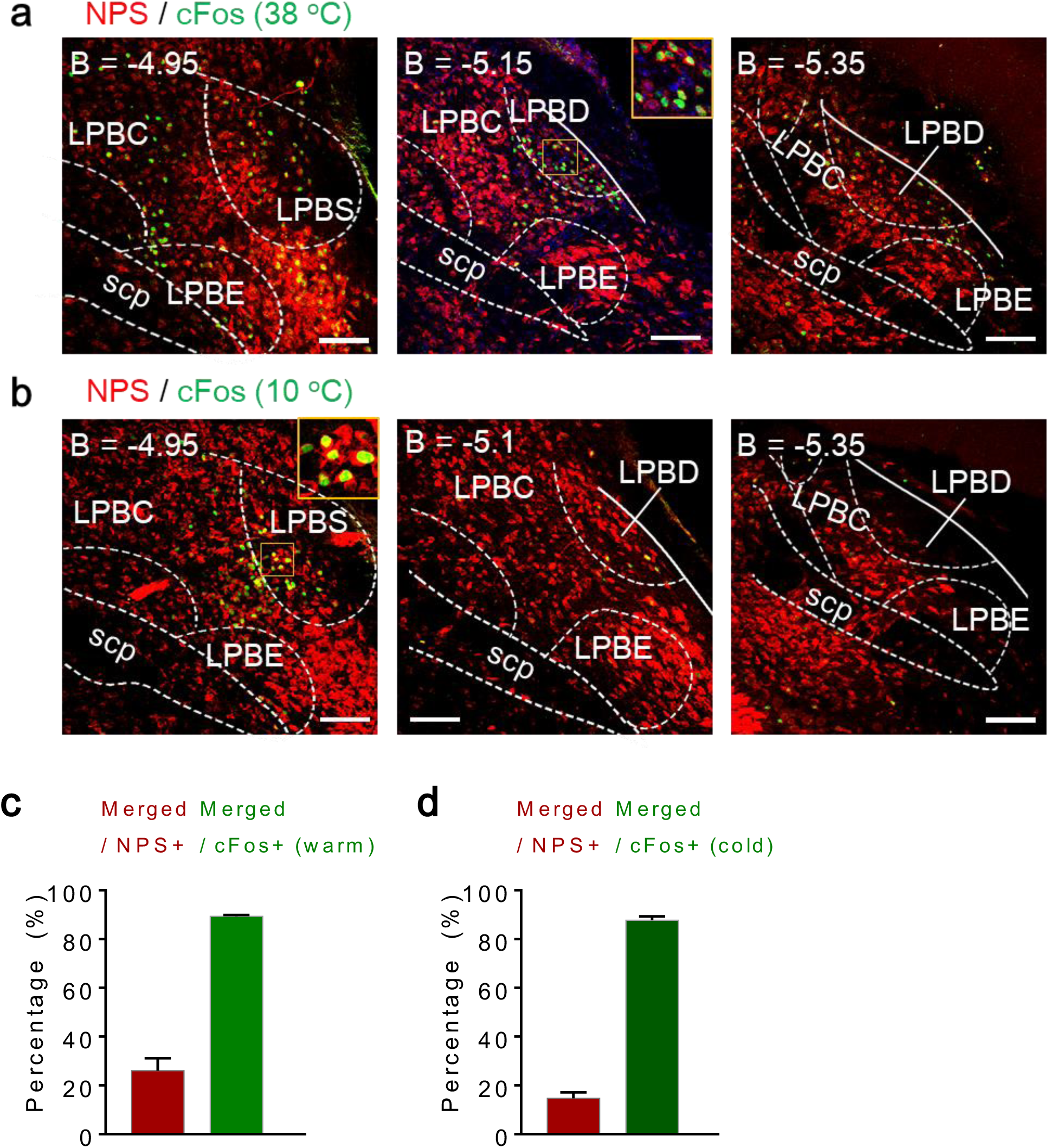
Properties of LPB NPS neurons, related to Fig. 2. **a, c**, The overlapping between NPS and warm-induced cFos immune activity in the LPB at different bregma sites. The merge ratio was ∼89.5 % of cFos^+^ cells (84 merged / 94 cFos^+^ cells = 89.48 %, n = 2). The number of merged cells was ∼22 % of NPS^+^ cells (84 merged / 377 NPS^+^ cells = 22.3 %). **b** & **d**, The overlapping between NPS and cold-induced cFos immune activity in the LPB at different bregma sites. The merge ratio was ∼89.6 % of cFos^+^ cells (79 merged / 88 cFos+ cells = 89.60 %, n = 2). The number of merged cells was ∼15 % of NPS^+^ cells (79 merged / 539 NPS+ cells = 14.7 %). Scale bars, 100 µm.

**Supplemental Fig. 5.**
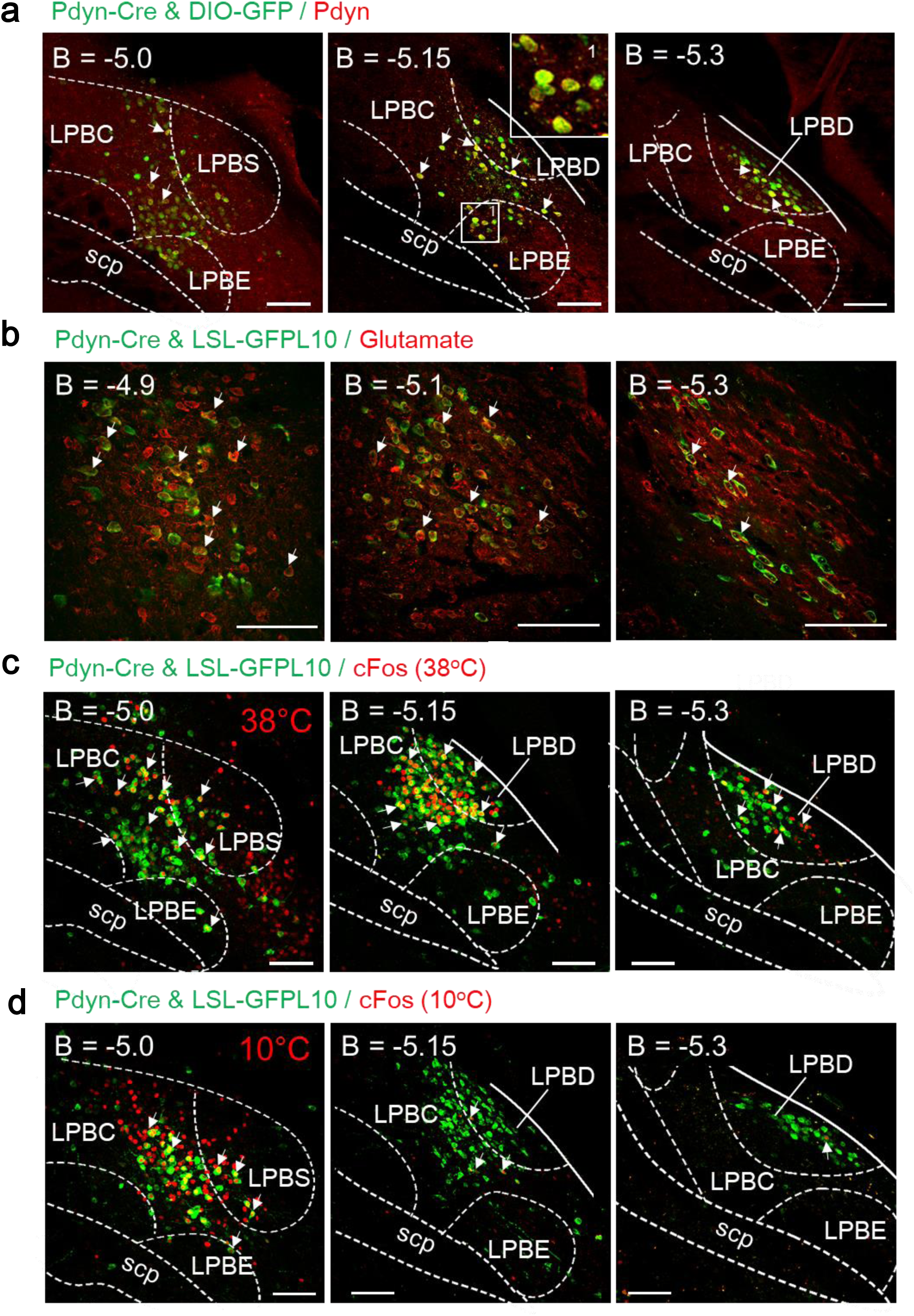
Properties of LPB Pdyn neurons, related to Fig. 2. **a**, The overlapping between Pdyn-Cre & DIO-GFP and Pdyn immune activity in the LPB. The merge ratio was ∼85 % (181 merged / 207 Pdyn-Cre & DIO-GFP^+^ cells = 87.44 %, from n = 3 mice). **b** The overlapping between Pdyn-Cre & LSL-GFPL10 and glutamate immune activity in the LPB. The merge ratio was ∼90 % of Pdyn-Cre & LSL-GFPL10 cells (133 merged / 144 Pdyn-Cre & LSL-GFPL10^+^ cells = 92.4 %, from n = 3 mice). The number of merged cells was ∼8 % of glutamate+ cells (133 merged / 1598 Vglut2^+^ cells = 8.3 %, from n = 3 mice). **c**, The overlapping between warm-induced cFos and Pdyn-Cre & LSL-GFPL10 in the LPB. The merge ratio was ∼40 % of cFos^+^ cells (172 merged / 388 cFos^+^ cells = 44.3 %, from n = 3 mice). The number of merged cells was ∼50 % of Pdyn-Cre & LSL-GFPL10 cells (172 merged / 329 Pdyn-Cre & LSL-GFPL10^+^ cells = 52.3 %, from n = 3 mice). **d**, The overlapping between cold-induced cFos and Pdyn-Cre & LSL-GFPL10 in the LPB. The merge ratio was ∼20 % of cFos^+^ cells (43 merged / 212 cFos^+^ cells = 20.3 %, from n = 3 mice). The number of merged cells was ∼15 % of Pdyn-Cre & LSL-GFPL10 cells (43 merged / 299 Pdyn-Cre & LSL-GFPL10^+^ cells = 14.4 %, from n = 3 mice). Scale bars, 100 µm. Arrows, examples of merged cells. B, bregma; LPB, lateral parabrachial nucleus; LPBC, lateral parabrachial nucleus, central part; LPBD, lateral parabrachial nucleus, dorsal part; LPBE, lateral parabrachial nucleus, external part; LPBS, lateral parabrachial nucleus, superior part; scp, superior cerebellar peduncle.

**Supplemental Fig. 6.**
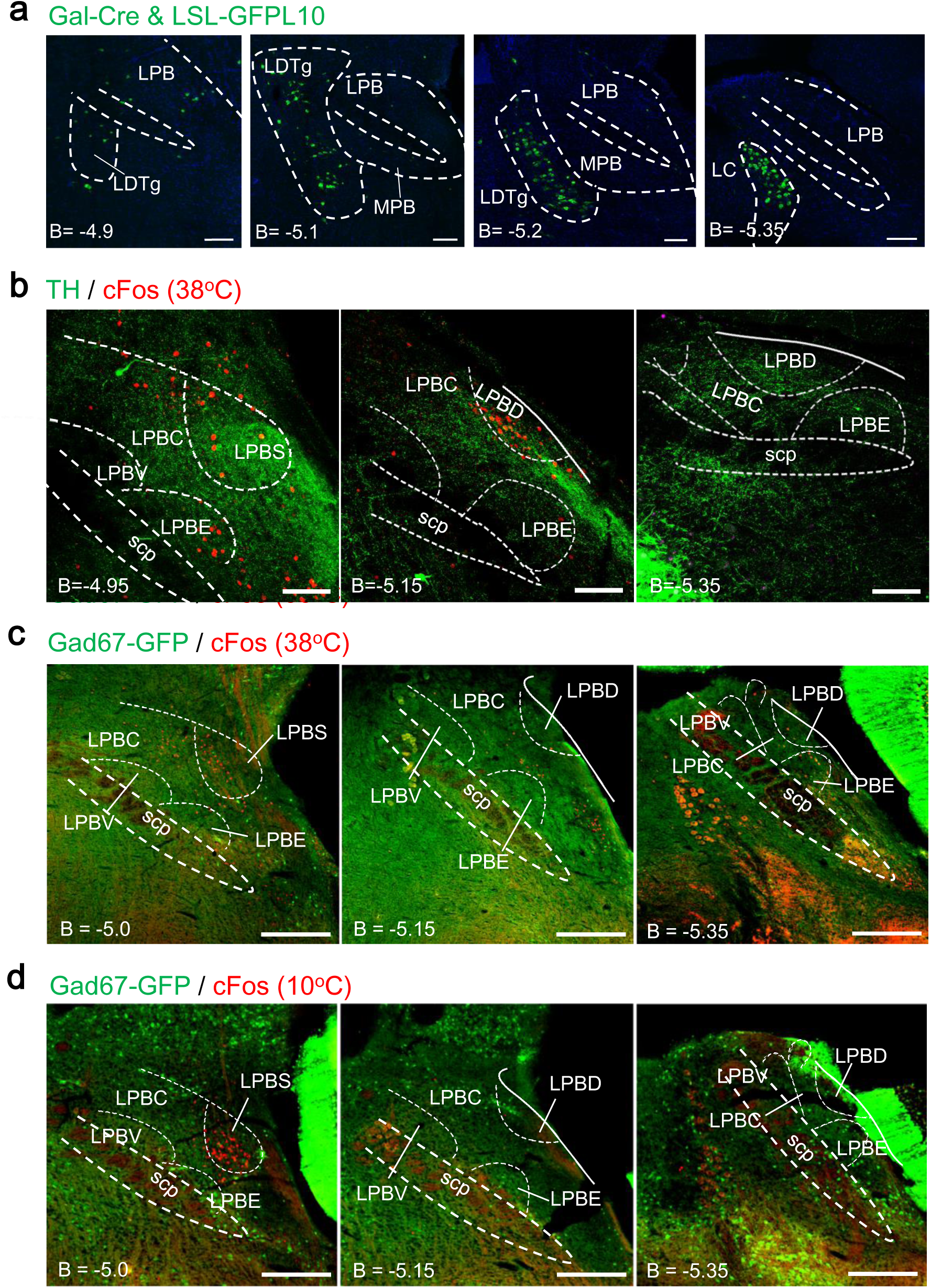
Properties of Galanin-Cre^+^, TH^+^ and GABAergic neurons in the LPB, related to Fig. 2. **a**, GFP expression of Galanin-Cre & LSL-GFPL10 mice in the LPB and the LDTg at different bregma sites. LTDg, laterodorsal tegmental nucleus; MPB, medial parabrachial nucleus. **b**, The staining of warm-induced cFos and TH (tyrosine hydroxylase) in the LPB showed nearly no TH^+^ neuron cell bodies in the LPB. **c**, The overlapping between warm-induced cFos and GABAergic marker Gad67-GFP (Gad67 is also called Gad1.) in the LPB at different bregma sites. The merge ratio was < 1 % (1 merged / 223 cFos^+^ cells = 0.45 %, n = 2). **d**, The overlapping between cold-induced cFos and GABAergic marker Gad67-GFP in the LPB at different bregma sites. The merge ratio was < 1 % (1 merged / 157 cFos^+^ cells = 0.6 %, n = 2). Scale bars, 200 µm.

**Supplemental Fig. 7.**
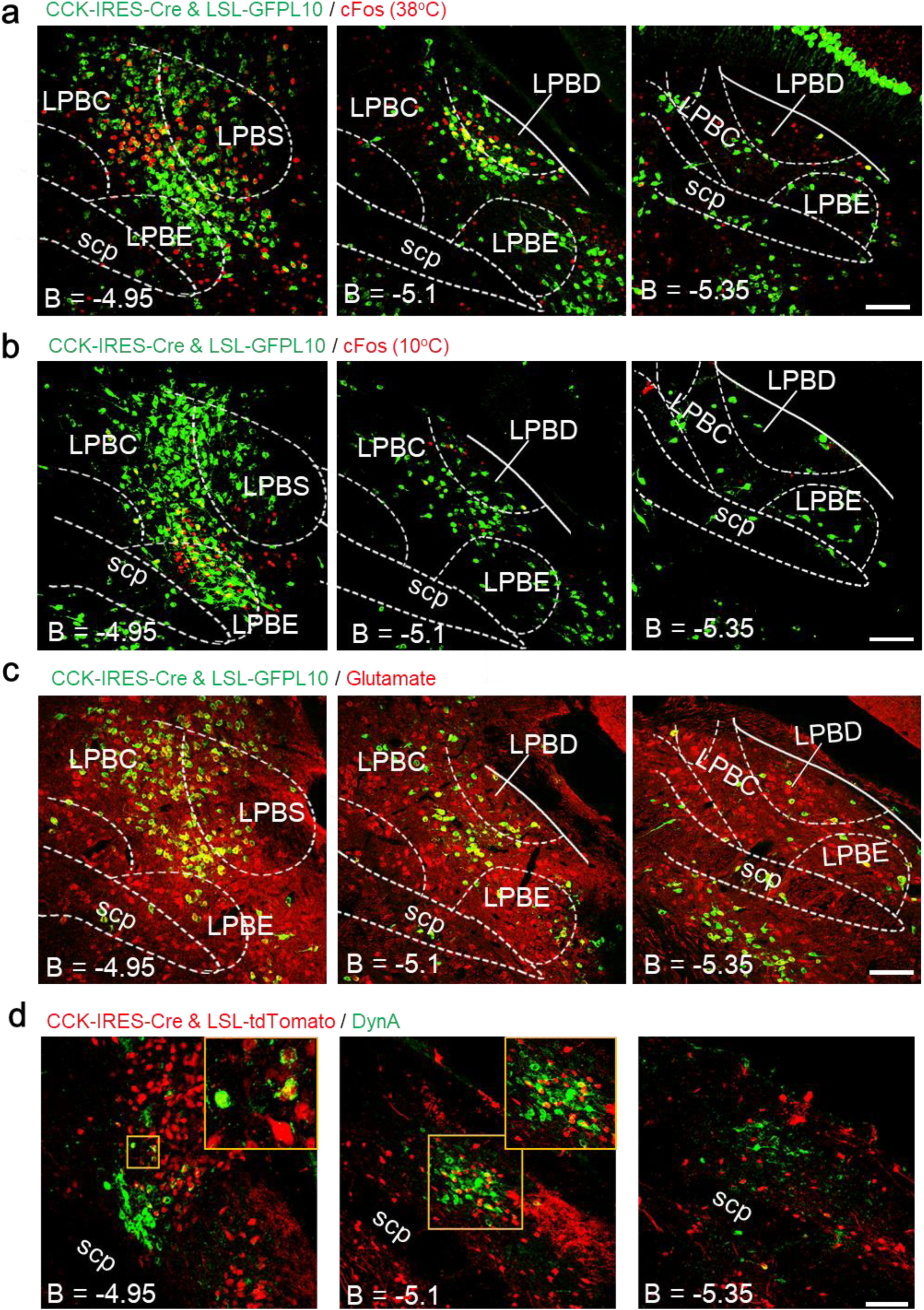
Properties of LPB CCK neurons, related to Fig. 2. **a**, The overlapping between warm-induced cFos and CCK-IRES-Cre & LSL-GFPL10 in the LPB. The merge ratio was ∼50 % of cFos+ cells (147 merged / 293 cFos^+^ cells = 50.2 %, from n = 3 mice). The number of merged cells was ∼20 % of CCK-Cre & LSL-GFPL10 cells (147 merged / 740 CCK-Cre & LSL-GFPL10^+^ cells = 19.9 %, from n = 3 mice). **b**, The overlapping between cold-induced cFos and CCK-Cre & LSL-GFPL10 in the LPB. The merge ratio was ∼20 % of cFos^+^ cells (49 merged / 207 cFos^+^ cells = 23.6 %, from n = 3 mice). The number of merged cells was ∼6 % of CCK-Cre & LSL-GFPL10 cells (49 merged / 855 CCK-Cre & LSL-GFPL10^+^ cells = 5.7 %, from n = 3 mice). **c**, The overlapping between CCK-Cre & LSL-GFPL10 and glutamate immune activity in the LPB. The merge ratio was ∼90 % of CCK-Cre & LSL-GFPL10 cells (349 merged / 403 CCK-Cre & LSL-GFPL10^+^ cells = 86.6 %, from n = 3 mice). **d**, The overlapping between CCK-Cre & LSL-tdTomato (Ai14) and DynA immune activity in the LPB. The merge ratio was ∼7 % of CCK-Cre & LSL-tdTomato cells (17 merged / 242 CCK-Cre & LSL-tdTomato^+^ cells = 7.0 %, from n = 3 mice). The number of merged cells was ∼10 % of DynA+ cells (17 merged / 162 DynA^+^ cells = 10.5 %, from n = 3 mice). Scale bars, 100 µm. B, bregma; LPB, lateral parabrachial nucleus; LPBC, lateral parabrachial nucleus, central part; LPBD, lateral parabrachial nucleus, dorsal part; LPBE, lateral parabrachial nucleus, external part; LPBS, lateral parabrachial nucleus, superior part; scp, superior cerebellar peduncle.

**Supplemental Fig. 8.**
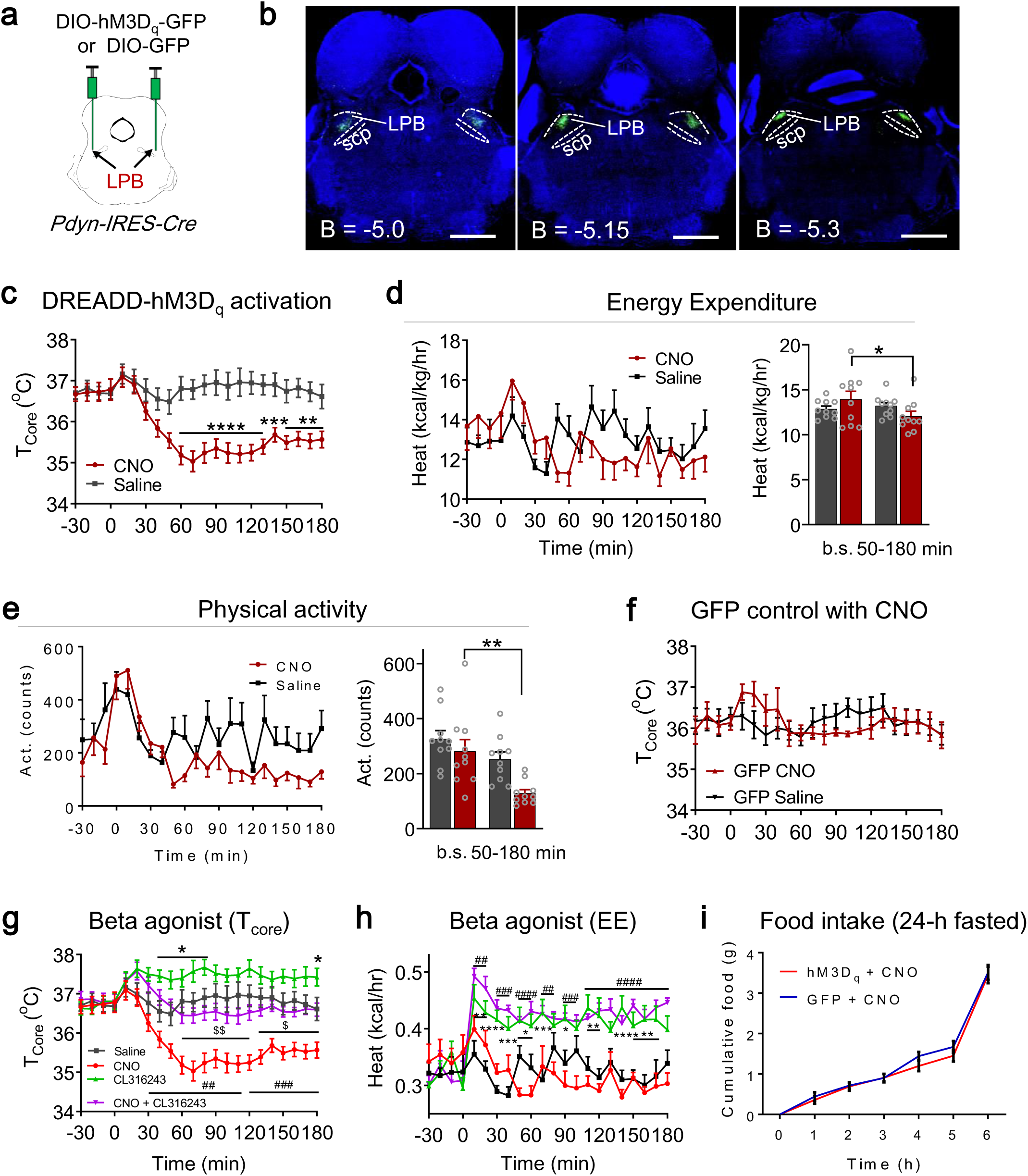
Phenotypes associated with DREADD activation of LPB^Pdyn^ neurons, related to Fig. 2. **a**, Scheme for DREADD activation of LPB^Pdyn & hM3Dq^ neurons. **b**, Viral expression of DIO-hM3D_q_ in LPB^Pdyn^ neurons. **c-e**, DREADD activation induced the reduction of T_core_ (**c**), energy expenditure (**d**) and physical activity (**e**). 2.5 mg/kg CNO was i.p. injected (n = 10 mice). **f**, CNO injection into control mice expressing GFP only did not change T_core_. **g**, β3 adrenergic receptor agonist CL316243 (1 mg/kg, i.p.) suppressed the T_core_ reduction induced by DREADD activation (2.5 mg/kg CNO, i.p.) (Saline, CNO, CL316243+CNO, n = 10 each; CL316243, n = 8). *, CL316243 group compares to Saline group; #, CL316243+CNO group compares to CNO group; $, CL316243+CNO group compares to CL316243 group. **h**, β3 adrenergic receptor agonist CL316243 increased energy expenditure (EE) during DREADD activation (2.5 mg/kg CNO, i.p.) (Saline, CNO, CL316243+CNO, n = 10 each; CL316243, n = 8). **i**, DREADD activation (2.5 mg/kg CNO, i.p.; GFP, n = 7; hM3D_q_, n = 10) did not affect food intake in 24-h fasted animals. All data are shown as mean ± s.e.m. All the p-values are calculated based on repeated measures two-way ANOVA with Bonferroni’s corrections except in (**d-e**) (ordinary two-way ANOVA with Bonferroni’s corrections). *p, ^#^p, ^$^p ≤ 0.05; **p, ^##^p, ^$$^p ≤ 0.01; ***p, ^###^p, ^$$$^p ≤ 0.001; ****p, ^####^p, ^$$$$^p ≤ 0.0001. Scale bars, 1 mm.

**Supplemental Fig. 9.**
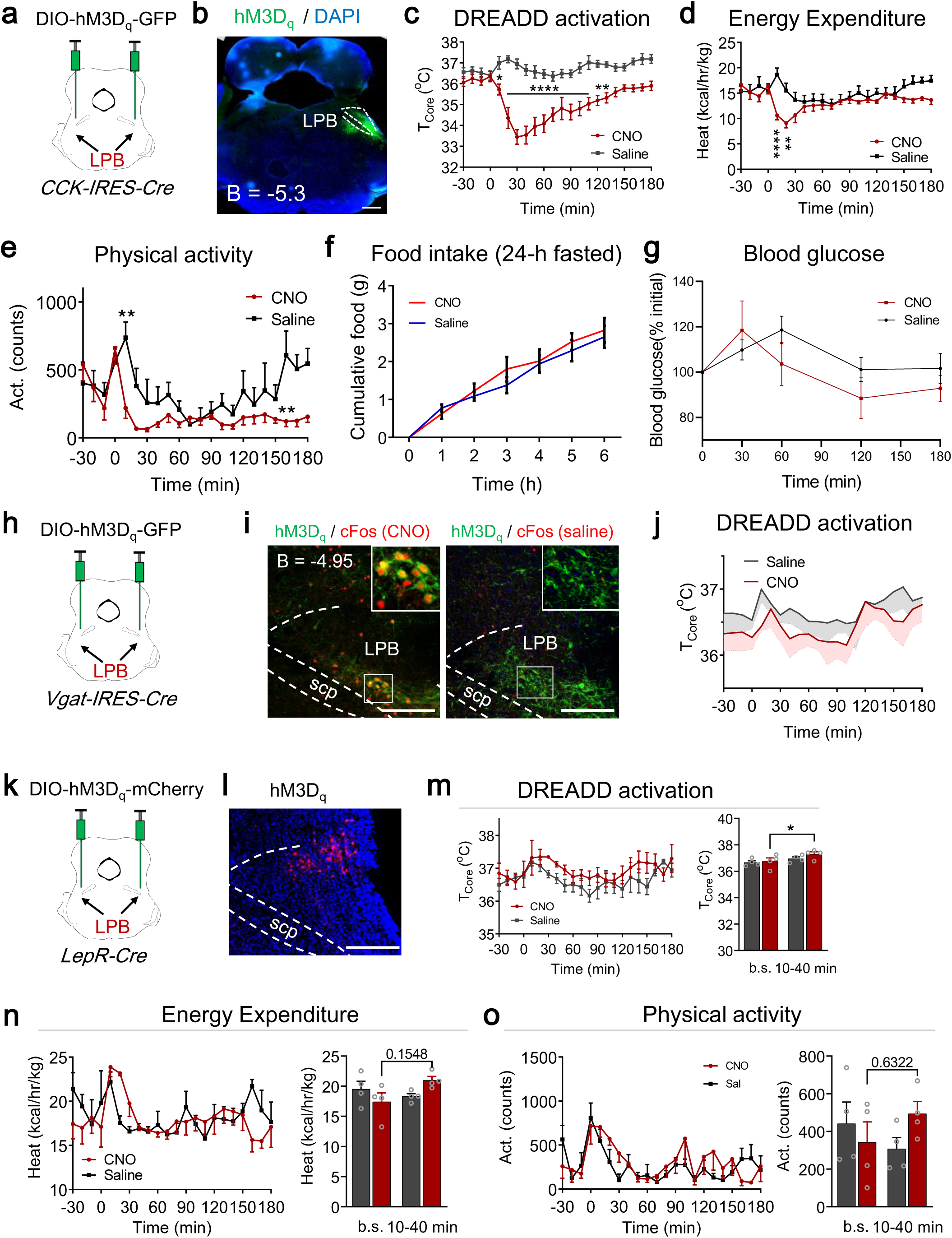
Phenotypes associated with DREADD activation of LPB neuron types, related to Fig. 2. **a**, Scheme for DREADD activation of LPB^CCK & hM3Dq^ neurons. **b**, Viral expression of DIO-hM3D_q_ in LPB^CCK^ neurons. **c-e**, DREADD activation induced the reduction of T_core_ (**c**), energy expenditure (**d**) and physical activity (**e**). 2.5 mg/kg CNO was injected (i.p.; n = 7 mice). **f**, DREADD activation (hM3D_q_, n = 7 mice) did not affect food intake in 24-h fasted animals. **g**, DREADD activation (hM3D_q_, n = 8 mice) slightly yet insignificantly reduced blood glucose level. **h**, Scheme for DREADD activation of LPB^Vgat^ neurons. **i**, Viral expression of DIO-hM3D_q_ in LPB^Vgat^ neurons. **j**, DREADD activation did not affect T_core_ significantly (n = 4 mice). **k**, Scheme for DREADD activation of LPB^LepR^ neurons. **l**, Viral expression of DIO-hM3D_q_ in LPB^LepR^ neurons. **m-o**, DREADD activation induced a slight increase of T_core_ (**m**), energy expenditure (**n**) and physical activity (**o**). 2.5 mg/kg CNO was injected (i.p.; n = 4 mice). All data are shown as mean ± s.e.m. All the p-values are calculated based on repeated measures two-way ANOVA with Bonferroni’s corrections except in (**m-o**) (ordinary two-way ANOVA with Bonferroni’s corrections). *p ≤ 0.05; **p ≤ 0.01; ***p≤ 0.001; ****p ≤ 0.0001; ns, not significant. Scale bars, 500 µm in (**b**), and 200 µm in (**i**) and (**l**).

**Supplemental Fig. 10.**
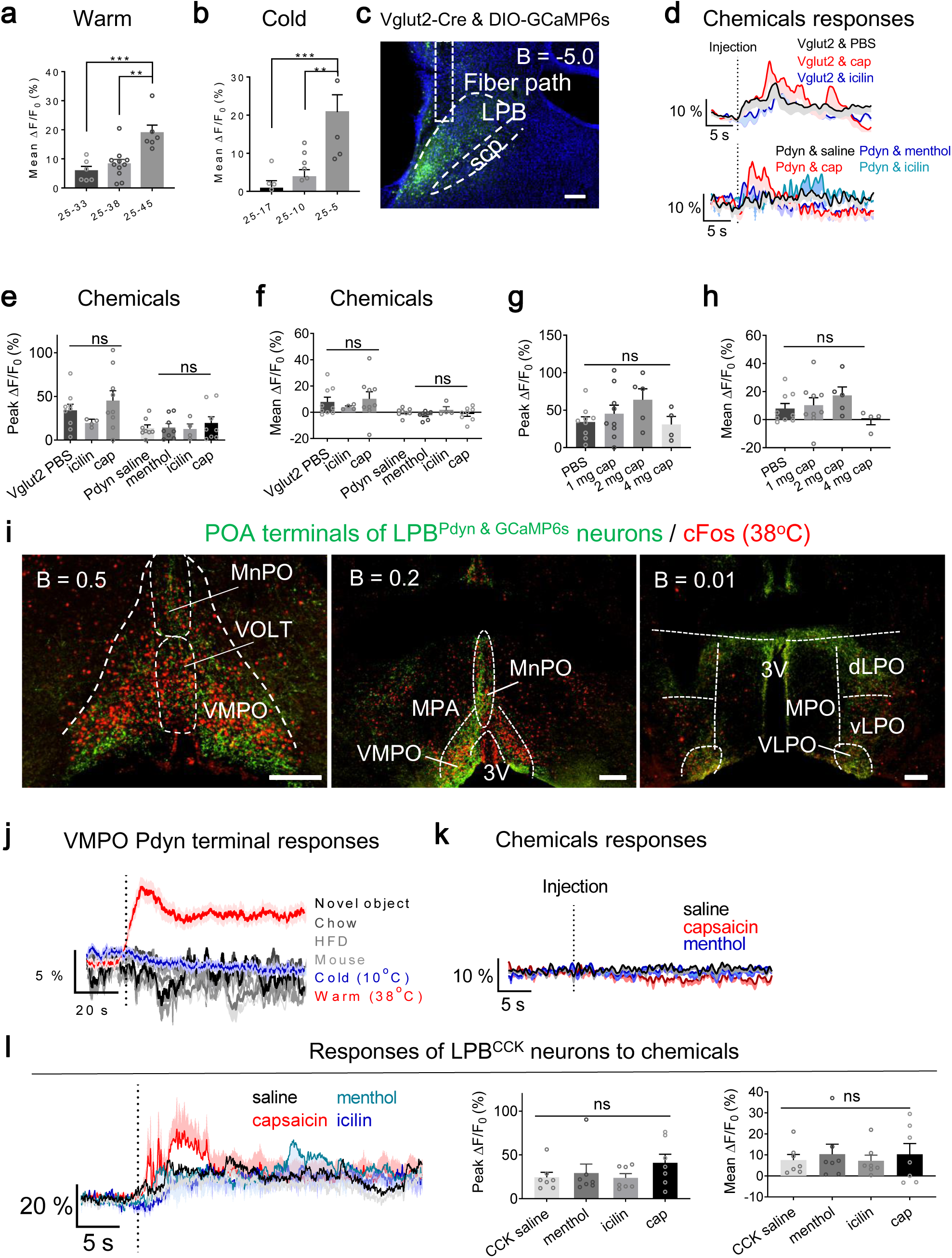
Fiber photometry of neural calcium, related to Fig. 3. **a-b**, Mean responses (t = (25 – 100 s)) to different warm (**a**) or cold (**b**) temperatures in LPB^Pdyn^ neurons (25 – 33 °C, 25 – 45 °C, 25 – 17 °C, 25 – 5 °C, n = 6 each; 25 – 38 °C, n = 11; 25 – 10 °C, n = 7). **c**, Viral expression of DIO-GCaMP6s in the LPB^Vglut2^ neurons. Scale bar, 200 µm. **d**, Calcium dynamics in response to capsaicin (cap, 1 mg/kg body weight, i.p.), menthol (10 mg/kg body weight, i.p.) or icilin (2.5 mg/kg body weight, i.p.) in LPB^Vglut2^ neurons and LPB^Pdyn^ neurons (Vglut2: PBS, n = 9; icilin, n = 4; cap (capsaicin), n = 9; Pdyn: saline, menthol, cap, n = 8 each; icilin, n = 4). **e-f**, The quantification of peak (**e**) and mean (**f**) responses to capsaicin, menthol or icilin in LPB^Vglut2^ neurons and LPB^Pdyn^ neurons. The mean is defined as the average response in 25-sec window after vehicle/chemicals injection. **g-h**, The quantification of peak (**g**) and mean (**h**) responses to different dosages (as indicated) of capsaicin in LPB^Vglut2^ neurons. ΔF/F_0_ represents change in GCaMP6s fluorescence from the mean level (t = (− 5 − 0 s)). **i**, Expression of GCaMP6s from LPB^Pdyn^ neuron terminals in the POA and warm-induced cFos. **j**, Calcium dynamics of LPB^Pdyn^ terminals in the VMPO in response to different stimuli (Novel object, HFD, Mouse, Cold, n = 7 each; Chow, n = 4; Warm, n = 11). ΔF/F_0_ represents change in GCaMP6s fluorescence from the mean level (t = (− 120 − 0 s)). HFD, high fat diet. **k**, Calcium dynamics of LPB^Pdyn^ terminals in the VMPO (n = 6 each) in response to capsaicin (2 mg/kg body weight, i.p.), menthol (10 mg/kg body weight, i.p.). ΔF/F_0_ represents change in GCaMP6s fluorescence from the mean level (t = (− 10 − 0 s)). **l**, Calcium dynamics in response to capsaicin (cap, 1 mg/kg body weight, i.p.), menthol (10 mg/kg body weight, i.p.) or icilin (2.5 mg/kg body weight, i.p.) in LPB^CCK^ neurons (n = 7 each). All data are shown as mean ± s.e.m. All the p-values are calculated based on ordinary one-way ANOVA. *p ≤ 0.05; **p ≤ 0.01; ***p ≤ 0.001; ns, not significant. VLPO, ventrolateral preoptic nucleus; dLPO and vLPO, dorsal and ventral part of lateral preoptic nucleus respectively; MnPO, median preoptic nucleus; OVLT, organum vasculosum laminae terminalis; MPA, medial preoptic area; MPO, medial preoptic nucleus; 3V, 3rd ventricle; VMPO, ventromedial preoptic nucleus.

**Supplemental Fig. 11.**
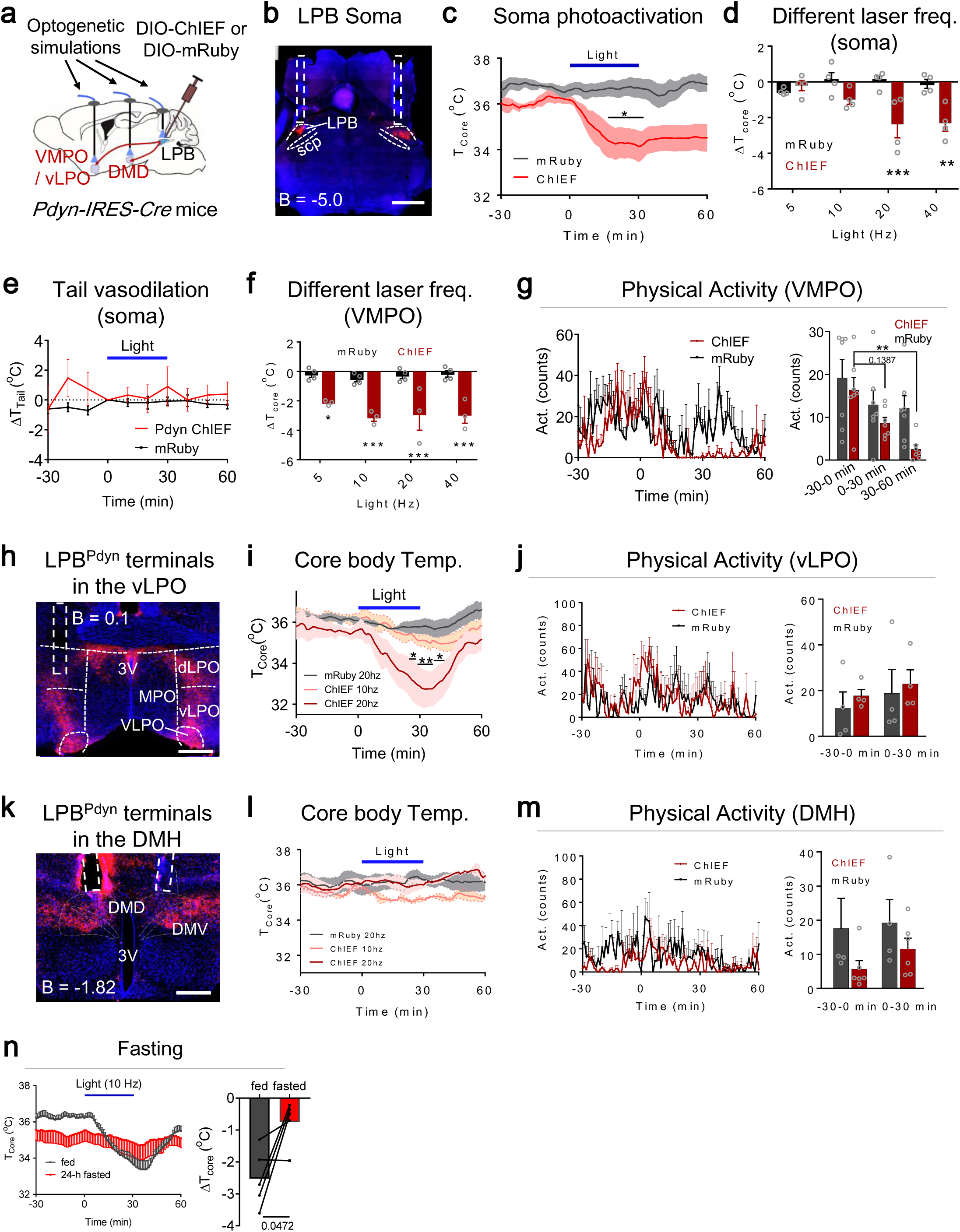
Phenotypes associated with activation of LPB^Pdyn^ projections, related to Fig. 4. **a**, Scheme for optogenetic stimulation LPB^Pdyn & ChIEF^ neural soma and projections at the VMPO, the vLPO and the DMH. **b**, Viral ChIEF expression in LPB^Pdyn^ neurons at the injection site. Scale bar, 1 mm. **c**, Changes of T_core_ after LPB soma photoactivation (Pdyn ChIEF, n = 3; Vglut2 GFP, n = 4). Laser pattern: 473 nm, 3 mW, 20 Hz, 30 min. **d**, ΔT_core_ after LPB soma photoactivation (ChIEF, n = 3; mRuby, n = 4) under different laser frequencies. Laser pattern: 473 nm, 6 mW, Hz as indicated, 30 min. **e**, No changes of tail temperatures after LPB soma photoactivation (Pdyn ChIEF, n = 3; Vglut2 GFP, n = 4). Laser pattern: 473 nm, 3 mW, 20 Hz, 30 min. **f**, ΔT_core_ after photoactivation of LPB^Pdyn^ neuron terminals in the VMPO (ChIEF, n = 3; mRuby, n = 4) under different laser frequencies. Laser pattern: 473 nm, 6 mW, Hz as indicated, 30 min. **g**, Changes of physical activity after photoactivation of LPB^Pdyn^ neuron terminals in the VMPO (n = 7 each). **h**, ChIEF expression in LPB^Pdyn^ neuron terminals in the vLPO. Scale bar, 500 μm. **i-j**, Changes of T_core_ (**i**) and physical activity (**j**) after photoactivation of LPB^Pdyn^ neuron terminals in the vLPO (n = 4 each). **k**, ChIEF expression in LPB^Pdyn^ neuron terminals in the DMH. Scale bar, 500 μm. **l-m**, Changes of T_core_ (**l**) and physical activity (**m**) after photoactivation of LPB^Pdyn^ neuron terminals in the DMH (n = 4 each). Laser pattern: 473 nm, 6 mW, 20 Hz, 30 min. **n**, Changes of T_core_ after photoactivation of LPB^Pdyn^ neuron terminals in the VMPO after 24-h fasting (n = 5). Laser pattern: 473 nm, 6 mW, 10 Hz, 30 min. All data are shown as mean ± s.e.m. The p-values are calculated based on ordinary two-way ANOVA with Bonferroni’s corrections (**d**; right panels in **g, j, m**), or repeated measures two-way ANOVA with Bonferroni’s corrections (**c, e, i, l**) or paired t test (right panel in **n**). *p ≤ 0.05; **p ≤ 0.01; ***p ≤ 0.001; ns, not significant.

**Supplemental Fig. 12.**
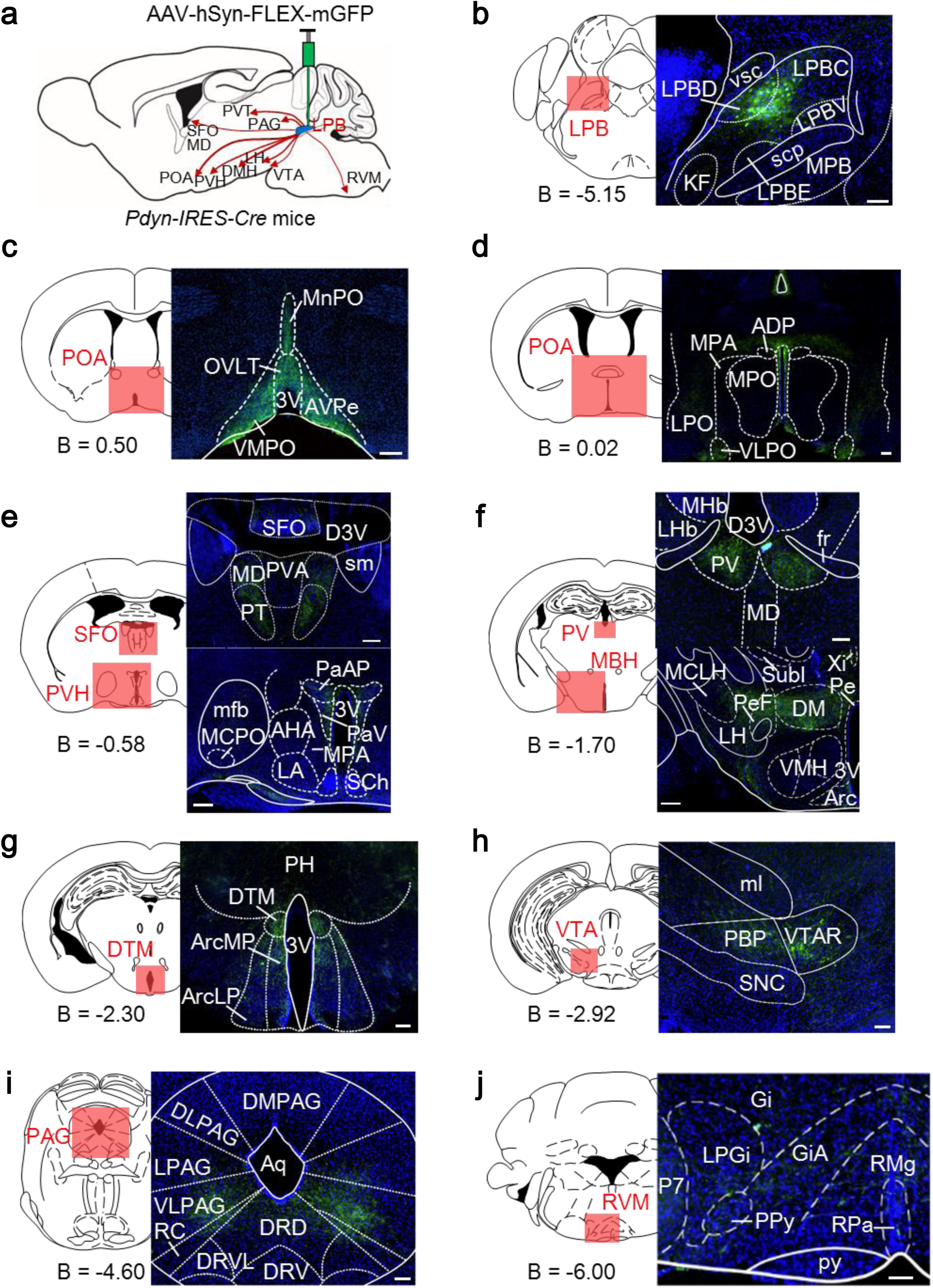
Whole-brain projections of LPB Pdyn neurons, related to Fig. 4. **a**, Summary of the projection pattern. AAV-hSyn-FLEX-mGFP was injected to the LPB of Pdyn-Cre mice. **b**, Viral expression of FLEX-mGFP (membrane-bound GFP) in the LPB^Pdyn^ neurons. **c-j**, The projection sites with high terminal fluorescence. Scale bars, 100 µm. B, bregma; LPB, lateral parabrachial nucleus; PAG, periaqueductal gray matter; PVT, paraventricular thalamic nucleus; RVM, rostral ventromedial medulla; SFO, subfornical organ; MD, mediodorsal thalamic nucleus; POA, preoptic area; PVH, paraventricular hypothalamic nucleus; DMH, dorsomedial hypothalamic nucleus; LH, lateral hypothalamic area; VTA, ventral tegmental area;; LPBC, lateral parabrachial nucleus, central part; LPBD, lateral parabrachial nucleus, dorsal part; LPBE, lateral parabrachial nucleus, external part; LPBV, lateral parabrachial nucleus, ventral part; scp, superior cerebellar peduncle; vsc, ventral spinocerebellar tract; KF, Klliker-Fuse nucleus; MPB, medial parabrachial nucleus; VMPO, ventromedial preoptic nucleus; 3V, 3rd ventricle; MnPO, median preoptic nucleus; AVPe, anteroventral periventricular nucleus; OVLT, organum vasculosum laminae terminalis; VLPO, ventrolateral preoptic nucleus; LPO, lateral preoptic area; MPA, medial preoptic area; MPO, medial preoptic nucleus; ADP, anterodorsal preoptic nucleus; D3V, dorsal 3rd ventricle; PVA, paraventricular thalamic nucleus, anterior part; PT, paratenial thalamic nucleus; sm, stria medullaris; PaAP, paraventricular hypothalamic nucleus, anterior parvic; mfb, medial forebrain bundle; MCPO, magnocellular preoptic nucleus; AHA, anterior hypothalamic area, anterior part; LA, lateroanterior hypothalamic nucleus; PaV, araventricular hypothalamic nucleus, ventral part; SCh, suprachiasmatic nucleus; MHb, medial habenular nucleus; LHb, lateral habenular nucleus; PV, paraventricular thalamic nucleus; fr, fasciculus retroflexus; MCLH, magnocellular nucleus of the lateral hypothalamus; Subl, subincertal nucleus; Xi, xiphoid thalamic nucleus; Pe, periventricular hypothalamic nucleus; PeF, perifornical nucleus; MBH, mediobasal hypothalamus; VMH, ventromedial hypothalamic nucleus; Arc, arcuate hypothalamic nucleus; DM, dorsomedial hypothalamic nucleus; PH, posterior hypothalamic nucleus; DTM, dorsal tuberomammillary nucleus; ArcMP, arcuate hypothalamic nucleus, medial posterior part; ArcLP, arcuate hypothalamic nucleus, lateroposterior part; PBP, parabrachial pigmented nucleus of the VTA; VTAR, ventral tegmental area, rostral part; SNC, substantia nigra, compact part; ml, medial lemniscus; DMPAG, dorsomedial periaqueductal gray; LPAG, lateral periaqueductal gray; DLPAG, dorsolateral periaqueductal gray; VLPAG, ventrolateral periaqueductal gray; RC, raphe cap; DRVL, dorsal raphe nucleus, ventrolateral part; DRV, dorsal raphe nucleus, ventral part; DRD, dorsal raphe nucleus, dorsal part; Aq, aqueduct; Gi, gigantocellular reticular nucleus; GiA, gigantocellular reticular nucleus, alpha part; ; LPGi, lateral paragigantocellular nucleus; RMg, raphe magnus nucleus; P7, perifacial zone; Ppy, parapyramidal nucleus; py, pyramidal tract; Rpa, raphe pallidus nucleus.

**Supplemental Fig. 13.**
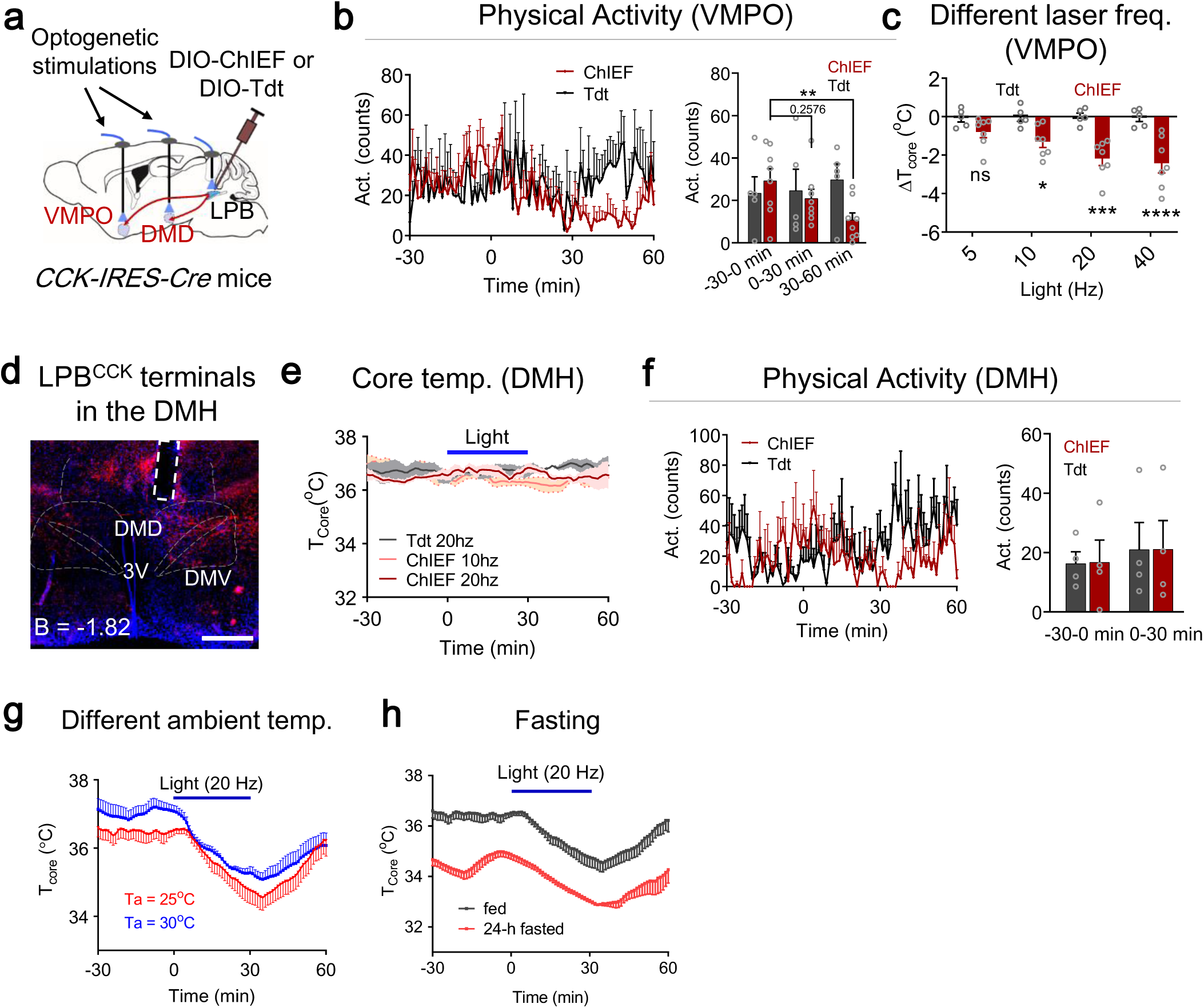
Phenotypes associated with activation of LPB^CCK^→POA/DMH projections, related to Fig. 4. **a**, Scheme for optogenetic stimulation LPB^CCK & ChIEF^ neural projections at the VMPO and the DMH. **b**, Changes of physical activity after photoactivation of LPB^CCK^ neuron terminals in the VMPO (ChIEF, n = 8; tdTomato, n = 5). **c**, ΔT_core_ after photoactivation of LPB^CCK^ neuron terminals in the VMPO (ChIEF, n = 7; tdTomato, n = 5) under different laser frequencies. Laser pattern: 473 nm, 3 mW, Hz as indicated, 30 min. **d**, ChIEF expression in LPB^CCK^ neuron terminals in the DMH. Scale bar, 500 μm. **e-f**, Changes of T_core_ (**e**) and physical activity (**f**) after photoactivation of LPB^CCK^ neuron terminals in the DMH (n = 4 each). Laser pattern: 473 nm, 6 mW, 20 Hz, 30 min. **g-h**, Changes of T_core_ after photoactivation of LPB^cck^ neuron terminals in the VMPO (n = 5 each) under different ambient temperatures (**g**) or after fasting (**h**). Behavioral tests were performed at T_a_ = 25 °C unless specified. Laser pattern: 473 nm, 3 mW, 20 Hz, 30 min. All data are shown as mean ± s.e.m. The p-values are calculated based on ordinary two-way ANOVA with Bonferroni’s corrections (**c**; right panel in **f)** or repeated measures two-way ANOVA with Bonferroni’s corrections (right panel in **b**). *p ≤ 0.05; **p ≤ 0.01; ***p ≤ 0.001; ****p ≤ 0.001; ns, not significant.

**Supplemental Fig. 14.**
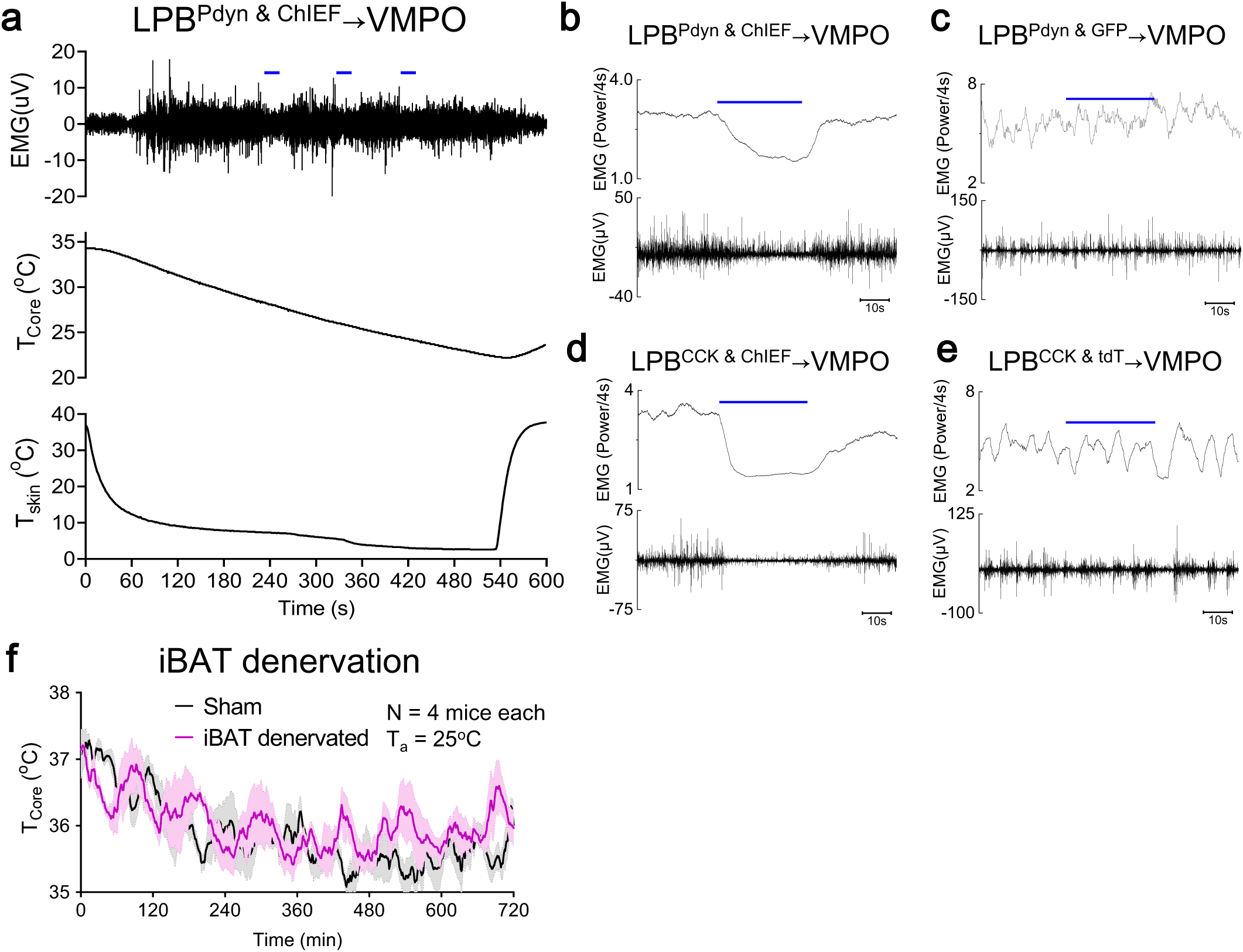
Shivering activity recording and iBAT denervation, related to Fig. 5. **a**, A representative EMG trace of cold-induced muscle shivering. Blue bars represent laser stimulations. Electromyography (EMG) was recorded from the nuchal muscle after anesthesia. T_core_ and T_skin_ were monitored simultaneously. Laser pattern: 473 nm, 6 mW, 20 Hz, 30 s. **b-c**, Example traces for terminal photoactivation of LPB^Pdyn & ChIEF^→VMPO circuitry (**b**) and their controls (**c**). **d-e**, Examples traces for terminal photoactivation of LPB^CCK & ChIEF^→VMPO circuitry (**c**) and their controls (**d**). **f**, Basal T_core_ of sham and iBAT denervated LPB^Pdyn & ChIEF^ mice under room temperature. All data are shown as mean ± s.e.m.

**Supplemental Fig. 15.**
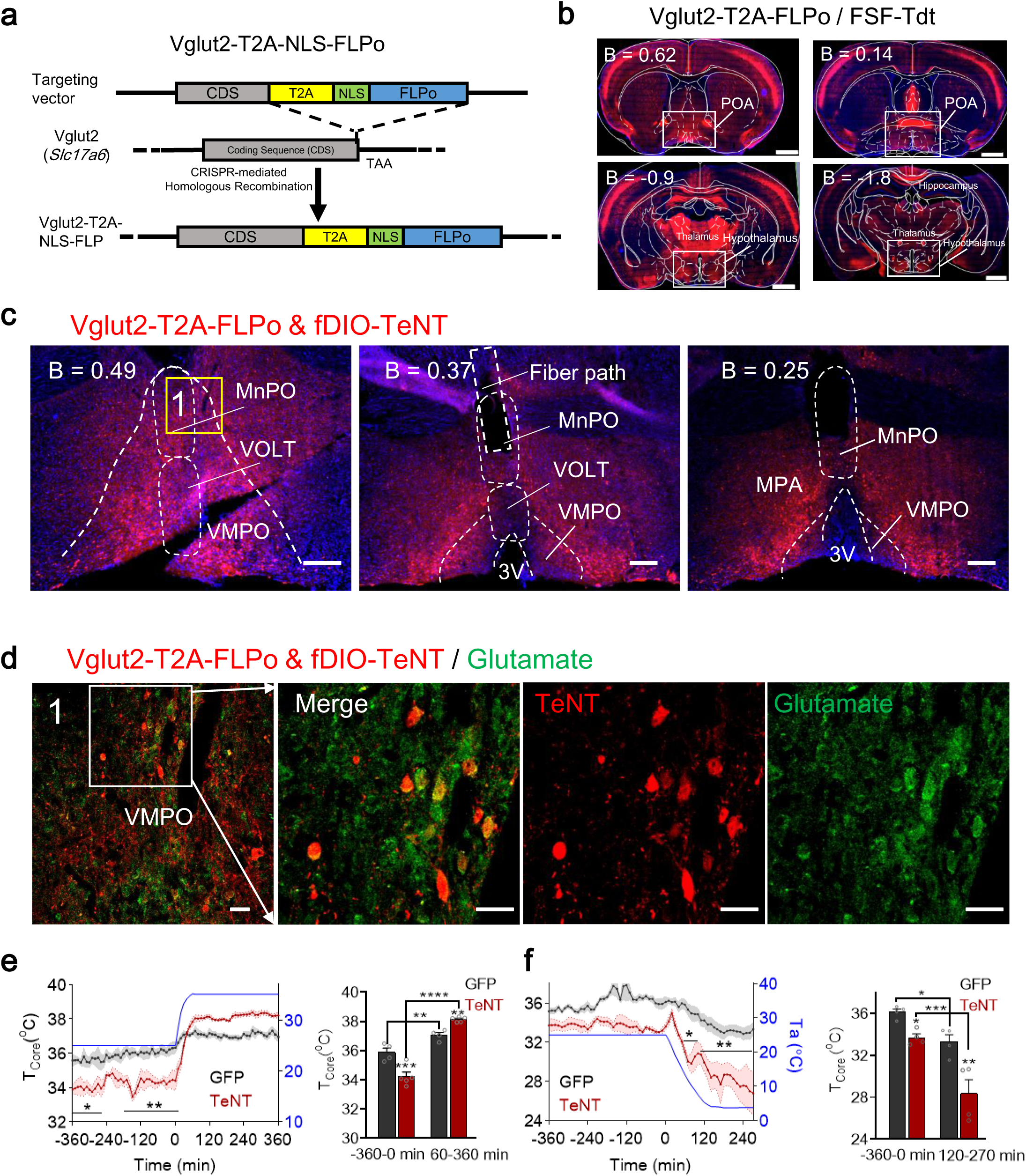
Characterization of Vglut2-T2A-FLPo mice, related to Fig. 5. **a**, Scheme for knocking-in the *NLS-T2A-FLPo* sequence into the *Vglut2* genomic locus using CRISPR/Cas9-mediated gene editing. The *NLS-T2A-FLPo* coding sequence was inserted right before the stop codon. **b**, The representative expression of Vglut2-T2A-FLPo & FSF-Tdt in the hypothalamus and the cortex. Scale bars, 1 mm. **c**, The representative expression of Vglut2-T2A-FLPo & fDIO-TeNT-mCherry in the POA. Scale bars, 200 µm. **d**, The overlapping of glutamate immune activity and Vglut2-T2A-FLPo & fDIO-TeNT-mCherry in the POA. The merged cells are ∼96% of glutamate neurons (194 merged / 201 glutamate+ = 96.5%, n = 2 mice). Scale bars, 50 µm. **e-f**, Blocking POA glutamatergic neurons by Vglut2-T2A-FLPo & fDIO-TeNT reduced the baseline T_core_ in RT (∼26°C) and impaired thermoregulation in both warm (**e**) and cold (**f**) (GFP, n = 4; TeNT, n = 5). All data are shown as mean ± s.e.m. The p-values are calculated based on repeated measure two-way ANOVA with Bonferroni’s corrections (left panels in **e** and **f**), or ordinary two-way ANOVA with Bonferroni’s corrections (right panels in **e** and **f**). *p ≤ 0.05; **p ≤ 0.01; ***p ≤ 0.001.

**Supplemental Fig. 16.**
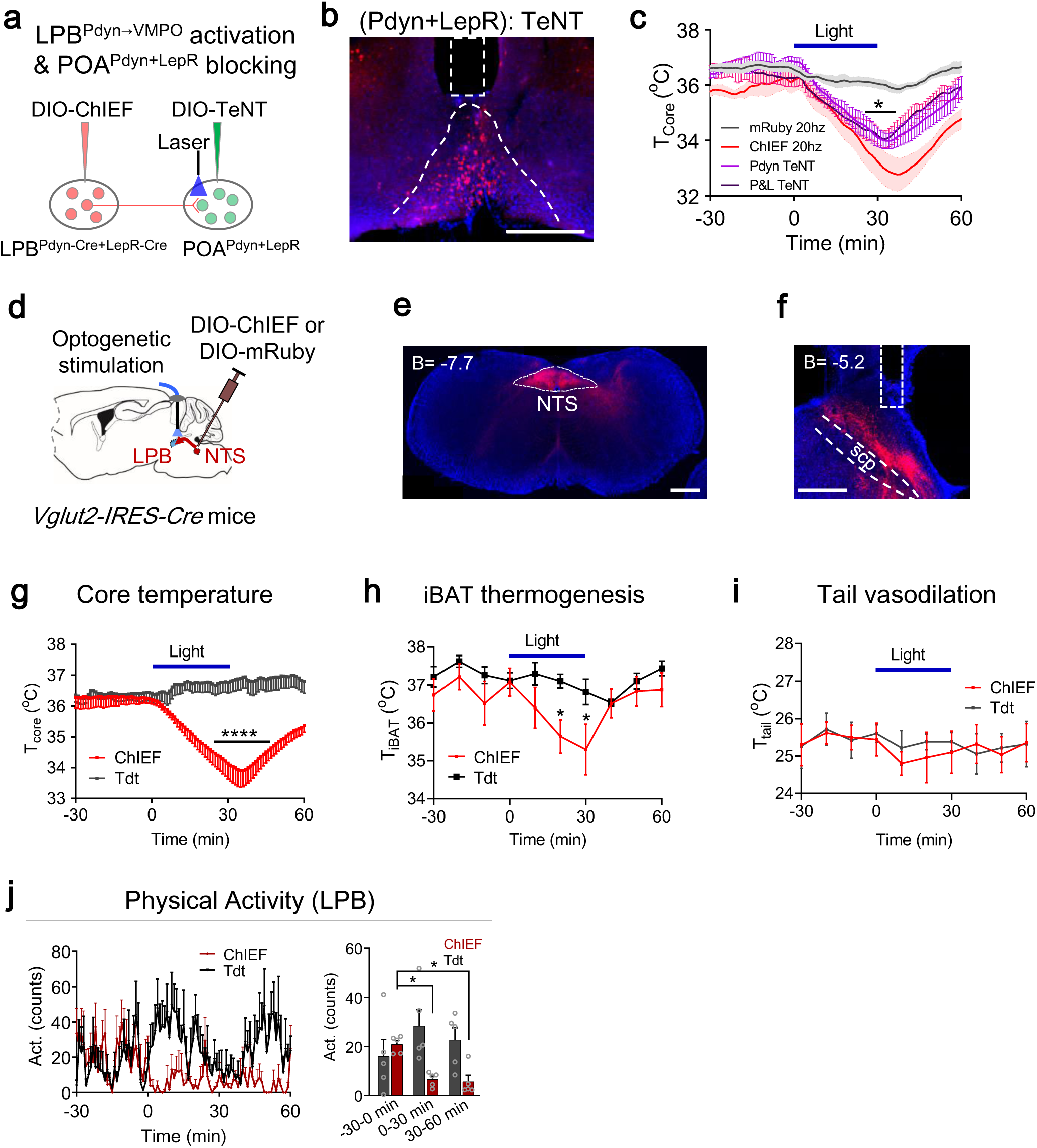
Role of POA LepR neurons in LPB^Pdyn^→POA circuitry, and role of NTS^Vglut2^→LPB projection in tail vasodilation, related to Fig. 5. **a**, Scheme to block POA LepR / Pdyn neurons while photoactivation of VMPO terminals from LPB^Pdyn^ neurons. **b**, The representative expression of (LepR & Pdyn–Cre) & DIO-TeNT-mCherry in the POA. **c**, Blocking of POA Pdyn neurons or (LepR + Pdyn) neurons could not significantly affect the hypothermia induced by activation of LPB^Pdyn^→VMPO circuitry (ChIEF, n = 5; mRuby, n = 5; Pdyn blocking, n = 4; LepR + Pdyn blocking, n = 4). Laser pattern: 473 nm, 6 mW, 20 Hz, 30 min. **d**, Scheme for optogenetic stimulation the NTS^Vglut2^→LPB projection. **e**, ChIEF expression in the NTS^Vglut2^ neurons. **f**, ChIEF expression of NTS^Vglut2^ neuron terminals in the LPB. **g-j**, Changes of T_core_ (**g**), T_iBAT_ (**h**), T_tail_ (**i**) and physical activity (**j**) after photoactivation of NTS^Vglut2^ neuron terminals in the LPB (n = 5 each). Laser pattern: 473 nm, 6 mW, 20 Hz, 30 min. All data are shown as mean ± s.e.m. All the p-values are calculated based on repeated measures two-way ANOVA with Bonferroni’s corrections. *p ≤ 0.05; ****p ≤ 0.0001; ns, not significant.

**Supplemental Fig. 17.**
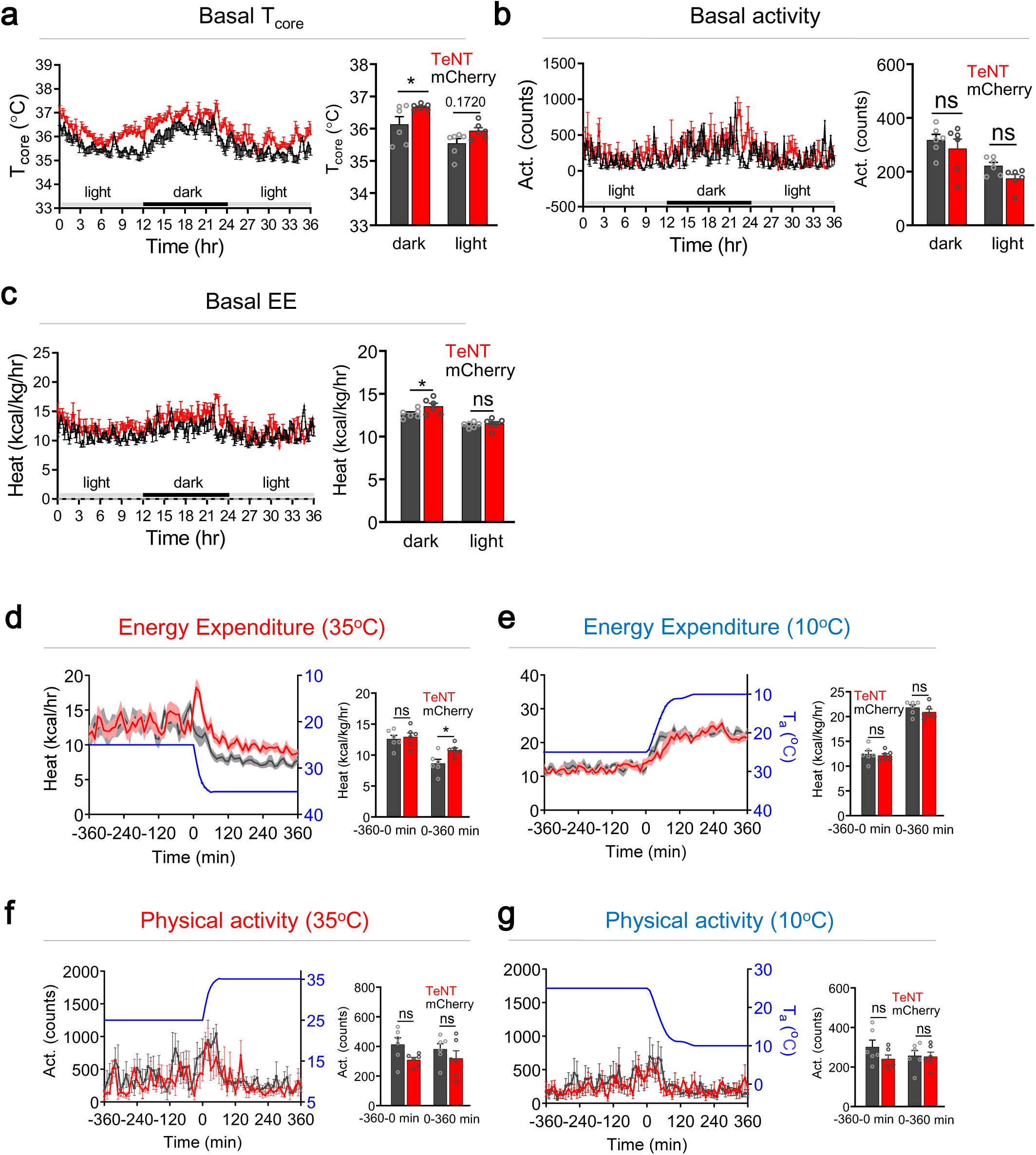
Requirement of LPB^Pdyn→VMPO^ circuitry in heat defense and fever limiting, related to Fig. 6. **a-c**, Changes in basal T_core_ (**a**), basal physical activity (**b**), and basal energy expenditure (**c**) after blocking of POA-projected LPB^Pdyn^ neurons (n = 6 each; 4-6 weeks post viral injection). **d-g**, Changes of energy expenditure (**d-e**), and physical activity (**f-g**) during warm challenge (**d, f**) or cold challenge (**e, g**), respectively, after blocking of POA-projected LPB^Pdyn^ neurons (n = 6 each; 6-8 weeks post viral injection). (– 360 – 0 min) and (0 – 360 min) represent the averaged EE between t = (– 360 – 0 min) and t = (0 – 360 min), respectively. All data are shown as mean ± s.e.m. All the p-values are calculated based on ordinary two-way ANOVA with Bonferroni’s corrections (right panels in **a-g**). *p ≤ 0.05; ns, not significant.

**Supplemental Fig. 18.**
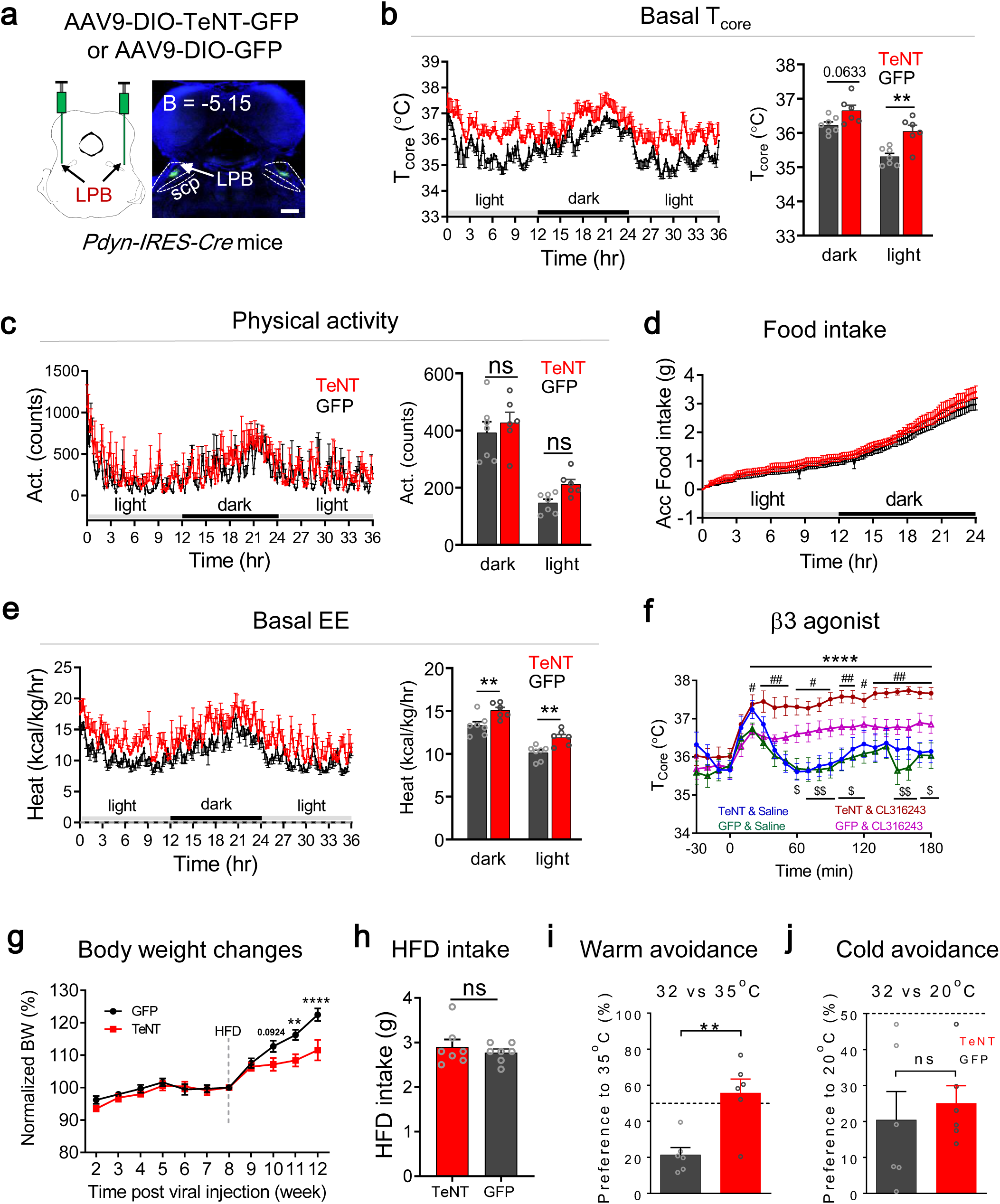
Requirement of LPB^Pdyn^ neurons in energy expenditure and thermal choice, related to Fig. 6. **a**, Scheme to block LPB^Pdyn^ neurons with neurotoxin TeNT and the viral expression. Scale bar, 500 µm. **b-e**, Changes in basal T_core_ (**b**), basal physical activity (**c**), accumulative food intake (**d**), and basal energy expenditure (**e**) after neural blocking (TeNT, n = 6; GFP, n = 7 ; 4-6 weeks post viral injection). **f**, Changes of T_core_ in response to injection of β3 agonist CL316243 (TeNT, n = 10 each; GFP, n = 8 each; 4-8 weeks post viral injection). *, TeNT & CL316243 group compares to TeNT & Saline group; #, TeNT & CL316243 group compares to GFP & CL316243 group; $, GFP & CL316243 group compares to GFP & Saline group. **j**, Changes of normalized body weight after neural blocking (n = 7 each) under chow (0-8 weeks) and high fat diet (HFD, 8-12 weeks). **h**, Averaged daily food intake under high fat diet (n = 7 each, 11 week). **i-j**, Changes of temperature preference to warm (**i**) or cold (**j**) temperatures, respectively, after neural blocking (n = 6 each). All data are shown as mean ± s.e.m. The p-values are calculated based on ordinary two-way ANOVA with Bonferroni’s corrections (right panels in **b, c, e)** or repeated measures two-way ANOVA with Bonferroni’s corrections (**f, g**), or unpaired t test (**h-j**). *^,#^p ≤ 0.05; **^,##,$$^ p ≤ 0.01; ***^,###,$$$^ p ≤ 0.001; **** p ≤ 0.0001; ns, not significant.

**Supplemental Fig. 19.**
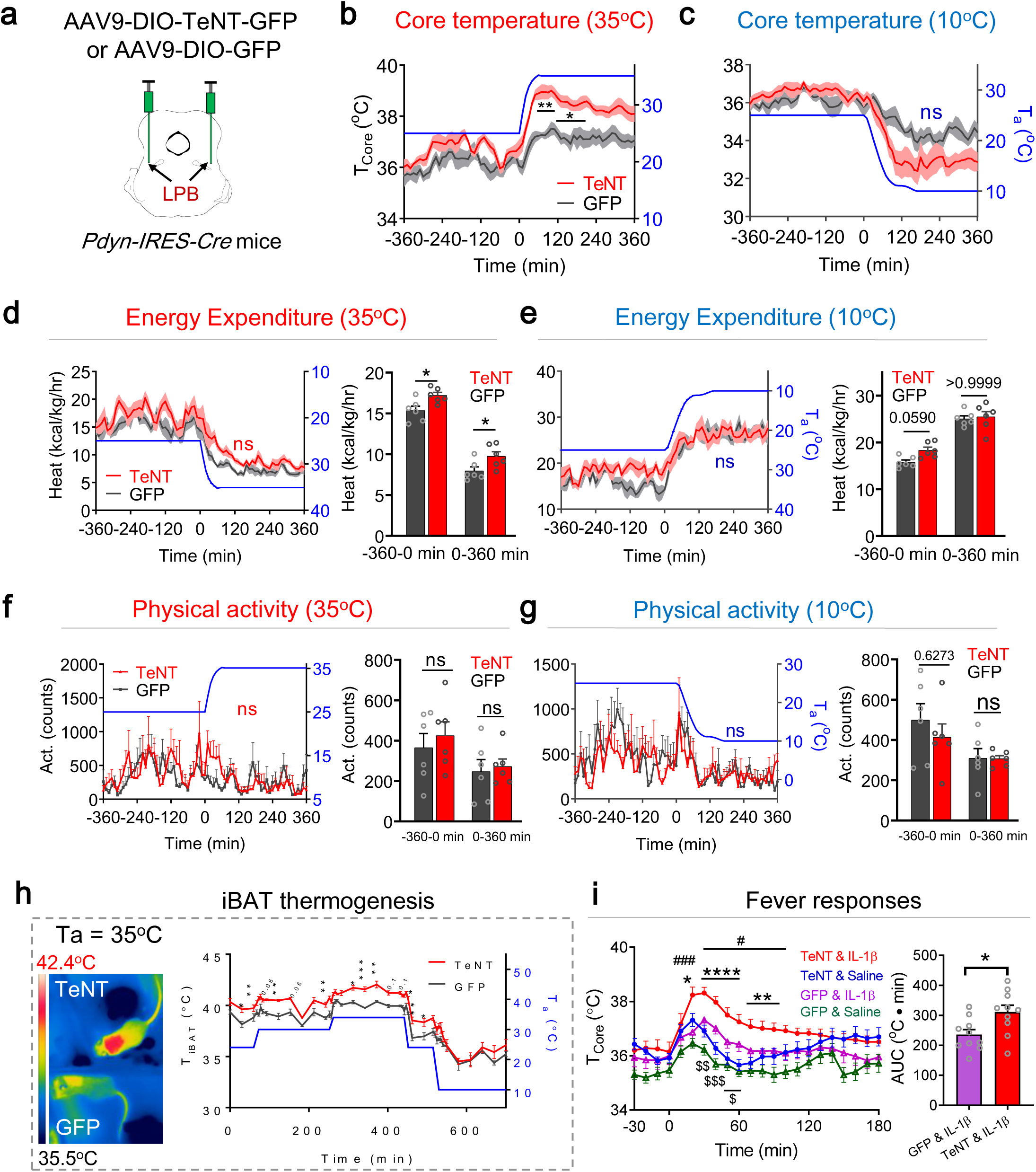
Requirement of LPB^Pdyn^ neurons in heat defense, iBAT thermogenesis and fever limiting, related to Fig. 6. **a**, Scheme to block LPB^Pdyn^ neurons with TeNT. **b-e**, Changes of T_core_ (**b-c**), energy expenditure (**d-e**), and physical activity (**f-g**) during warm challenge (**b, d, f**) or cold challenge (**c, e, g**) respectively after LPB^Pdyn^ neural blocking (n = 6 each). (– 360 – 0 min) and (0 – 360 min) represent the averaged EE between t = (– 360 – 0 min) and t = (0 – 360 min), respectively. **h**, Changes of T_iBAT_ under different ambient temperatures (T_a_) recorded by an infrared camera (n = 4 each). The coat hair on top of the iBATs was shaved. **i**, Fever responses after injection of IL-1β (n = 10 each group). *, TeNT & IL-1β group compares to TeNT & Saline group; #, TeNT & IL-1β group compares to GFP & IL-1β group; $, GFP & IL-1β group compares to GFP & Saline group. All data are shown as mean ± s.e.m. The p-values are calculated based on ordinary two-way ANOVA with Bonferroni’s corrections (right panels in **d-g)** or repeated measure two-way ANOVA with Bonferroni’s corrections (**b, c, h, i**), or unpaired t test (right panel in **i**). *^,#,$^p ≤ 0.05; **^,$$^ p ≤ 0.01; ***^,###,$$$^ p ≤ 0.001; **** p ≤ 0.0001; ns, not significant.

**Supplemental Fig. 20.**
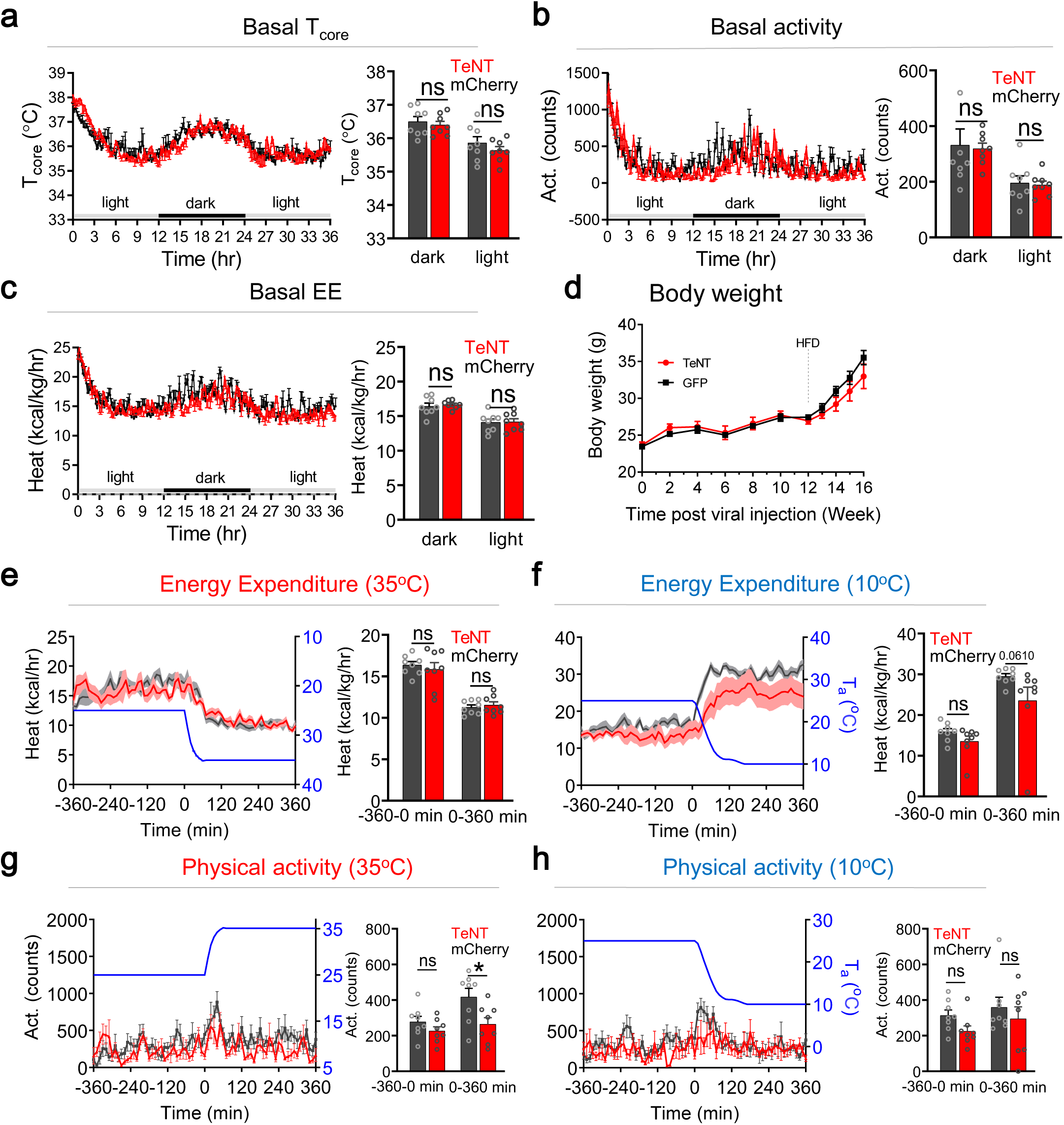
Requirement of LPB^CCK→^VMPO circuitry in heat defense and fever limiting, related to Fig. 6. **a-c**, Changes of basal T_core_ (**a**), basal physical activity (**b**), and basal energy expenditure (**c**) after blocking of POA-projected LPB^CCK^ neurons (n = 8 each; 4-6 weeks post viral injection). **d**, Changes of body weight after neural blocking under chow (0-12 weeks) and high fat diet (HFD, 12-16 weeks; n = 6 each). **e-g**, Changes of energy expenditure (**e-f**), and physical activity (**g-h**) during warm challenge (**e, g**) or cold challenge (**f, h**), respectively, after blocking of POA-projected LPB^CCK^ neurons (n = 8 each; 6-8 weeks post viral injection). (– 360 – 0 min) and (0 – 360 min) represent the averaged EE between t = (– 360 – 0 min) and t = (0 – 360 min), respectively. All data are shown as mean ± s.e.m. The p-values are calculated based on ordinary two-way ANOVA with Bonferroni’s corrections (right panels in **a-c, e-h)** or repeated measures two-way ANOVA with Bonferroni’s corrections (**d**). ^*^p ≤ 0.05; ns, not significant.

**Table S1.**
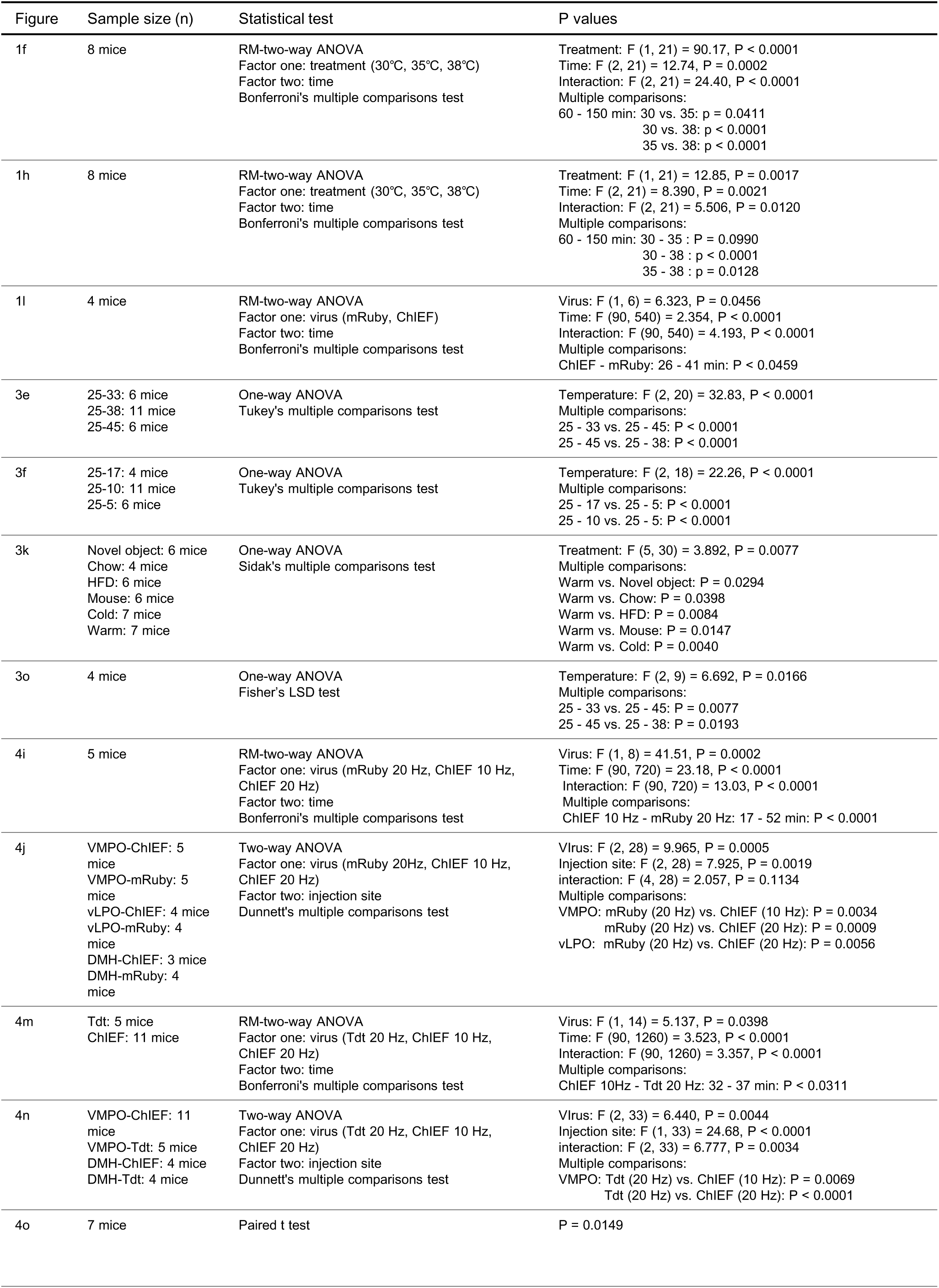
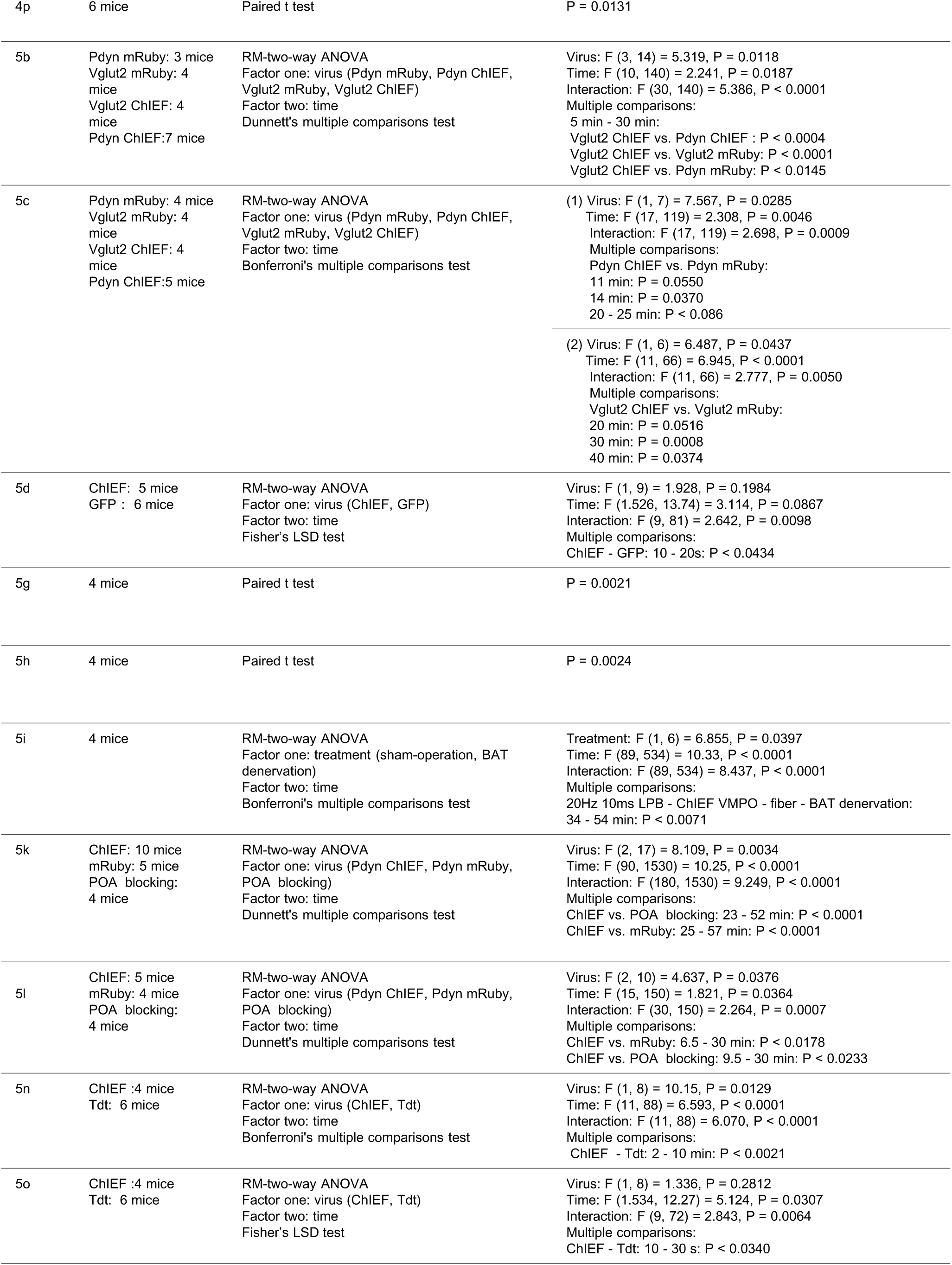
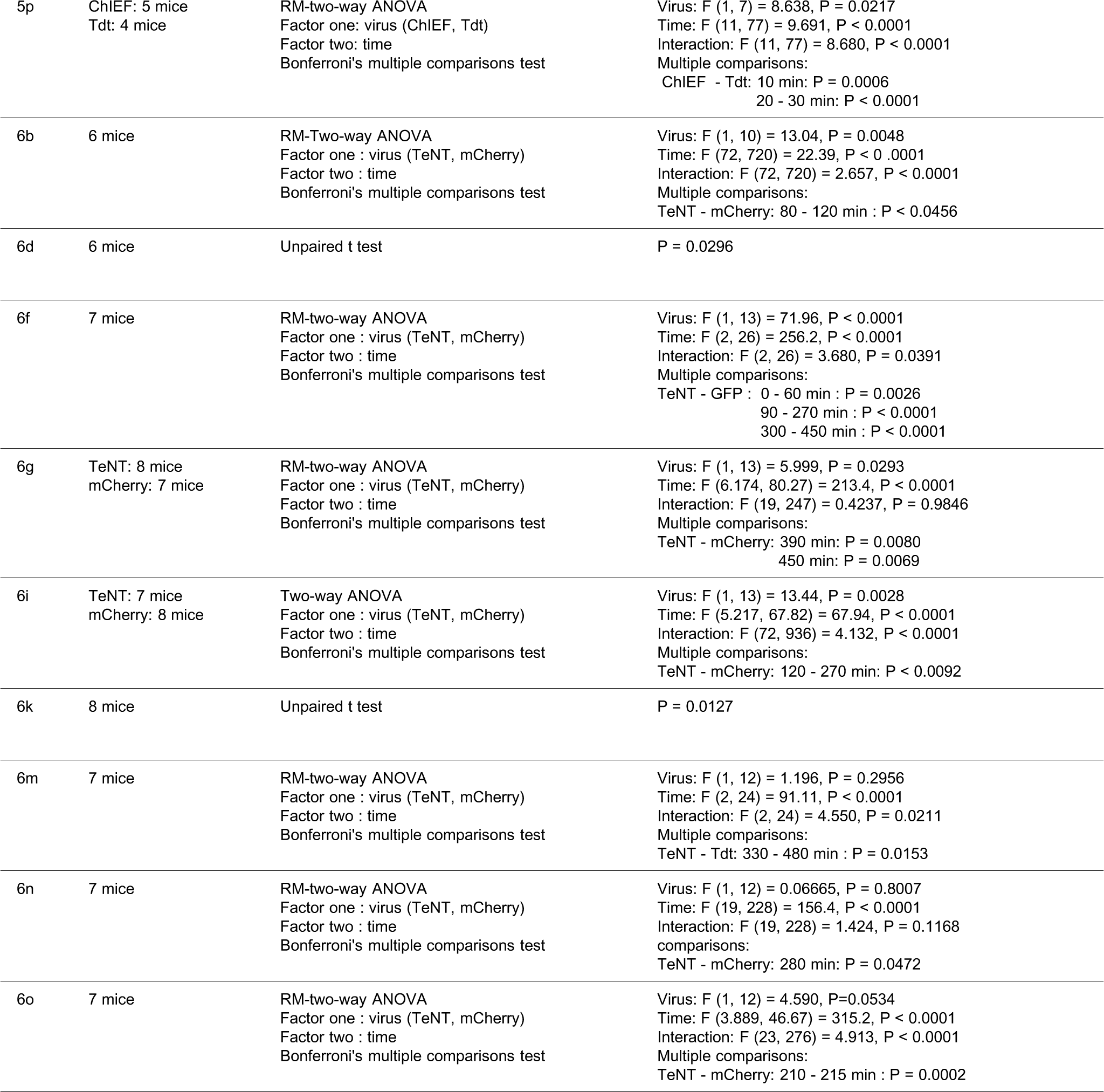
Summary of statistical analyses.

| Figure | Sample size (n) | Statistical test | P values |
| --- | --- | --- | --- |
| 1f | 8 mice | RM-two-way ANOVA<br>Factor one: treatment (30°C, 35°C, 38°C)<br>Factor two: time<br>Bonferroni's multiple comparisons test | Treatment: F (1, 21) = 90.17, P < 0.0001<br>Time: F (2, 21) = 12.74, P = 0.0002<br>Interaction: F (2, 21) = 24.40, P < 0.0001<br>Multiple comparisons:<br>60 - 150 min: 30 vs. 35: p = 0.0411<br>30 vs. 38: p < 0.0001<br>35 vs. 38: p < 0.0001 |
| 1h | 8 mice | RM-two-way ANOVA<br>Factor one: treatment (30°C, 35°C, 38°C)<br>Factor two: time<br>Bonferroni's multiple comparisons test | Treatment: F (1, 21) = 12.85, P = 0.0017<br>Time: F (2, 21) = 8.390, P = 0.0021<br>Interaction: F (2, 21) = 5.506, P = 0.0120<br>Multiple comparisons:<br>60 - 150 min: 30 - 35 : P = 0.0990<br>30 - 38 : p < 0.0001<br>35 - 38 : p = 0.0128 |
| 1l | 4 mice | RM-two-way ANOVA<br>Factor one: virus (mRuby, ChIEF)<br>Factor two: time<br>Bonferroni's multiple comparisons test | Virus: F (1, 6) = 6.323, P = 0.0456<br>Time: F (90, 540) = 2.354, P < 0.0001<br>Interaction: F (90, 540) = 4.193, P < 0.0001<br>Multiple comparisons:<br>ChIEF - mRuby: 26 - 41 min: P < 0.0459 |
| 3e | 25-33: 6 mice<br>25-38: 11 mice<br>25-45: 6 mice | One-way ANOVA<br>Tukey's multiple comparisons test | Temperature: F (2, 20) = 32.83, P < 0.0001<br>Multiple comparisons:<br>25 - 33 vs. 25 - 45: P < 0.0001<br>25 - 45 vs. 25 - 38: P < 0.0001 |
| 3f | 25-17: 4 mice<br>25-10: 11 mice<br>25-5: 6 mice | One-way ANOVA<br>Tukey's multiple comparisons test | Temperature: F (2, 18) = 22.26, P < 0.0001<br>Multiple comparisons:<br>25 - 17 vs. 25 - 5: P < 0.0001<br>25 - 10 vs. 25 - 5: P < 0.0001 |
| 3k | Novel object: 6 mice<br>Chow: 4 mice<br>HFD: 6 mice<br>Mouse: 6 mice<br>Cold: 7 mice<br>Warm: 7 mice | One-way ANOVA<br>Sidak's multiple comparisons test | Treatment: F (5, 30) = 3.892, P = 0.0077<br>Multiple comparisons:<br>Warm vs. Novel object: P = 0.0294<br>Warm vs. Chow: P = 0.0398<br>Warm vs. HFD: P = 0.0084<br>Warm vs. Mouse: P = 0.0147<br>Warm vs. Cold: P = 0.0040 |
| 3o | 4 mice | One-way ANOVA<br>Fisher's LSD test | Temperature: F (2, 9) = 6.692, P = 0.0166<br>Multiple comparisons:<br>25 - 33 vs. 25 - 45: P = 0.0077<br>25 - 45 vs. 25 - 38: P = 0.0193 |
| 4i | 5 mice | RM-two-way ANOVA<br>Factor one: virus (mRuby 20 Hz, ChIEF 10 Hz, ChIEF 20 Hz)<br>Factor two: time<br>Bonferroni's multiple comparisons test | Virus: F (1, 8) = 41.51, P = 0.0002<br>Time: F (90, 720) = 23.18, P < 0.0001<br>Interaction: F (90, 720) = 13.03, P < 0.0001<br>Multiple comparisons:<br>ChIEF 10 Hz - mRuby 20 Hz: 17 - 52 min: P < 0.0001 |
| 4j | VMPO-ChIEF: 5 mice<br>VMPO-mRuby: 5 mice<br>vLPO-ChIEF: 4 mice<br>vLPO-mRuby: 4 mice<br>DMH-ChIEF: 3 mice<br>DMH-mRuby: 4 mice | Two-way ANOVA<br>Factor one: virus (mRuby 20Hz, ChIEF 10 Hz, ChIEF 20 Hz)<br>Factor two: injection site<br>Dunnett's multiple comparisons test | Virus: F (2, 28) = 9.965, P = 0.0005<br>Injection site: F (2, 28) = 7.925, P = 0.0019<br>interaction: F (4, 28) = 2.057, P = 0.1134<br>Multiple comparisons:<br>VMPO: mRuby (20 Hz) vs. ChIEF (10 Hz): P = 0.0034<br>mRuby (20 Hz) vs. ChIEF (20 Hz): P = 0.0009<br>vLPO: mRuby (20 Hz) vs. ChIEF (20 Hz): P = 0.0056 |
| 4m | Tdt: 5 mice<br>ChIEF: 11 mice | RM-two-way ANOVA<br>Factor one: virus (Tdt 20 Hz, ChIEF 10 Hz, ChIEF 20 Hz)<br>Factor two: time<br>Bonferroni's multiple comparisons test | Virus: F (1, 14) = 5.137, P = 0.0398<br>Time: F (90, 1260) = 3.523, P < 0.0001<br>Interaction: F (90, 1260) = 3.357, P < 0.0001<br>Multiple comparisons:<br>ChIEF 10Hz - Tdt 20 Hz: 32 - 37 min: P < 0.0311 |
| 4n | VMPO-ChIEF: 11 mice<br>VMPO-Tdt: 5 mice<br>DMH-ChIEF: 4 mice<br>DMH-Tdt: 4 mice | Two-way ANOVA<br>Factor one: virus (Tdt 20 Hz, ChIEF 10 Hz, ChIEF 20 Hz)<br>Factor two: injection site<br>Dunnett's multiple comparisons test | Virus: F (2, 33) = 6.440, P = 0.0044<br>Injection site: F (1, 33) = 24.68, P < 0.0001<br>interaction: F (2, 33) = 6.777, P = 0.0034<br>Multiple comparisons:<br>VMPO: Tdt (20 Hz) vs. ChIEF (10 Hz): P = 0.0069<br>Tdt (20 Hz) vs. ChIEF (20 Hz): P < 0.0001 |
| 4o | 7 mice | Paired t test | P = 0.0149 |
| 4p | 6 mice | Paired t test | P = 0.0131 |
| 5b | Pdyn mRuby: 3 mice<br>Vglut2 mRuby: 4 mice<br>Vglut2 ChIEF: 4 mice<br>Pdyn ChIEF: 7 mice | RM-two-way ANOVA<br>Factor one: virus (Pdyn mRuby, Pdyn ChIEF, Vglut2 mRuby, Vglut2 ChIEF)<br>Factor two: time<br>Dunnett's multiple comparisons test | Virus: F (3, 14) = 5.319, P = 0.0118<br>Time: F (10, 140) = 2.241, P = 0.0187<br>Interaction: F (30, 140) = 5.386, P < 0.0001<br>Multiple comparisons:<br>5 min - 30 min:<br>Vglut2 ChIEF vs. Pdyn ChIEF : P < 0.0004<br>Vglut2 ChIEF vs. Vglut2 mRuby: P < 0.0001<br>Vglut2 ChIEF vs. Pdyn mRuby: P < 0.0145 |
| 5c | Pdyn mRuby: 4 mice<br>Vglut2 mRuby: 4 mice<br>Vglut2 ChIEF: 4 mice<br>Pdyn ChIEF: 5 mice | RM-two-way ANOVA<br>Factor one: virus (Pdyn mRuby, Pdyn ChIEF, Vglut2 mRuby, Vglut2 ChIEF)<br>Factor two: time<br>Bonferroni's multiple comparisons test | (1) Virus: F (1, 7) = 7.567, P = 0.0285<br>Time: F (17, 119) = 2.308, P = 0.0046<br>Interaction: F (17, 119) = 2.698, P = 0.0009<br>Multiple comparisons:<br>Pdyn ChIEF vs. Pdyn mRuby:<br>11 min: P = 0.0550<br>14 min: P = 0.0370<br>20 - 25 min: P < 0.086<br>(2) Virus: F (1, 6) = 6.487, P = 0.0437<br>Time: F (11, 66) = 6.945, P < 0.0001<br>Interaction: F (11, 66) = 2.777, P = 0.0050<br>Multiple comparisons:<br>Vglut2 ChIEF vs. Vglut2 mRuby:<br>20 min: P = 0.0516<br>30 min: P = 0.0008<br>40 min: P = 0.0374 |
| 5d | ChIEF: 5 mice<br>GFP : 6 mice | RM-two-way ANOVA<br>Factor one: virus (ChIEF, GFP)<br>Factor two: time<br>Fisher's LSD test | Virus: F (1, 9) = 1.928, P = 0.1984<br>Time: F (1.526, 13.74) = 3.114, P = 0.0867<br>Interaction: F (9, 81) = 2.642, P = 0.0098<br>Multiple comparisons:<br>ChIEF - GFP: 10 - 20s: P < 0.0434 |
| 5g | 4 mice | Paired t test | P = 0.0021 |
| 5h | 4 mice | Paired t test | P = 0.0024 |
| 5i | 4 mice | RM-two-way ANOVA<br>Factor one: treatment (sham-operation, BAT denervation)<br>Factor two: time<br>Bonferroni's multiple comparisons test | Treatment: F (1, 6) = 6.855, P = 0.0397<br>Time: F (89, 534) = 10.33, P < 0.0001<br>Interaction: F (89, 534) = 8.437, P < 0.0001<br>Multiple comparisons:<br>20Hz 10ms LPB - ChIEF VMPO - fiber - BAT denervation:<br>34 - 54 min: P < 0.0071 |
| 5k | ChIEF: 10 mice<br>mRuby: 5 mice<br>POA blocking: 4 mice | RM-two-way ANOVA<br>Factor one: virus (Pdyn ChIEF, Pdyn mRuby, POA blocking)<br>Factor two: time<br>Dunnett's multiple comparisons test | Virus: F (2, 17) = 8.109, P = 0.0034<br>Time: F (90, 1530) = 10.25, P < 0.0001<br>Interaction: F (180, 1530) = 9.249, P < 0.0001<br>Multiple comparisons:<br>ChIEF vs. POA blocking: 23 - 52 min: P < 0.0001<br>ChIEF vs. mRuby: 25 - 57 min: P < 0.0001 |
| 5l | ChIEF: 5 mice<br>mRuby: 4 mice<br>POA blocking: 4 mice | RM-two-way ANOVA<br>Factor one: virus (Pdyn ChIEF, Pdyn mRuby, POA blocking)<br>Factor two: time<br>Dunnett's multiple comparisons test | Virus: F (2, 10) = 4.637, P = 0.0376<br>Time: F (15, 150) = 1.821, P = 0.0364<br>Interaction: F (30, 150) = 2.264, P = 0.0007<br>Multiple comparisons:<br>ChIEF vs. mRuby: 6.5 - 30 min: P < 0.0178<br>ChIEF vs. POA blocking: 9.5 - 30 min: P < 0.0233 |
| 5n | ChIEF :4 mice<br>Tdt: 6 mice | RM-two-way ANOVA<br>Factor one: virus (ChIEF, Tdt)<br>Factor two: time<br>Bonferroni's multiple comparisons test | Virus: F (1, 8) = 10.15, P = 0.0129<br>Time: F (11, 88) = 6.593, P < 0.0001<br>Interaction: F (11, 88) = 6.070, P < 0.0001<br>Multiple comparisons:<br>ChIEF - Tdt: 2 - 10 min: P < 0.0021 |
| 5o | ChIEF :4 mice<br>Tdt: 6 mice | RM-two-way ANOVA<br>Factor one: virus (ChIEF, Tdt)<br>Factor two: time<br>Fisher's LSD test | Virus: F (1, 8) = 1.336, P = 0.2812<br>Time: F (1.534, 12.27) = 5.124, P = 0.0307<br>Interaction: F (9, 72) = 2.843, P = 0.0064<br>Multiple comparisons:<br>ChIEF - Tdt: 10 - 30 s: P < 0.0340 |
| 5p | ChIEF: 5 mice<br>Tdt: 4 mice | RM-two-way ANOVA<br>Factor one : virus (ChIEF, Tdt)<br>Factor two : time<br>Bonferroni's multiple comparisons test | Virus: F (1, 7) = 8.638, P = 0.0217<br>Time: F (11, 77) = 9.691, P < 0.0001<br>Interaction: F (11, 77) = 8.680, P < 0.0001<br>Multiple comparisons:<br>ChIEF - Tdt: 10 min: P = 0.0006<br>20 - 30 min: P < 0.0001 |
| 6b | 6 mice | RM-Two-way ANOVA<br>Factor one : virus (TeNT, mCherry)<br>Factor two : time<br>Bonferroni's multiple comparisons test | Virus: F (1, 10) = 13.04, P = 0.0048<br>Time: F (72, 720) = 22.39, P < 0.0001<br>Interaction: F (72, 720) = 2.657, P < 0.0001<br>Multiple comparisons:<br>TeNT - mCherry: 80 - 120 min : P < 0.0456 |
| 6d | 6 mice | Unpaired t test | P = 0.0296 |
| 6f | 7 mice | RM-two-way ANOVA<br>Factor one : virus (TeNT, mCherry)<br>Factor two : time<br>Bonferroni's multiple comparisons test | Virus: F (1, 13) = 71.96, P < 0.0001<br>Time: F (2, 26) = 256.2, P < 0.0001<br>Interaction: F (2, 26) = 3.680, P = 0.0391<br>Multiple comparisons:<br>TeNT - GFP : 0 - 60 min : P = 0.0026<br>90 - 270 min : P < 0.0001<br>300 - 450 min : P < 0.0001 |
| 6g | TeNT: 8 mice<br>mCherry: 7 mice | RM-two-way ANOVA<br>Factor one : virus (TeNT, mCherry)<br>Factor two : time<br>Bonferroni's multiple comparisons test | Virus: F (1, 13) = 5.999, P = 0.0293<br>Time: F (6.174, 80.27) = 213.4, P < 0.0001<br>Interaction: F (19, 247) = 0.4237, P = 0.9846<br>Multiple comparisons:<br>TeNT - mCherry: 390 min: P = 0.0080<br>450 min: P = 0.0069 |
| 6i | TeNT: 7 mice<br>mCherry: 8 mice | Two-way ANOVA<br>Factor one : virus (TeNT, mCherry)<br>Factor two : time<br>Bonferroni's multiple comparisons test | Virus: F (1, 13) = 13.44, P = 0.0028<br>Time: F (5.217, 67.82) = 67.94, P < 0.0001<br>Interaction: F (72, 936) = 4.132, P < 0.0001<br>Multiple comparisons:<br>TeNT - mCherry: 120 - 270 min: P < 0.0092 |
| 6k | 8 mice | Unpaired t test | P = 0.0127 |
| 6m | 7 mice | RM-two-way ANOVA<br>Factor one : virus (TeNT, mCherry)<br>Factor two : time<br>Bonferroni's multiple comparisons test | Virus: F (1, 12) = 1.196, P = 0.2956<br>Time: F (2, 24) = 91.11, P < 0.0001<br>Interaction: F (2, 24) = 4.550, P = 0.0211<br>Multiple comparisons:<br>TeNT - Tdt: 330 - 480 min : P = 0.0153 |
| 6n | 7 mice | RM-two-way ANOVA<br>Factor one : virus (TeNT, mCherry)<br>Factor two : time<br>Bonferroni's multiple comparisons test | Virus: F (1, 12) = 0.06665, P = 0.8007<br>Time: F (19, 228) = 156.4, P < 0.0001<br>Interaction: F (19, 228) = 1.424, P = 0.1168<br>comparisons:<br>TeNT - mCherry: 280 min: P = 0.0472 |
| 6o | 7 mice | RM-two-way ANOVA<br>Factor one : virus (TeNT, mCherry)<br>Factor two : time<br>Bonferroni's multiple comparisons test | Virus: F (1, 12) = 4.590, P=0.0534<br>Time: F (3.889, 46.67) = 315.2, P < 0.0001<br>Interaction: F (23, 276) = 4.913, P < 0.0001<br>Multiple comparisons:<br>TeNT - mCherry: 210 - 215 min : P = 0.0002 |

**Table S2.** Summary of stereotaxic coordinates.

| Brain regions |  | AP<br>(mm) | DV<br>(mm) | ML<br>(mm) | Angle (°) | Application |
| --- | --- | --- | --- | --- | --- | --- |
| LPB | P | -4.95 | -3.75 | ±1.5 | 0 (L/R) | glass pipette tip |
| LPB | C | -5.3 | -3.70 | ±1.5 | 0 (L/R) | glass pipette tip |
| VMPO |  | 0.75 | -5.0 | 0 | 0 (L/R) | glass pipette tip |
| LPB | P | -4.95 | -3.60 | ±1.5 | 0 (L/R) | fiber optic cables |
| LPB | C | -5.3 | -3.50 | ±1.5 | 0 (L/R) | fiber optic cables |
| VMPO |  | 0.75 | -4.5 | 0 | 0 (L/R) | fiber optic cables |
| vLPO |  | 0.35 | -5.3 | ±0.75 | 0(L/R) | fiber optic cables |
| DMD | a | -1.25 | -5.0 | ±0.30 | 0(L/R) | fiber optic cables |
|  | b | -1.25 | -4.8 | ±1.00 | 8(L/R) | fiber optic cables |

AP, anteroposterior; DV, dorsoventral; ML, mediolateral
P, Pdyn-Cre; C, CCK-Cre
a, b, represents different implant methods so as to co-implant fiber optic cable with POA.
L, left; R, right.

## REFERENCES

1. T. P. Yeo, Heat stroke: a comprehensive review. AACN Clin Issues 15, 280–293 (2004).

2. S. Shibolet, M. C. Lancaster, Y. Danon, Heat stroke: a review. Aviat Space Environ Med 47, 280–301 (1976).

3. F. G. Gaudio, C. K. Grissom, Cooling Methods in Heat Stroke. J Emerg Med 50, 607–616 (2016).

4. W. B. Liedtke, Deconstructing mammalian thermoregulation. Proc Natl Acad Sci U S A, (2017).

5. E. W. Freeman, K. Sherif, Prevalence of hot flushes and night sweats around the world: a systematic review. Climacteric 10, 197–214 (2007).

6. L. L. Sievert, Subjective and objective measures of hot flashes. Am J Hum Biol 25, 573–580 (2013).

7. R. R. Freedman, Menopausal hot flashes: mechanisms, endocrinology, treatment. J Steroid Biochem Mol Biol 142, 115–120 (2014).

8. C. L. Tan, Z. A. Knight, Regulation of Body Temperature by the Nervous System. Neuron 98, 31–48 (2018).

9. K. N. Shaun F. Morrison, Central neural pathways for thermoregulation. Frontiers in Bioscience 16, 74–104 (2011).

10. K. Nakamura, Central circuitries for body temperature regulation and fever. Am J Physiol-Reg I 301, R1207–R1228 (2011).

11. E. S. Bachman et al., betaAR signaling required for diet-induced thermogenesis and obesity resistance. Science 297, 843–845 (2002).

12. J. M. Friedman, J. L. Halaas, Leptin and the regulation of body weight in mammals. Nature 395, 763–770 (1998).

13. F. Wang et al., Sensory Afferents Use Different Coding Strategies for Heat and Cold. Cell Rep 23, 2001–2013 (2018).

14. D. A. Yarmolinsky et al., Coding and Plasticity in the Mammalian Thermosensory System. Neuron 92, 1079–1092 (2016).

15. C. Ran, M. A. Hoon, X. Chen, The coding of cutaneous temperature in the spinal cord. Nat Neurosci 19, 1201–1209 (2016).

16. K. Nakamura, S. F. Morrison, A thermosensory pathway mediating heat-defense responses. Proc Natl Acad Sci U S A 107, 8848–8853 (2010).

17. Y. Xue et al., In vitro thermosensitivity of rat lateral parabrachial neurons. Neurosci Lett 619, 15–20 (2016).

18. K. Nakamura, S. F. Morrison, A thermosensory pathway that controls body temperature. Nat Neurosci 11, 62–71 (2008).

19. D. Kroeger et al., Galanin neurons in the ventrolateral preoptic area promote sleep and heat loss in mice. Nat Commun 9, 4129 (2018).

20. E. C. Harding et al., A Neuronal Hub Binding Sleep Initiation and Body Cooling in Response to a Warm External Stimulus. Current Biology 28, 2263–2273 (2018).

21. Z. D. Zhao et al., A hypothalamic circuit that controls body temperature. Proc Natl Acad Sci U S A 114, 2042–2047 (2017).

22. S. B. G. Abbott, C. B. Saper, Median preoptic glutamatergic neurons promote thermoregulatory heat loss and water consumption in mice. J Physiol, (2017).

23. S. Yu et al., Glutamatergic Preoptic Area Neurons That Express Leptin Receptors Drive Temperature-Dependent Body Weight Homeostasis. Journal of Neuroscience 36, 5034–5046 (2016).

24. C. L. Tan et al., Warm-Sensitive Neurons that Control Body Temperature. Cell 167, 47–59 (2016).

25. H. W. Kun Song, Gretel B. Kamm,Jörg Pohle, Fernanda de Castro Reis, Paul Heppenstall, Hagen Wende, Jan Siemens, The TRPM2 channel is a hypothalamic heat sensor that limits fever and can drive hypothermia. Science 353, 1393–1398 (2016).

26. S. L. Padilla, C. W. Johnson, F. D. Barker, M. A. Patterson, R. D. Palmiter, A Neural Circuit Underlying the Generation of Hot Flushes. Cell Rep 24, 271–277 (2018).

27. T. A. Wang et al., Thermoregulation via Temperature-Dependent PGD2 Production in Mouse Preoptic Area. Neuron, (2019).

28. E. P. S. Conceicao, C. J. Madden, S. F. Morrison, Neurons in the rat ventral lateral preoptic area are essential for the warm-evoked inhibition of brown adipose tissue and shivering thermogenesis. Acta Physiol (Oxf), e13213 (2018).

29. T. Yahiro, N. Kataoka, Y. Nakamura, K. Nakamura, The lateral parabrachial nucleus, but not the thalamus, mediates thermosensory pathways for behavioural thermoregulation. Sci Rep 7, 5031 (2017).

30. D. Tupone, G. Cano, S. F. Morrison, Thermoregulatory inversion: a novel thermoregulatory paradigm. Am J Physiol Regul Integr Comp Physiol 312, R779–R786 (2017).

31. J. C. Geerling et al., Genetic identity of thermosensory relay neurons in the lateral parabrachial nucleus. Am J Physiol Regul Integr Comp Physiol 310, R41–54 (2016).

32. D. Engblom, M. Ek, A. Ericsson-Dahlstrand, A. Blomqvist, Activation of prostanoid EP(3) and EP(4) receptor mRNA-expressing neurons in the rat parabrachial nucleus by intravenous injection of bacterial wall lipopolysaccharide. J Comp Neurol 440, 378–386 (2001).

33. J.-F. B. Caroline Gauriau, Pain pathways and parabrachial circuits in the rat. (2001).

34. G. M. Andrade-Franze et al., Lateral parabrachial nucleus and opioid mechanisms of the central nucleus of the amygdala in the control of sodium intake. Behav Brain Res 316, 11–17 (2017).

35. D. Mu et al., A central neural circuit for itch sensation. Science 357, 695–699 (2017).

36. C. F. Roncari et al., The lateral parabrachial nucleus and central angiotensinergic mechanisms in the control of sodium intake induced by different stimuli. Behav Brain Res 333, 17–26 (2017).

37. M. E. Carter, M. E. Soden, L. S. Zweifel, R. D. Palmiter, Genetic identification of a neural circuit that suppresses appetite. Nature 503, 111–114 (2013).

38. Q. Wu, M. S. Clark, R. D. Palmiter, Deciphering a neuronal circuit that mediates appetite. Nature 483, 594–597 (2012).

39. A. L. Alhadeff, J. P. Baird, J. C. Swick, M. R. Hayes, H. J. Grill, Glucagon-like Peptide-1 receptor signaling in the lateral parabrachial nucleus contributes to the control of food intake and motivation to feed. Neuropsychopharmacology 39, 2233–2243 (2014).

40. A. L. Alhadeff, D. Golub, M. R. Hayes, H. J. Grill, Peptide YY signaling in the lateral parabrachial nucleus increases food intake through the Y1 receptor. Am J Physiol Endocrinol Metab 309, E759–766 (2015).

41. A. S. Garfield et al., A parabrachial-hypothalamic cholecystokinin neurocircuit controls counterregulatory responses to hypoglycemia. Cell Metab 20, 1030–1037 (2014).

42. J. N. Flak et al., Leptin-inhibited PBN neurons enhance responses to hypoglycemia in negative energy balance. Nat Neurosci 17, 1744–1750 (2014).

43. M. H. Qiu, M. C. Chen, P. M. Fuller, J. Lu, Stimulation of the Pontine Parabrachial Nucleus Promotes Wakefulness via Extra-thalamic Forebrain Circuit Nodes. Curr Biol 26, 2301–2312 (2016).

44. S. Kaur et al., Glutamatergic signaling from the parabrachial nucleus plays a critical role in hypercapnic arousal. J Neurosci 33, 7627–7640 (2013).

45. A. Kobayashi, T. Osaka, Involvement of the parabrachial nucleus in thermogenesis induced by environmental cooling in the rat. Pflugers Arch 446, 760–765 (2003).

46. R. Cintron-Colon et al., Activation of Kappa Opioid Receptor Regulates the Hypothermic Response to Calorie Restriction and Limits Body Weight Loss. Curr Biol 29, 4291–4299 e4294 (2019).

47. M. Schneeberger et al., Regulation of Energy Expenditure by Brainstem GABA Neurons. Cell, (2019).

48. H. Kaciuba-Uscilko, R. Grucza, Gender differences in thermoregulation. Curr Opin Clin Nutr Metab Care 4, 533–536 (2001).

49. J. Y. Lin, M. Z. Lin, P. Steinbach, R. Y. Tsien, Characterization of engineered channelrhodopsin variants with improved properties and kinetics. Biophysical Journal 96, 1803–1814 (2009).

50. B. Zingg et al., AAV-Mediated Anterograde Transsynaptic Tagging: Mapping Corticocollicular Input-Defined Neural Pathways for Defense Behaviors. Neuron 93, 33–47 (2017).

51. D. G. Tervo et al., A Designer AAV Variant Permits Efficient Retrograde Access to Projection Neurons. Neuron 92, 372–382 (2016).

52. A. R. Nectow et al., Rapid Molecular Profiling of Defined Cell Types Using Viral TRAP. Cell Rep 19, 655–667 (2017).

53. J. Vriens, B. Nilius, T. Voets, Peripheral thermosensation in mammals. Nat Rev Neurosci 15, 573–589 (2014).

54. L. Petreanu, D. Huber, A. Sobczyk, K. Svoboda, Channelrhodopsin-2-assisted circuit mapping of long-range callosal projections. Nat Neurosci 10, 663–668 (2007).

55. B. B. Holloway et al., Monosynaptic glutamatergic activation of locus coeruleus and other lower brainstem noradrenergic neurons by the C1 cells in mice. J Neurosci 33, 18792–18805 (2013).

56. N. C. Bal, S. K. Maurya, S. Singh, X. H. T. Wehrens, M. Periasamy, Increased Reliance on Muscle-based Thermogenesis upon Acute Minimization of Brown Adipose Tissue Function. Journal of Biological Chemistry 291, 17247–17257 (2016).

57. M. He et al., Strategies and Tools for Combinatorial Targeting of GABAergic Neurons in Mouse Cerebral Cortex. Neuron 91, 1228–1243 (2016).

58. P. M. Pilowsky, A. K. Goodchild, Baroreceptor reflex pathways and neurotransmitters: 10 years on. J Hypertens 20, 1675–1688 (2002).

59. R. A. Pinol et al., Brs3 neurons in the mouse dorsomedial hypothalamus regulate body temperature, energy expenditure, and heart rate, but not food intake. Nat Neurosci 21, 1530–1540 (2018).

60. J. A. Dimicco, D. V. Zaretsky, The dorsomedial hypothalamus: a new player in thermoregulation. Am J Physiol Regul Integr Comp Physiol 292, R47–63 (2007).

61. A. R. Nectow et al., Identification of a Brainstem Circuit Controlling Feeding. Cell 170, 429–442 e411 (2017).

62. S. Yu et al., Preoptic leptin signaling modulates energy balance independent of body temperature regulation. Elife 7, (2018).

63. G. Abreu-Vieira, C. Xiao, O. Gavrilova, M. L. Reitman, Integration of body temperature into the analysis of energy expenditure in the mouse. Mol Metab 4, 461–470 (2015).

64. A. Bartelt, J. Heeren, Adipose tissue browning and metabolic health. Nat Rev Endocrinol 10, 24–36 (2014).

65. K. A. Virtanen et al., Functional brown adipose tissue in healthy adults. N Engl J Med 360, 1518–1525 (2009).

66. H. N. Tang et al., Plasticity of adipose tissue in response to fasting and refeeding in male mice. Nutr Metab (Lond) 14, 3 (2017).

67. N. J. Rothwell, M. J. Stock, Effect of chronic food restriction on energy balance, thermogenic capacity, and brown-adipose-tissue activity in the rat. Biosci Rep 2, 543–549 (1982).

68. X. Yang, H. B. Ruan, Neuronal Control of Adaptive Thermogenesis. Front Endocrinol (Lausanne) 6, 149 (2015).

69. S. F. Morrison, C. J. Madden, D. Tupone, Central Neural Regulation of Brown Adipose Tissue Thermogenesis and Energy Expenditure. Cell Metab 19, 741–756 (2014).

70. H. B. Ruan et al., O-GlcNAc transferase enables AgRP neurons to suppress browning of white fat. Cell 159, 306–317 (2014).

71. R. Szymusiak, E. Satinoff, Acute thermoregulatory effects of unilateral electrolytic lesions of the medial and lateral preoptic area in rats. Physiol Behav 28, 161–170 (1982).

72. H. J. Carlisle, Effect of preoptic and anterior hypothalamic lesions on behavioral thermoregulation in the cold. J Comp Physiol Psychol 69, 391–402 (1969).

73. T. Osaka, Cold-induced thermogenesis mediated by GABA in the preoptic area of anesthetized rats. Am J Physiol Regul Integr Comp Physiol 287, R306–313 (2004).

74. K. Nakamura, S. F. Morrison, Preoptic mechanism for cold-defensive responses to skin cooling. J Physiol 586, 2611–2620 (2008).

75. M. W. Adler, E. B. Geller, C. E. Rosow, J. Cochin, The opioid system and temperature regulation. Annu Rev Pharmacol Toxicol 28, 429–449 (1988).

76. M. A. Mittelman-Smith, H. Williams, S. J. Krajewski-Hall, N. T. McMullen, N. E. Rance, Role for kisspeptin/neurokinin B/dynorphin (KNDy) neurons in cutaneous vasodilatation and the estrogen modulation of body temperature. Proc Natl Acad Sci U S A 109, 19846–19851 (2012).

77. S. Scarpellini Cda, L. H. Gargaglioni, L. G. Branco, K. C. Bicego, Role of preoptic opioid receptors in the body temperature reduction during hypoxia. Brain Res 1286, 66–74 (2009).

78. M. A. Ansonoff et al., Antinociceptive and hypothermic effects of Salvinorin A are abolished in a novel strain of kappa-opioid receptor-1 knockout mice. J Pharmacol Exp Ther 318, 641–648 (2006).

79. L. Xin, E. B. Geller, M. W. Adler, Body temperature and analgesic effects of selective mu and kappa opioid receptor agonists microdialyzed into rat brain. J Pharmacol Exp Ther 281, 499–507 (1997).

80. M. Tanaka, M. J. McKinley, R. M. McAllen, Preoptic-raphe connections for thermoregulatory vasomotor control. The Journal of neuroscience : the official journal of the Society for Neuroscience 31, 5078–5088 (2011).

81. M. Tanaka, K. Nagashima, R. M. McAllen, K. Kanosue, Role of the medullary raphe in thermoregulatory vasomotor control in rats. J Physiol 540, 657–664 (2002).

82. N. L. S. Machado et al., A Glutamatergic Hypothalamomedullary Circuit Mediates Thermogenesis, but Not Heat Conservation, during Stress-Induced Hyperthermia. Current Biology 28, 2291–2301 (2018).

83. Y. Nakamura, K. Nakamura, S. F. Morrison, Different populations of prostaglandin EP3 receptor-expressing preoptic neurons project to two fever-mediating sympathoexcitatory brain regions. Neuroscience 161, 614–620 (2009).

84. K. Yoshida, X. Li, G. Cano, M. Lazarus, C. B. Saper, Parallel preoptic pathways for thermoregulation. The Journal of neuroscience : the official journal of the Society for Neuroscience 29, 11954–11964 (2009).

85. G. T. Dodd et al., The Thermogenic Effect of Leptin Is Dependent on a Distinct Population of Prolactin-Releasing Peptide Neurons in the Dorsomedial Hypothalamus. Cell Metab 20, 639–649 (2014).

86. Y. Zhang et al., Leptin-receptor-expressing neurons in the dorsomedial hypothalamus and median preoptic area regulate sympathetic brown adipose tissue circuits. The Journal of neuroscience : the official journal of the Society for Neuroscience 31, 1873–1884 (2011).

87. C. Shang et al., Divergent midbrain circuits orchestrate escape and freezing responses to looming stimuli in mice. Nat Commun 9, 1232 (2018).

88. Z. D. Zhao et al., Zona incerta GABAergic neurons integrate prey-related sensory signals and induce an appetitive drive to promote hunting. Nat Neurosci 22, 921–932 (2019).

89. C. Shang et al., A subcortical excitatory circuit for sensory-triggered predatory hunting in mice. Nat Neurosci 22, 909–920 (2019).

90. K. Nakamura, S. F. Morrison, Central efferent pathways for cold-defensive and febrile shivering. J Physiol 589, 3641–3658 (2011).

91. X. Y. Li et al., Agrp neurons project to the medial preoptic area and modulate maternal nest-building. J Neurosci 39, 456–471 (2018).

